# Next-generation cytosine base editors with minimized unguided DNA and RNA off-target events and high on-target activity

**DOI:** 10.1101/2020.02.11.944165

**Authors:** Yi Yu, Thomas Leete, David A. Born, Lauren Young, Luis A. Barrera, Seung-Joo Lee, Holly A. Rees, Giuseppe Ciaramella, Nicole M. Gaudelli

**Affiliations:** Beam Therapeutics (Cambridge, MA, USA)

## Abstract

Cytosine base editors (CBEs) are molecular machines which enable efficient and programmable reversion of T•A to C•G point mutations in the human genome without induction of DNA double strand breaks^1, 2^. Recently, the foundational cytosine base editor (CBE) ‘BE3’, containing rAPOBEC1, was reported to induce unguided, genomic DNA^3, 4^ and cellular RNA^5^ cytosine deamination when expressed in living cells. To mitigate spurious off-target events, we developed a sensitive, high-throughput cellular assay to select next-generation CBEs that display reduced spurious deamination profiles relative to rAPOBEC1-based CBEs, whilst maintaining equivalent or superior on-target editing frequencies. We screened 153 CBEs containing cytidine deaminase enzymes with diverse sequences and identified four novel CBEs with the most promising on/off target ratios. These spurious-deamination-minimized CBEs (BE4 with either RrA3F, AmAPOBEC1, SsAPOBEC3B, or PpAPOBEC1) were further optimized for superior on- and off-target DNA editing profiles through structure-guided mutagenesis of the deaminase domain. These next-generation CBEs display comparable overall DNA on-target editing frequencies, whilst eliciting a 10- to 49-fold reduction in C-to-U edits in the transcriptome of treated cells, and up to a 33-fold overall reduction in unguided off-target DNA deamination relative to BE4 containing rAPOBEC1. Taken together, these next-generation CBEs represent a new collection of base editing tools for applications in which minimization of spurious deamination is desirable and high on-target activity is required.

---

Base editors are gene editing tools for efficient and programmable correction of pathogenic point mutations for both research and therapeutic applications^1, 6^. Unlike CRISPR-associated nuclease gene approaches, base editors do not create double-stranded DNA breaks and therefore minimize the formation of undesired editing byproducts, including insertions, deletions, translocations, and other large-scale chromosomal rearrangements^1, 2, 6, 7^. Cytosine base editors (CBEs) are comprised of a cytosine deaminase fused to an impaired form of Cas9 (D10A) which is tethered to one (BE3) or two (BE4) monomers of uracil glycosylase inhibitor (UGI)^8^. This architecture of CBEs enables the conversion of C•G base pairs to T•A base pair in human genomic DNA, through the formation of an uracil intermediate^1^.

Although CBEs lead to robust on-target DNA base editing efficiency in a variety of contexts (*e.g*. rice, wheat, human cells and bacteria, reviewed here^2^), recent reports have demonstrated that treatment of cells with high doses of BE3 can lead to low, but detectable spurious cytosine deamination in both DNA^3, 4^ and cellular RNA5 which occur in an unguided fashion, independent of the sgRNA sequence used. Specifically, in treatment of rice with BE3, “substantial” genome-wide spurious C to T SNVs occurred, above background, and enriched in genic regions^3^. Furthermore, in a study conducted by Zuo and coworkers that evaluated spurious DNA editing events resultant from microinjection of BE3 in mouse embryos, a mutation rate of one in ten million bases was detected, which resulted in approximately 300 additional single nucleotide variants (SNVs) compared to untreated cells^4^. Even though this rate of mutation is within the range that occurs naturally in mouse and human somatic cells^9, 10^, given the therapeutic importance of CBEs, we were motivated to develop next-generation CBEs that function efficiently at their on-target loci with minimal off-target spurious deamination relative to the foundational base editors, BE3/4, which contain rAPOBEC1.

Since both the reported DNA^3, 4^ and RNA^5^ off-target deamination events result from unguided, Cas9-independent deamination events, we hypothesized that these undesired editing byproducts were caused by the intrinsic ssDNA binding affinities of the cytosine deaminase itself. The canonical CBE base editor BE3, used in the aforementioned spurious deamination reports^3–5^, contains an N-terminal cytidine deaminase rAPOBEC1, an enzyme that has been extensively characterized to deaminate both DNA^11,12^ and RNA^10,13^ when expressed in mammalian, avian, and bacterial cells. Indeed, CBEs containing rAPOBEC-1 (BE3, BE4, BE4max^1,8,14^) are widely utilized base editing tools due to their overall high on-target DNA editing efficiencies, but it has not been extensively evaluated whether existing, and/or engineered deaminases might provide similar high on-target DNA editing efficiency whilst preserving a minimized unguided, deaminase dependent, off-target profile.

In order to screen a wide range of next-generation CBE candidates for preferred on- and off-target editing profiles, we established a high-throughput assay to evaluate unguided ssDNA deamination. Since spurious deamination in the genome has been reported to occur most frequently in highly transcribed regions of the genome^3,4^, it is tempting to hypothesize that rAPOBEC1 is most able to access transiently available ssDNA that is generated during DNA replication or transcription (**Fig. 1a**). Therefore, we sought to mimic the availability of genomic ssDNA by presenting this substrate *via* a secondary R-loop generated by an orthogonal SaCas9/sgRNA complex and quantified the amount of unguided editing on this ssDNA substrate with fully intact CBEs (**Fig. 1b**). Here, “*in cis*” activity refers to on-target DNA base editing and “*in trans*” activity refers to base editing in the secondary SaCas9-induced R loop, to which the base editor is not directed by its own sgRNA, mimicking the transient, unguided off-targeting editing events in the genome observed in mice^4^ and rice^3^.

**Figure 1.**
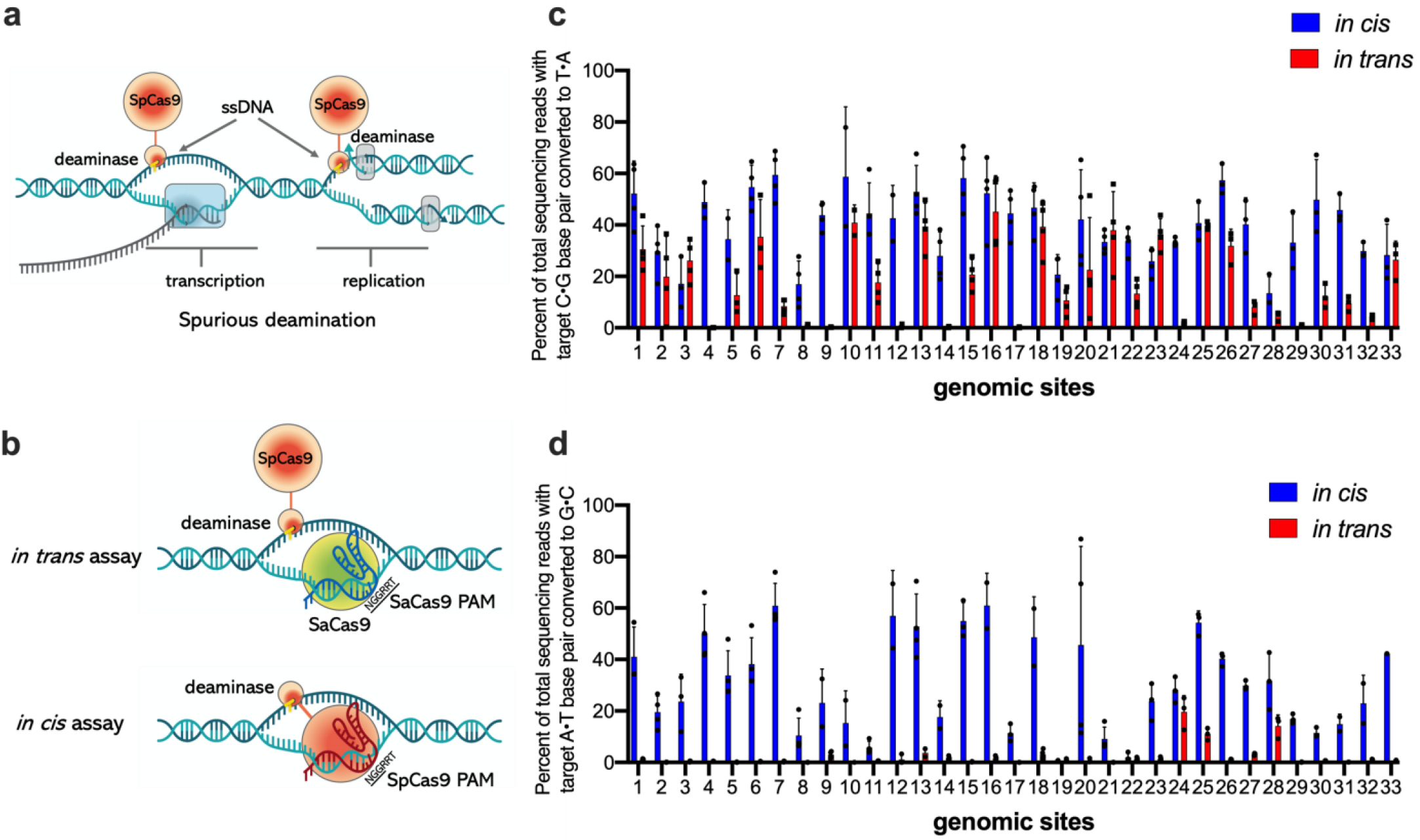
Unguided ssDNA deamination and *in cis/in trans* assay. **a**, potential ssDNA formation in the genome during transcription or replication. **b**, experimental design of *in cis/in trans* assay. Separate constructs encoding SaCas9, gRNA for SaCas9 and base editor were used to transfect HEK293T cells. *in cis* and *in trans* activity was measured in different transfections but at the target site with NGGRRT PAM sequence. **c**, *in cis/in trans* activities of BE4 with rAPOBEC1 and **d**, ABE7.10 variant at 33 genomic sites. Blue bars indicate *in cis*, on target editing, red bars indicate *in trans* editing. Base editing efficiencies were reported for the most edited base in the target sites. Values and error bars reflect the mean and s.d. of three independent biological replicates.

Crucially, we assessed the validity and sensitivity of this on- and off-target editing evaluation assay using BE4 and ABE7.10 treated cells. Previously, it has been shown that cells treated with BE3 (CBE with rAPOBEC-1), but not ABE7.10, display an increase in unguided, spurious deamination in genomic DNA^3, 4^. Consistent with these findings, our assay also showed that cells treated with BE4 (with rAPOBEC1) led to much greater levels of *in trans* editing than ABE7.10 (**Fig. 1c-d**). The sensitivity of the assay is demonstrated by the result that treatment of cells with an ABE7.10 variant led to >0.5% A-to-G editing at 20 of 33 loci tested *in trans*, up to a maximum of 25% (**Fig. 1d**). We speculate that the sensitivity of our assay may be attributed to both the presentation of the ssDNA substrate *via* stable R-loop generated by catalytically impaired Sa-Cas9 nickase with two UGI protomers attached (Sa-Cas9(D10A)-UGI-UGI)^15^ and the fact that the deamination events are measured by Illumina amplicon sequence with at least 5,000 reads per sample.

First, we used this cellular assay to test if mutagenesis of deaminases can be used to reduce *in trans* activity, which has been previously shown as a method to reduce RNA off-target editing and bystander editing^5, 16, 17^. Utilizing a homology model of rAPOBEC1 (**Supplementary Fig. 1**), based on hA3C crystal structures^18^, we identified 15 residues that we predicted to be important for ssDNA binding and 8 that affect catalytic activity. Through mutagenesis of these 23 residues, we identified 7 high-fidelity (HiFi) mutations in rAPOBEC1 (R33A, W90F, K34A, R52A, H122A, H121A, Y120F) that reduce *in trans* activity without dramatically reducing *in cis* activity (Supplementary Fig. 2). Notably, mutations of R52, H122, H121, Y120 in rAPOBEC1 have not been reported previously as methods to reduce unwanted editing activity. However, BE4 (containing rAPOBEC1) with single or double HiFi mutations led to either retention of some *in trans* activity or dramatically reduced *in cis* activity in cells (**Supplementary Fig. 3**), which prevents their broad application as a therapeutic reagent. Consequently, we began a screening campaign to survey alternative cytidine deaminases with improved editing profile in base editing context.

We began our search for next-generation CBEs with a preliminary screen of CBEs containing cytidine deaminases from well-characterized families including APOBEC1, APOBEC2, APOBEC3, APOBEC4, AID, CDA, etc. Three APOBEC1s (hAPOBEC1, PpAPOBEC1, OcAPOBEC1, MdAPOBEC1) showed a high *in cis*/*in trans* ratio at select sites (**Fig. 2a and Supplementary Fig. 4**) and, of note, primary sequence alignment of the examined APOBEBC1s with rAPOBEC1 reveal a common phenylalanine substitution at position 120 (**Supplementary Fig. 5**), a mutation identified from our structure-guided mutagenesis (Y120 in rAPOBEC1). Conversely, BE4 constructs containing deaminases which yield high *in trans* activity (rAPOBEC1, mAPOBEC1, maAPOBEC1, hA3A) all contain tyrosine at this position (**Supplementary Fig. 5**). This observation supports the predicted function of HiFi mutations and might provide explanation to the different behavior from these two groups of cytidine deaminases. BE4 variants containing PpAPOBEC1 deaminase (68% sequence identify as rAPOBEC1) showed comparable on-target DNA activity to BE4 and on average 2.3-fold decrease in *in trans* activity across 10 sites tested (**Supplementary Fig. 6**). HiFi mutations of PpAPOBEC1 were predicted based on sequence alignment (**Supplement Fig. 5**), and BE4 with PpAPOBEC1 containing either H122A or R33A mutations display desirable editing profiles (**Supplementary Fig. 6**), with 0.75x and 0.74x average *in cis* activities and 33 and 13-fold reduction in average *in trans* activities as compared to BE4 with rAPOBEC1, respectively. Together, these data make BE4 with PpAPOBEC1 the preferred CBE from our first round of screening.

Encouraged by these results, a more exhaustive screen of 43 APOBEC-like cytidine deaminases with broad sequence diversity was performed (**Fig. 2a, Supplementary Fig.6 and Supplementary Note 1**). We performed a protein BLAST with hAPOBEC1 as the query sequence to generate a sequence similarity network (SSN) with the top 1000 sequences, enabling us to select cytosine deaminases with broad sequence diversity. From this screening campaign, three constructs (BE4s with RrA3F, AmAPOBEC1, or SsAPOBEC3B) showed robust on-target DNA editing activities that are comparable to BE4 (with rAPOBEC1) with 1.05x, 0.71x, and 0.91x average *in cis* activities respectively and 2.3, 13.5, and 6.1-fold decrease on average *in trans* activity (**Fig. 2b and Supplementary Fig. 8-10**). Notably, BE4 constructs with either RrA3F or SsAPOBEC3B displayed comparably higher editing frequencies at GC target sites that are not well edited with BE4 (with rAPOBEC1) (**Supplementary Fig. 8**). Additionally, we observed variations in editing windows of *in cis* and *in trans* editing with these editors (**Supplementary Fig. 9**). Finally, we expanded our screen again to interrogate a new set of 80 putative cytidine deaminases from other protein families but none of these deaminases showed > 0.5% editing efficiency in the context of BE4 at the site tested.

**Figure 2.**
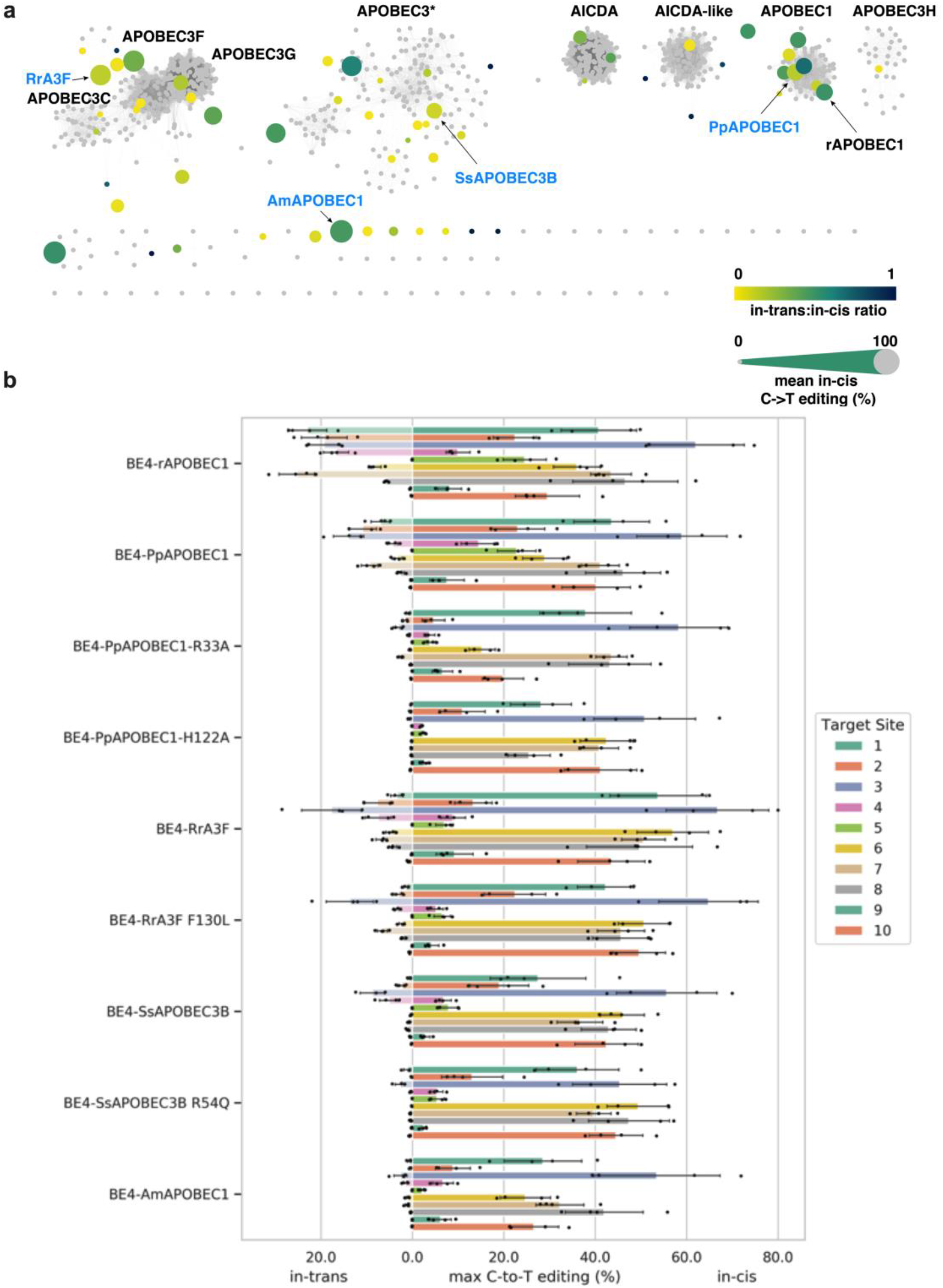
Deaminase similarity network and identified next generation CBEs with high *in cis* activities and reduced *in trans* activities compared to BE4 with rAPOBEC1. **A**, Similarity network of APOBEC-like deaminases. This network was constructed using methods described in Supplementary Note 1.The size of the dots represents the average *in cis* activity; the color of the dots represents the average *in trans/in cis* ratio. Average *in cis/in trans* activity was calculated based on three independent biological replicates on target site 1, 4, 6, and the actual editing efficiencies are displayed in Supplementary Fig.4 and Supplementary Fig.7. **b**, Comparison of *in cis* and *in trans* editing frequencies of mammalian cells treated with next generation CBEs (BE4 with PpAPOBEC1[wt, H122], RrA3F [wt, F130L], AmAPOBEC1, SsAPOBEC3B[wt, R54Q] at 10 genomic sites. Base editing efficiencies were reported for the most edited base in the target sites. Values and error bars reflect the mean and s.d. of 4 independent biological replicates.

We further optimized the BE4 editors (with RrA3F, AmAPOBEC1, or SsAPOBEC3B) by rational mutagenesis **Supplementary Fig. 2-3**). We installed rationally designed HiFi mutations from our rAPOBEC1 studies based on homology model (**Supplementary Fig. 11**) into these four BE4 editors. Two mutants (RrA3F F130L and SsAPOBEC3B R54Q) showed further improved editing profile (**Fig. 2b and Supplementary Fig. 9**), with 1.03x and 0.90x average *in cis* activities and 3.8 and 19.2-fold decrease in average to *in trans* activities, respectively, relative to BE4 containing rAPOBEC1. Together these engineered, alternative deaminase BE4 constructs offer high *in cis* with reduced *in trans* editing activity.

With these next-generation CBEs in hand we evaluated a sub-set [BE4 with PpAPOBEC1 (wt, H122A or R33A), RrA3F (wt), AmAPOBEC1 (wt), SsAPOBEC3B (wt)], to further characterize their off-target RNA activity. Plasmid-based overexpression of BE3 containing rAPOBEC1, induced “extensive transcriptome-wide RNA cytosine deamination”, and as such we evaluated our next-generation CBEs in a similar assay5. Satisfyingly, all six next-generation BE4s tested showed > 20-fold reduction in C-to-U edits as compared to BE4 with rAPOBEC1 (**Fig. 3a**). Notably, treatment of cells with BE4s containing RrA3F or SsAPOBEC3B, led to frequencies of C-to-U edits that are comparable to cells treated with nCas9 (D10A) alone. Additionally, deep-sequencing analysis of selected regions in the transcriptome reveal C-to-U editing outcomes consistent with whole transcriptome sequencing data (**Fig. 3b**). Taken together, these data suggest that the next-generation CBEs result in reduced spurious deamination in the cellular transcriptome as compared to BE3 or 4 containing rAPOBEC1.

**Figure 3.**
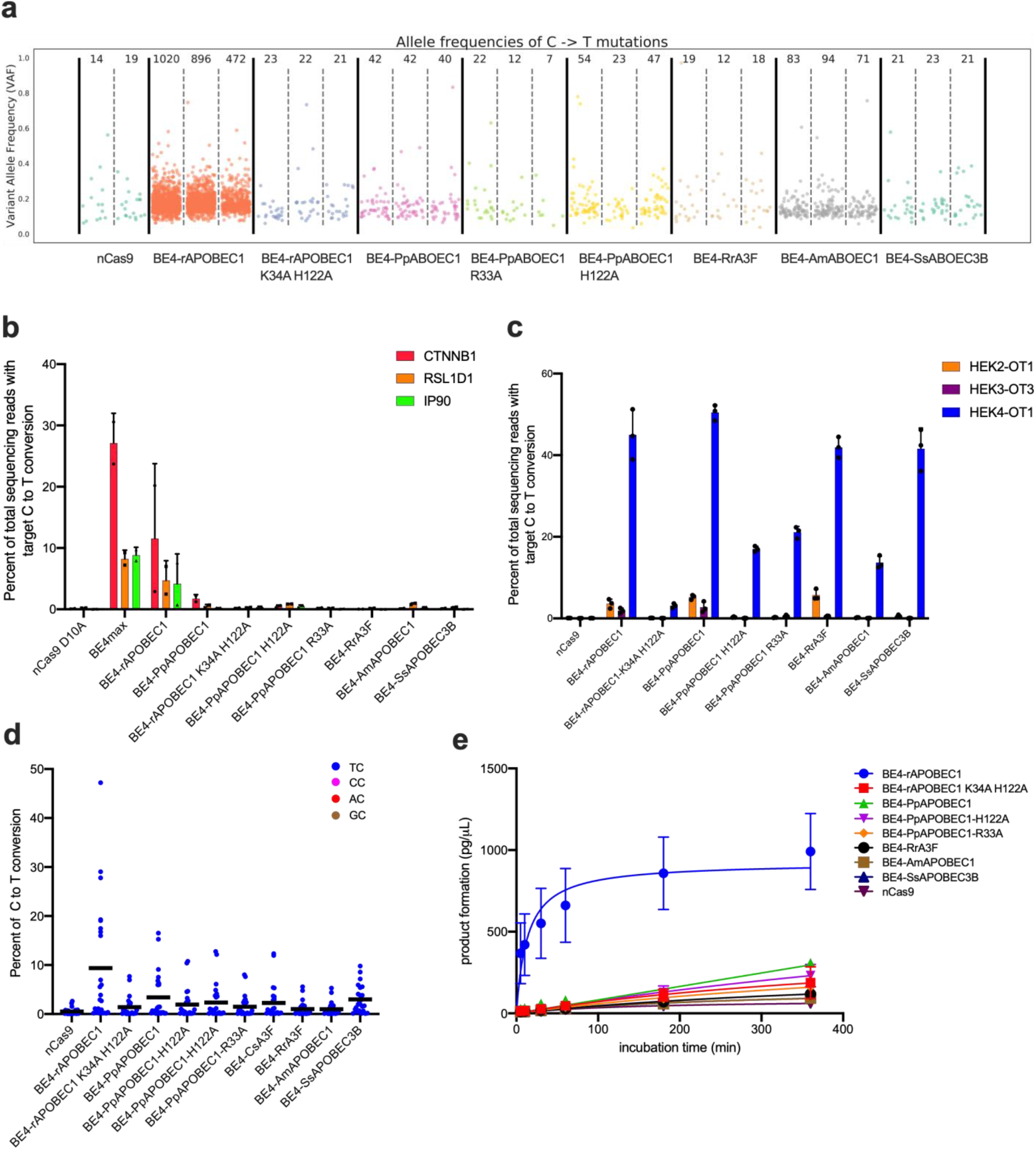
Next-generation CBEs with reduced DNA and RNA off-target editing relative to BE4 in mammalian cells. **a**, Whole transcriptome sequencing **b**, and target RNA sequencing of Hek293T cells expressing spurious deamination minimized cytosine base editors. **c**, percentage of C to T editing at know guided off-target sites. **d**, percentage of C to T editing in *in vitro* enzymatic assay on single strand DNA substrates. C to U editing of core next-generation CBEs on ssDNA substrates. Colored dots represent NC local sequence context of edit. Black line indicates average editing efficiency across target cytosines in substrates. **e**, time course of product formation in *in vitro* enzymatic assay from cell lysates containing selected CBEs. The sequences of the oligos used in d and e are listed in Supplementary Table S3. Values and error bars reflect the mean and s.d. of independent biological triplicates (**a,b,c**) or duplicates (**d, e**).

Next, we evaluated guide-dependent DNA off-target editing at known Cas9 off-target loci^19^ associated with 3 SpCas9 sgRNAs. Guide-dependent off-target activities of BE4 with PpAPOBEC1 were similar to BE4 with rAPOBEC1 (**Fig. 3c** and **Supplementary Fig. 12a-d**). Interestingly, some next-generation CBEs showed reduced guide-dependent off-target editing for at least one sgRNA tested, and our HiFi mutations also reduced guide-dependent off-target editing efficiency (**Fig. 3c and Supplementary Fig. 12a-d**). For example, at three of the most highly edited off-target sites (Hek2, site1; Hek3, site3; Hek4, site1), cells treated with BE4 containing AmAPOBEC1 engendered at least 18.8, 26.7, and 3.3-fold reduction in guide-dependent off-target editing than BE4 with rAPOBEC1 respectively (**Fig. 3c**). Notably, BE4 with PpAPOBEC1 H122A showed more than a 3-fold reduction in guide-dependent off-target editing than BE4 with PpAPOBEC1 at these three sites with no observable decrease in on-target editing (**Fig 3c**). These data indicate that next-generation CBEs can yield more favorable or equivalent guided off-target editing profiles as compared to BE4 containing rAPOBEC1. Furthermore, to validate that base editing outcomes resultant from our next-generation CBES were not due to differences in editor expression, we quantified the amount of protein produced from cells transfected with our next-generation CBEs and BE4 and show that next-generation CBE protein levels are comparable to amounts observed for BE4 (See **Supplementary Note 2**).

As a secondary evaluation of unguided DNA off-target editing, we developed an *in vitro* assay, utilizing free, synthetic ssDNA and CBE protein, as a further validation of our results obtained with the *in cis/in trans* assay described above. Total cell lysate containing base editor proteins was harvested from cells, normalized, and mixed with two synthesized oligos that contain 11 or 13 cytosines between cytosine-free adaptors, covering all NC motifs. In this assay, six next-generation CBE editors showed an average of 1.0-3.4% C-to-U editing efficiency as compared to BE4 with rAPOBEC1 which has an average of 9.4% C-to-U (data are across all 24 Cs contained within the two substrates (**Fig. 3d and Supplementary Fig. 13**). The increased ssDNA editing activity of BE4 containing rAPOBEC1, relative to our next-generation CBEs, was further supported by a time-course assay in which both the absolute level and apparent rate of deamination by BE4 with rPOABEC1 was greater than our next-generation CBEs (**Fig. 3e**). We observed 12 to 37-fold more C-to-U containing ssDNA at 5 min and 2.2 to 9.6-fold more product formed at 6 h by BE4 with rAPOBEC1 compared to our next-generation CBEs (**Fig. 3e**). Together these data suggest that we have discovered alternative, next-generation deaminases with reduced activity on exposed ssDNA, a feature that is especially important for therapeutic application of base editors.

Here, we report several new next-generation CBEs with minimized un-guided RNA and DNA off-target editing through a screening of a variety of sequence diverse cytidine deaminases. We developed two high-throughput assays to evaluate unguided ssDNA editing efficiency and from a total of 153 deaminases screened, four enzymes (PpAPOBEC1, RrA3F, AmAPOBEC1, and SsAPOBEC3B) were identified to have reduced off-target editing and high on-target editing. Together with structure-guided mutagenesis on these four constructs we highlight 8 next-generation CBEs (BE4-PpAPOBEC1, BE4-PpAPOBEC1 H122A, BE4-PpAPOBEC1 R33A, BE4-RrA3F, BE4-RrA3F F130L, BE4-AmAPOBEC1 and BE4-SsAPOBEC3B and BE4-SsAPOBEC3B R54Q) with reduced to minimized off-target editing efficiency and comparable on-target editing efficiency to BE4 containing rAPOBEC1. Transcriptome-wide RNA deamination associated with expression of these editors was comparable to that of nCas9(D10A)-2xUGI, whilst the average on-target editing was about 3.9- to 5.7-fold higher than BE4 with rAPOBEC1 with previously published SECURE mutations (R33A, K34A)^5^. Our next-generation CBEs also showed ~2 to 9-fold reduction in editing efficiency on free ssDNA oligos in *in vitro* enzymatic assay. Since the most efficient next-generation CBE can be site- and application-dependent, we recommend evaluating each of our eight next-generation CBEs for every new target of interest. However, if experimental design necessitates only one next-generation CBE be used, we advise researchers to utilize either BE4 containing PpAPOBEC1 H122A or BE4 containing RrA3F instead of the foundational BE4 with rAPOBEC1 to minimize aforementioned spurious DNA and RNA deamination events associated with rAPOBEC1. Next-generation CBEs reported here are superior replacements for the canonical BE4 and we anticipate that they will be invaluable tools for the genome editing field.

## Materials and Methods

### General Methods

Constructs used in this study were obtained by USER assembly, Gibson assembly, or synthesized by Genscript. Gene fragments used for PCR were purchased as mammalian codon-optimized gene fragments from IDT. PCR were performed with primers ordered from IDT using either Phusion U DNA Polymerase Green MultiPlex PCR Master Mix (ThermoFisher) or Q5 Hot Start High-Fidelity 2x Master Mix (New England Biolabs). Endo-free plasmids used for mammalian transfection were prepped using ZymoPURE II Plasmid Midiprep (Zymo Research Corporation) from 50 mL Mach1 (ThermoFisher) culture. Sequences for CBEs, protospacer sequences for sgRNA, and oligos used in this study can be found in **Supplementary Table 1-4**.

### HEK293T cell culture

HEK293T cells [CLBTx013, American Type Cell Culture Collection (ATCC)] were cultured in Dulbecco’s modified Eagles medium plus Glutamax (10566-016, Thermo Fisher Scientific) with 10% (v/v) fetal bovine serum (A31606-02, Thermo Fisher Scientific) following culture method on ATCC website. Cell culture incubator was set to 37 °C with 5% CO_2_. Cells were tested negative for mycoplasma after receipt from supplier.

### Transfection conditions and harvest of cells

HEK293T cells were seeded onto 96 or 48-well well Poly-D-Lysine treated BioCoat plates (Corning) at a density of 12,000 cells/well (96-well) or 35, 000 cells/well (48-well). Transfection of HEK293T cells were done after 18-24 h. To each well of cells in 96-well (48-well) plate, 90 ng (300 ng) base editor or control plasmid, 30 ng (100 ng) sgRNA plasmid and 1 μL (1.5 μL) Lipofectamine 2000 (ThermoFisher Scientific) were added following manufacturer’s instructions. For *in trans* editing experiments, 60 ng (180 ng) nSaCas9 (D10A)-2xUGI plasmid were added to the transfection mixture. After ~64 h of incubation, cells were harvested. For NGS amplicon sequencing, media was aspirated and 50-100 μL QuickExtract^™^ DNA Extraction Solution (Lucigen) were added to each well. gDNA extraction was performed according to manufacturer’s instructions; For RNA extraction (48 well-plate), 300 μL RTL plus buffer (RNasy Plus 96 kit, Qiagen) was added to each well. RIPA buffer (100 μL per well, ThermoFisher Scientific) was used to lysis the cells for protein quantification purpose. For *in vitro* enzymatic assays (48 well-plate), each well of cells were lysed with 100 μL M-per buffer (ThermoFisher Scientific).

### Next generation sequencing (NGS) and data analysis for on-target and off-target DNA editing

Genomic DNA samples were amplified and prepared for high throughput sequencing as previously reported6. Briefly, 2 μL of gDNA was added to a 25 μL PCR reaction containing Phusion U Green Multiplex PCR Master Mix and 0.5 μM of each forward and reverse primer. Following amplification, PCR products were barcoded using unique Illumina barcoding primer pairs. Barcoding reactions contained 0.5 μM of each Illumina forward and reverse primer, 1 μL of PCR mixture containing amplified genomic site of interest, and Q5 Hot Start High-Fidelity 2x Master Mix in a total volume of 25 μL. All PCR conditions were carried out as previously published. Primers used for site-specific mammalian cell genomic DNA amplification are listed in **Supplementary Table 3**.

NGS data were analyzed by performing four general steps: (1) Illumina demultiplexing, (2) read trimming and filtering, (3) alignment of all reads to the expected amplicon sequence, and (4) generation of alignment statistics and quantification of editing rates. Each step is described in more detail in **Supplementary Note 2**.

### Transcriptome sequencing for RNA off-target editing

Total RNA extraction was done using RNasy Plus 96 kit (Qiagen) following manufacturer’s protocol. An extra on-column DNase I (RNase-Free DNase Set, Qiagen) digestion step was added before the washing step following manufacturer’s instructions.

cDNA samples were generated from the isolated mRNA using SuperScript IV One-Step RT-PCR System (Thermo Fisher Scientific) according to the manufacturer’s instructions. NGS for targeted RNA sequencing was performed using the same protocol as for DNA editing. For whole transcriptome sequencing, mRNA isolation was performed from 100 ng total RNA was done using NEBNext Poly(A) mRNA Magnetic Isolation Module (NEB). Exome sequencing library preparation was performed using NEBNext^®^ Ultra^™^ II Directional RNA Library Prep Kit for Illumina following manufacturer’s instructions. The optional 2_nd_ SPRI beads selection was performed to remove residue adaptor contamination. The libraries made were analyzed using fragment analyzer (Agilent) and sequencing was conducted at Novogene on NovaSeq S4 flow cell. Data analysis was performed as described in **Supplementary Note 4**.

### *in vitro* enzymatic assays

Cells were lysed in M-per buffer and concentration of Cas9 was performed using automated Ella assay using Ella instrument (Protein Simple). An aliquot of 5 μL cell lysate or Cas9 standard solution was mixed with 45 μL sample diluent (D-13) and the mixture was added to 48-digoxigenin cartridges. Cas9 in base editor complex were quantified using anti-Cas9 antibody (7A9-A3A, Novus Biologicals). The protein concentration was adjusted to 0.1 nM (final concentration) and mixed with 1 μL oligo (oligo sequence included in **Supplementary Table 3**) at 0.5 μM concentration in reaction buffer (20 mM Tris pH 7.5, 150 mM NaCl, 1 mM DTT, 10% glycerol) for indicated amount of time. The assay was quenched by heat-inactivation at 95 °C for 3 min and the product formation was quantified using percentage of C to T conversion (NGS) and input amount of oligos.

## Acknowledgements

Bob Gantzer, Jeremy Decker, and Matt Humes are thanked for NGS support. Ian Slaymaker, Jason Gehrke, and Michael Packer are thanked for suggestions in experimental design.

## Author contributions

YY and NMG conceived of the work and designed experiments. LAB contributed to design of sgRNAs used in this study. YY and TCL conducted all experiments and performed data collection. DAB performed the generation of cytidine deaminase similarity network and deaminase selection. SYL designed mutagenesis of rAPOBEC1 and created PyMol figures. YY, DAB, LY, TCL generated all other figures. LY, DAB and LAB conducted all statistical analyses of NGS data. NMG and GC supervised the research. NMG, YY and HAR wrote the manuscript. All authors edited the manuscript.

## Data availability

Core next-generation CBEs described in this work are deposited on Addgene. High-throughput sequencing data will be deposited in the NCBI Sequence Read Archive (PRJNA595157) upon publication.

## Code accessibility

All software tools used for data analysis are publicly available. Detailed information about versions and parameters used, as well as shell commands, are provided in Supplementary Note 3 and 4.

## References

1. Komor, A.C., Kim, Y.B., Packer, M.S., Zuris, J.A. & Liu, D.R. Programmable editing of a target base in genomic DNA without double-stranded DNA cleavage. Nature 533, 420–424 (2016).

2. Rees, H.A. & Liu, D.R. Base editing: precision chemistry on the genome and transcriptome of living cells. Nat Rev Genet 19, 770–788 (2018).

3. Jin, S. et al. Cytosine, but not adenine, base editors induce genome-wide off-target mutations in rice. Science 364, 292–295 (2019).

4. Zuo, E. et al. Cytosine base editor generates substantial off-target single-nucleotide variants in mouse embryos. Science 364, 289–292 (2019).

5. Grunewald, J. et al. Transcriptome-wide off-target RNA editing induced by CRISPR-guided DNA base editors. Nature 569, 433–437 (2019).

6. Gaudelli, N.M. et al. Programmable base editing of A*T to G*C in genomic DNA without DNA cleavage. Nature 551, 464–471 (2017).

7. Webber, B.R. et al. Highly efficient multiplex human T cell engineering without double-strand breaks using Cas9 base editors. Nat Commun 10, 5222 (2019).

8. Komor, A.C. et al. Improved base excision repair inhibition and bacteriophage Mu Gam protein yields C:G-to-T:A base editors with higher efficiency and product purity. Sci Adv 3, eaao4774 (2017).

9. Milholland, B. et al. Differences between germline and somatic mutation rates in humans and mice. Nat Commun 8, 15183 (2017).

10. Lynch, M. Evolution of the mutation rate. Trends Genet 26, 345–352 (2010).

11. Saraconi, G., Severi, F., Sala, C., Mattiuz, G. & Conticello, S.G. The RNA editing enzyme APOBEC1 induces somatic mutations and a compatible mutational signature is present in esophageal adenocarcinomas. Genome Biol 15, 417 (2014).

12. Harris, R.S., Petersen-Mahrt, S.K. & Neuberger, M.S. RNA editing enzyme APOBEC1 and some of its homologs can act as DNA mutators. Mol Cell 10, 1247–1253 (2002).

13. Teng, B., Burant, C.F. & Davidson, N.O. Molecular cloning of an apolipoprotein B messenger RNA editing protein. Science 260, 1816–1819 (1993).

14. Koblan, L.W. et al. Improving cytidine and adenine base editors by expression optimization and ancestral reconstruction. Nat Biotechnol (2018).

15. Nishimasu, H. et al. Crystal Structure of Staphylococcus aureus Cas9. Cell 162, 1113–1126 (2015).

16. Grunewald, J. et al. CRISPR DNA base editors with reduced RNA off-target and self-editing activities. Nat Biotechnol 37, 1041–1048 (2019).

17. Rees, H.A., Wilson, C., Doman, J.L. & Liu, D.R. Analysis and minimization of cellular RNA editing by DNA adenine base editors. Sci Adv 5, eaax5717 (2019).

18. Shi, K. et al. Structural basis for targeted DNA cytosine deamination and mutagenesis by APOBEC3A and APOBEC3B. Nat Struct Mol Biol 24, 131–139 (2017).

19. Tsai, S. Q. et al. GUIDE-seq enables genome-wide profiling of off-target cleavage by CRISPR-Cas nucleases. Nat Biotechno 33, 187–197, doi:10.1038/nbt.3117 (2015).

