## Supplementary files for "Next-generation cytosine base editors with minimized unguided DNA and RNA off-target events and high on-target activity"

**Table of Contents**

**Supplementary Figure 1**: rAPOBEC1 homology model generated by SWISSMODEL using hAPOBEC3C structure……………………………………………………………………………………………..……4

**Supplementary Figure 2**. *in cis/in trans* editing activities of BE4 with rAPOBEC1 mutants shown in **Supplementary Figure 1**…………………………...……………………………………………………………5

**Supplementary Figure 3**: *in cis/in trans* editing activities of BE4-rAPOBEC1 with HiFi mutations at 10 target sites…………………………………………………………………………………..…………...……6

**Supplementary Figure 4**: *in cis/in trans* editing activities of CBEs tested in 1st round screening at site 1, 4, 6. … ……………………..……………………………………………………………………………………7

**Supplementary Figure 5.** Sequence alignment of CBEs tested in the 1st round screening……………..….8

**Supplementary Figure 11**: Homology models of next generation CBEs based on existing crystal structures. ……….……………………………………………………………………………………………15

**Supplementary Figure 12**: Guided off-target editing of selected next generation CBEs…………………..16

**Supplementary Figure 13**. C to T editing efficiency of selected CBEs on ssDNA substrates in *in vitro* enzymatic assay…………………………………………………………………………………………….17

**Supplementary Table 1**: Amino acid sequences of all deaminases tested in this study…….………..…..18

**Supplementary Table 2**: DNA sequences of sgRNA used in this study.…………………………...……..27

**Supplementary Table 3**: DNA sequences of oligos used in this study…………………………………….28

**Supplementary Table 4**: DNA sequences of mammalian expression plasmids for the core CBEs showed in this study.……………………………………………………………………………….……...……30

**Supplementary Note 1**: Methods of creating similarity network of cytidine deaminases............................33

**Table of Contents Continued**

**Supplementary Note 2**: Discussion about protein expression level of base editors. ………..…....…..…34

**Supplementary Figure 14**. Quantification of mRNA and protein concentration of base editors in HEK293T cells transfected base editor plasmids……………….................................................................................35

**Supplementary Note 3**: Targeted NGS analysis details…………..…..…..………………………...……...36

**Supplementary Note 4**: Transcriptome sequencing analysis methods ….…………………….……….…...38

**References**: ……………………………………………………………………………...……….…………..39

**Supplementary Figure 1.** rAPOBEC1 homology model generated by SWISSMODEL using hAPOBEC3C structure (PDB ID 3VM8). ssDNA from hAPOBEC3A structure (PDB ID 5SWW) is manually docked. **a,** mutations predicated to affect ssDNA binding. **b,** mutations predicted to affect catalytical activity.

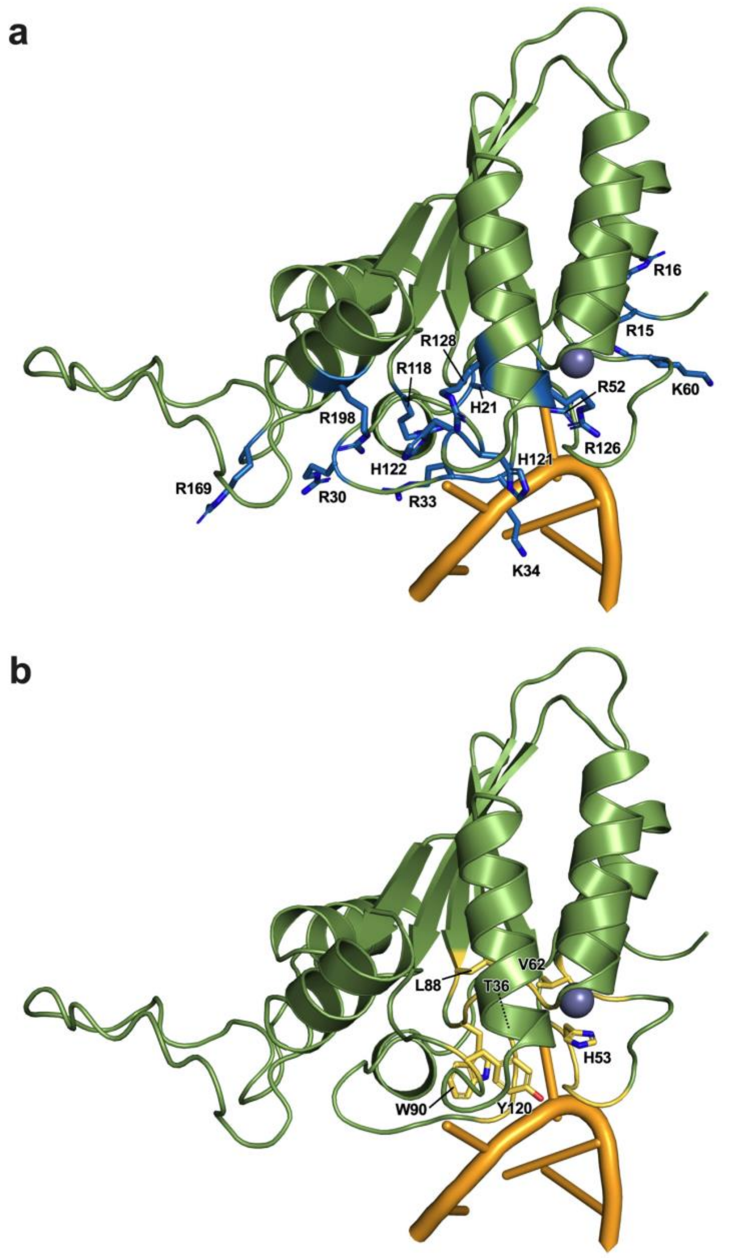

**Supplementary Figure 2.** *in cis* and *in trans* editing activities of BE4 with rAPOBEC1 mutants shown in Supplementary Figure 1 at site 1, 4, 6. Base editing efficiencies were reported for the most edited base in the target sites. Values reflect the mean of independent biological triplicates.

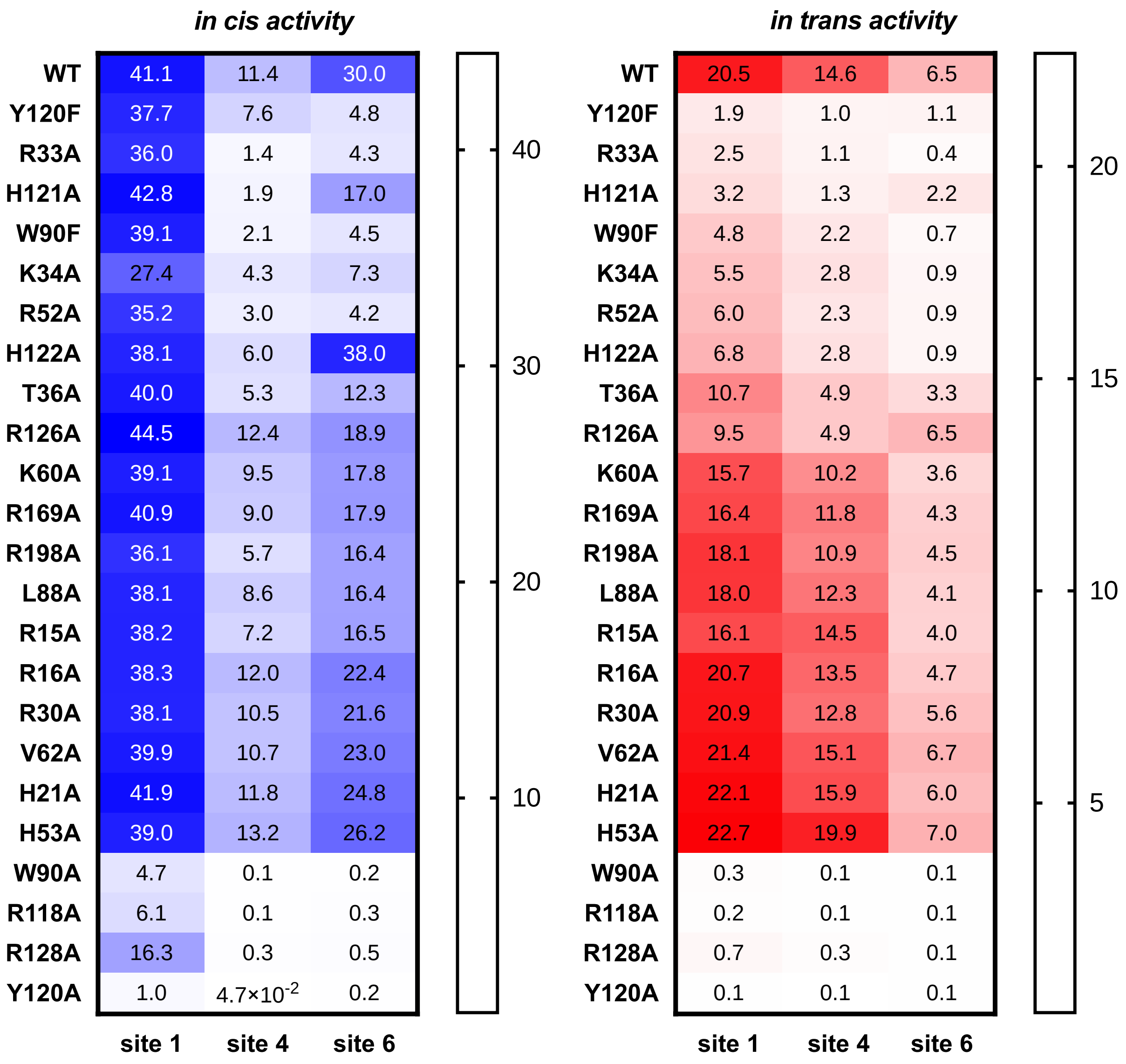

**Supplementary Figure 3.** *in cis/in trans* editing activities of BE4-rAPOBEC1 with HiFi mutations at 10 target sites. Base editing efficiencies were reported for the most edited base in the target sites. Values and error bars reflect the mean and s.d. of four independent biological replicates.

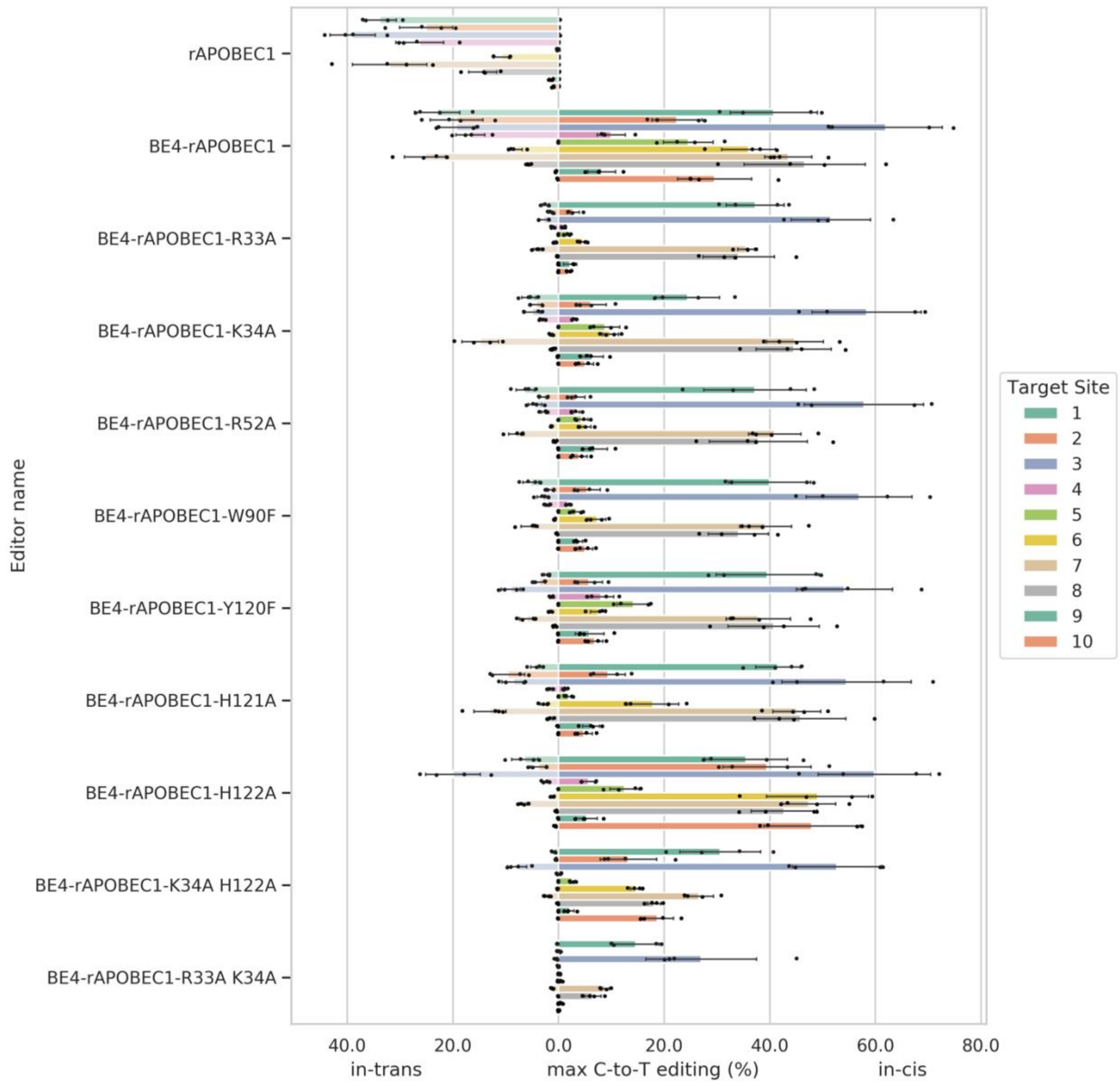

**Supplementary Figure 4.** *in cis/in trans* editing activities of CBEs tested in 1st round screening at site 1, 4, 6. Base editing efficiencies were reported for the most edited base in the target sites. Values and error bars reflect the mean and s.d. of independent biological triplicates.

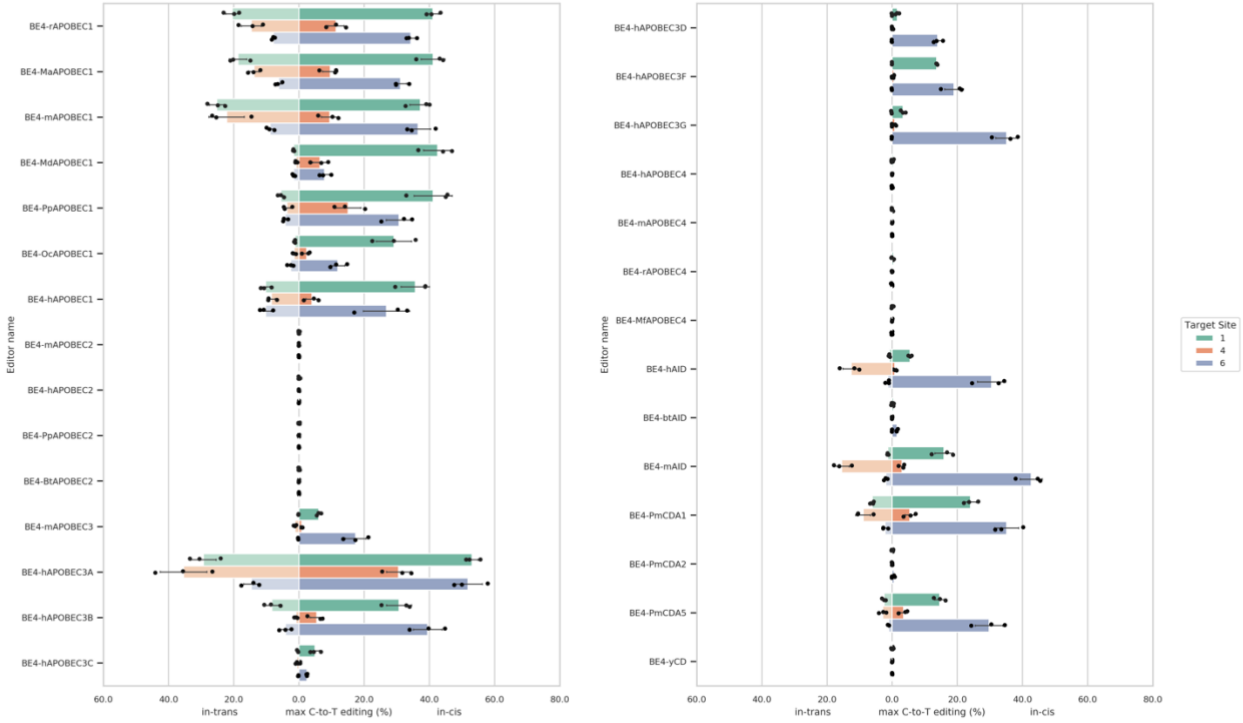

**Supplementary Figure 5.** *S*equence alignment of CBEs tested in the 1st round screening.

The amino acid residues that align to HiFi mutations in rAPOBEC1 are highlighted.

```
                  33                 52               90                        120
BE4-rAPOBEC1   -ELRKETCLLYEINWG...IWRHTSQNTNK...CSITWFLSWSPC...RYPHVTLFIYIARLYHHA-
BE4-mdAPOBEC1  -ELRKETCLLYEIKW-...IWRHSNQNTSQ...CSITWFLSWSPC...HYPNVTLAIFISRLYWHM-
BE4-maAPOBEC1  -ELRKETCLLYEIRW-...IWRHTGQNTSR...CSIVWFLSWSPC...GHPNVTLFIYAARLYHHT-
BE4-mAPOBEC1   -ELRKETCLLYEINW-...VWRHTSQNTSN...CSITWFLSWSPC...RHPYVTLFIYIARLYHHT-
BE4-ocAPOBEC1  -ELRKEACLLYEIKW-...TWRSSGKNTTN...CSITWFLSWSPC...QHPGVTLIIFVARLFQHM-
BE4-hAPOBEC1   -ELRKEACLLYEIKW-...IWRSSGKNTTN...CSITWFLSWSPC...RHPGVTLVIYVARLFWHM-
BE4-ppAPOBEC1  -ELRKETCLLYEIKW-...IWRSSGKNTTN...CSITWFLSWSPC...QHPGVTLVIYVARLFWHW-
BE4-mAPOBEC3   -YHRMKPYLCYQLEQ-...---KGCLLSEK...VTITCYLTWSPC...DRPDLILHIYTSRLYFHW-
BE4-hAPOBEC3D  -CGRNESWLCFTMEV-...FRKRGVFRNQV...YEVTWYTSWSPC...RHSNVNLTIFTARLCYFW-
BE4-hAPOBEC3F  -YGRNESWLCFTMEV-...SWKRGVFRNQV...YEVTWYTSWSPC...RHSNVNLTIFTARLYYFW-
BE4-hAPOBEC3C  -NDRNETWLCFTVEG-...SWKTGVFRNQV...YQVTWYTSWSPC...RHSNVNLTIFTARLYYFQ-
BE4-hAPOBEC3G  -RGRHETYLCYEVER-...NQRRGFLCNQA...YRVTCFTSWSPC...KNKHVSLCIFTARIYDD--
BE4-hAPOBEC3A  -IGRHKTYLCYEVER-...DQHRGFLHNQA...YRVTWFISWSPC...ENTHVRLRIFAARIYDY--
BE4-hAPOBEC3B  -LRRRQTYLCYEVER-...DQHMGFLCNEA...YRVTWFISWSPC...ENTHVRLRIFAARIYDY--
BE4-rAPOBEC4   -TYPQTKHLTFYELR-...GLASNCTGSHT...RHIILYSNNSPC...NYPEVTLSVFFSQLYHTEM
BE4-hAPOBEC4   -TFPQTKHLTFYELK-...GHASSCTGNYI...RHIILYSNNSPC...TYPGITLSIYFSQLYHTEM
BE4-mAPOBEC4   -KGRHETYLCYVVKR-...SLDFGHLRNKS...YRVTWFTSWSPC...WNPNLSLRIFTARLYFCE-
BE4-mfAPOBEC4  -TYPQTKHLTFYELK-...GHASSCTGNYI...RHIILYCNNSPC...TYPGITLSIYFSQLYHTEM
BE4-pmCDA2     -QKPRGTVILFYVEG-...AVNYNKQGTSI...CTLHCYSTYSPC...-STGVRVVIHCCRIYELDV
BE4-pmCDA1     -TERHRTYVIFDVKP-...LW--GYIINNP...YAMTWYMSWSPC...EEQGHTLTMHFSRIYDRDR
BE4-pmCDA5     -TERHRTYVIFDVKP-...LW--GYIINNP...YAMTWYMSWSPC...EEQGHTLMMHFSRLYDRDR
BE4-hAID       -KGRRETYLCYVVKR-...SLDFGYLRNKN...YRVTWFTSWSPC...GNPNLSLRIFTARLYFCE-
BE4-mAID       -KGRRETYLCYVVKR-...SLDFGYLRNKN...YRVTWFTSWSPC...GNPNLSLRIFTARLYFCE-
BE4-btAID      -KGRHETYLCYVVKR-...SLDFGHLRNKA...YRVTWFTSWSPC...GYPNLSLRIFTARLYFCDK
BE4-btAPOBEC2  -SGRNKTFLCYVVEA-...QASRGYLEDEH...YMVTWYVSSSPC...KTKNLRLLILVGRLFMWE-
BE4-mAPOBEC2   -SGRNKTFLCYVVEV-...QATQGYLEDEH...YNVTWYVSSSPC...KTKNLRLLILVSRLFMWE-
BE4-hAPOBEC2   -SGRNKTFLCYVVEA-...QASRGYLEDEH...YNVTWYVSSSPC...KTKNLRLLILVGRLFMWE-
BE4-pgAPOBEC2  -SGRNKTFLCYVVEA-...QASRGYLEDEH...YNVTWYVSSSPC...KTKNLRLLILVGRLFMWE-
BE4-yCD        -----EAALGYKEGG-...NKDGSVLGRGH...KDTTLYTTLSPC...---GIPRCVVGENVNF---
```

**Supplementary Figure 6.** *in cis/in trans* activities of BE4-PpAPOBEC1 and BE4-PpAPOBEC with HiFi mutations at 10 target sites. Base editing efficiencies were reported for the most edited base in the target sites. Values and error bars reflect the mean and s.d. of four independent biological replicates.

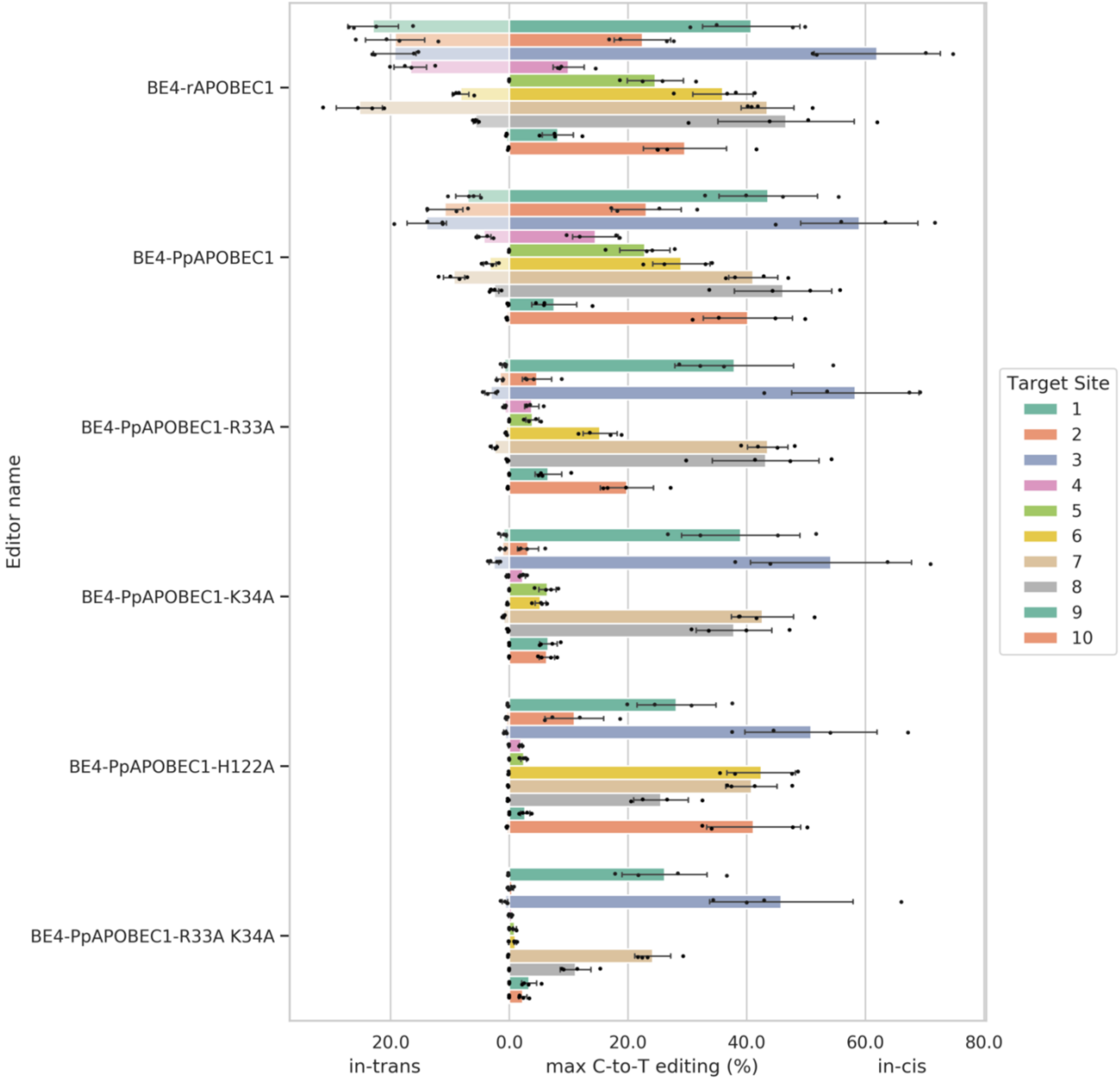

**Supplementary Figure 7.** *in cis/in trans* editing activities of CBEs tested in 2st round screening at site 1, 4, 6. Base editing efficiencies were reported for the most edited base in the target sites. Values and error bars reflect the mean and s.d. of independent biological triplicates.

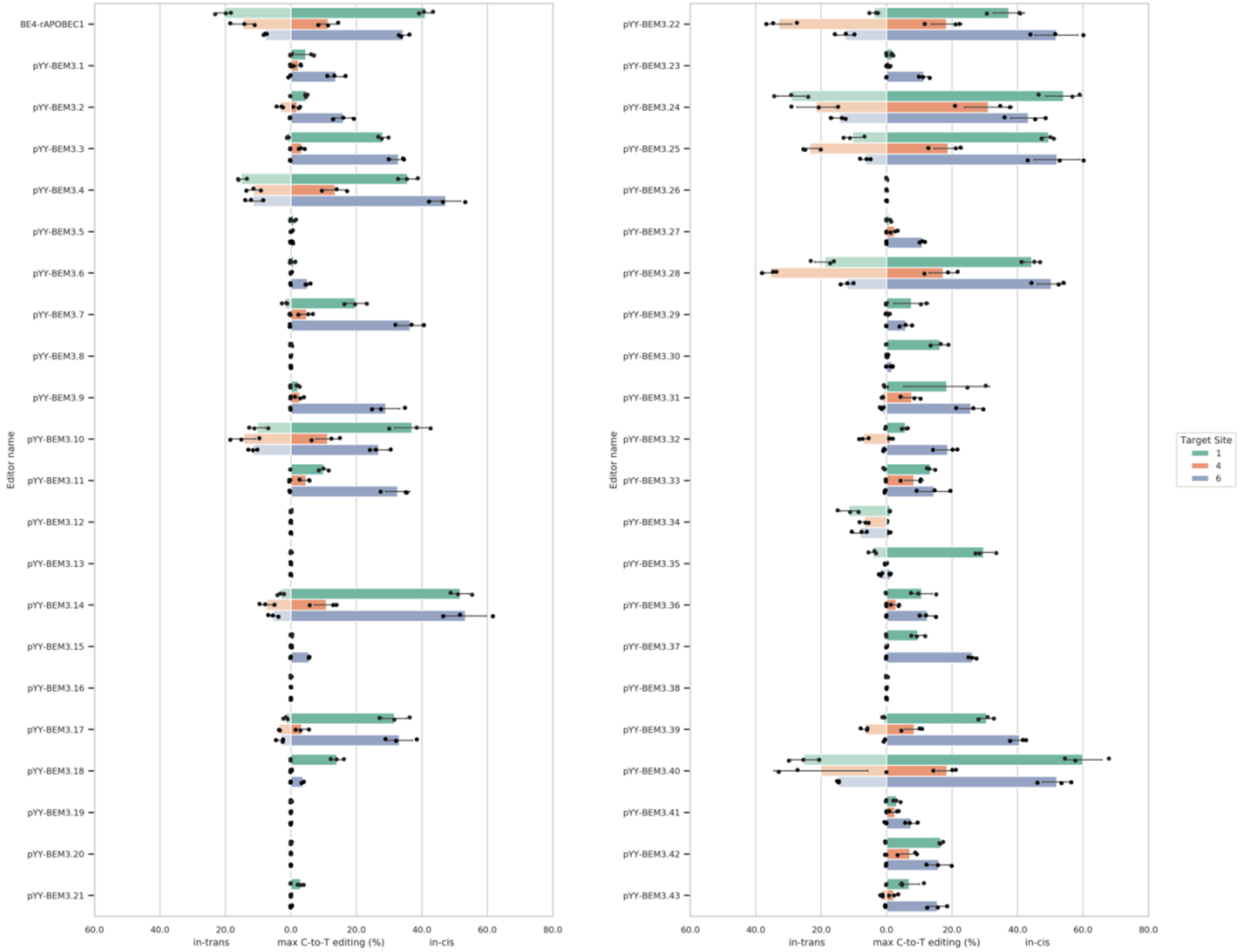

**Supplementary Figure 8.** Prior base preference of CBEs shown in Figure 2b. Values used to generate the heatmap reflect the mean of four independent biological duplicates.

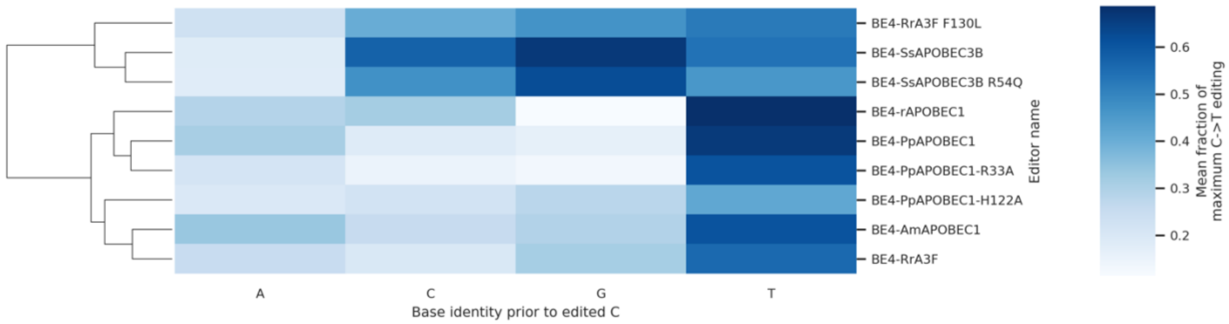

**Supplementary Figure 9.** Editing window of CBEs shown in Figure 2b at 10 target sites. Values reflect the mean of four independent biological replicates. *In cis* and *in trans* editing are presented as blue and orange heatmaps respectively.

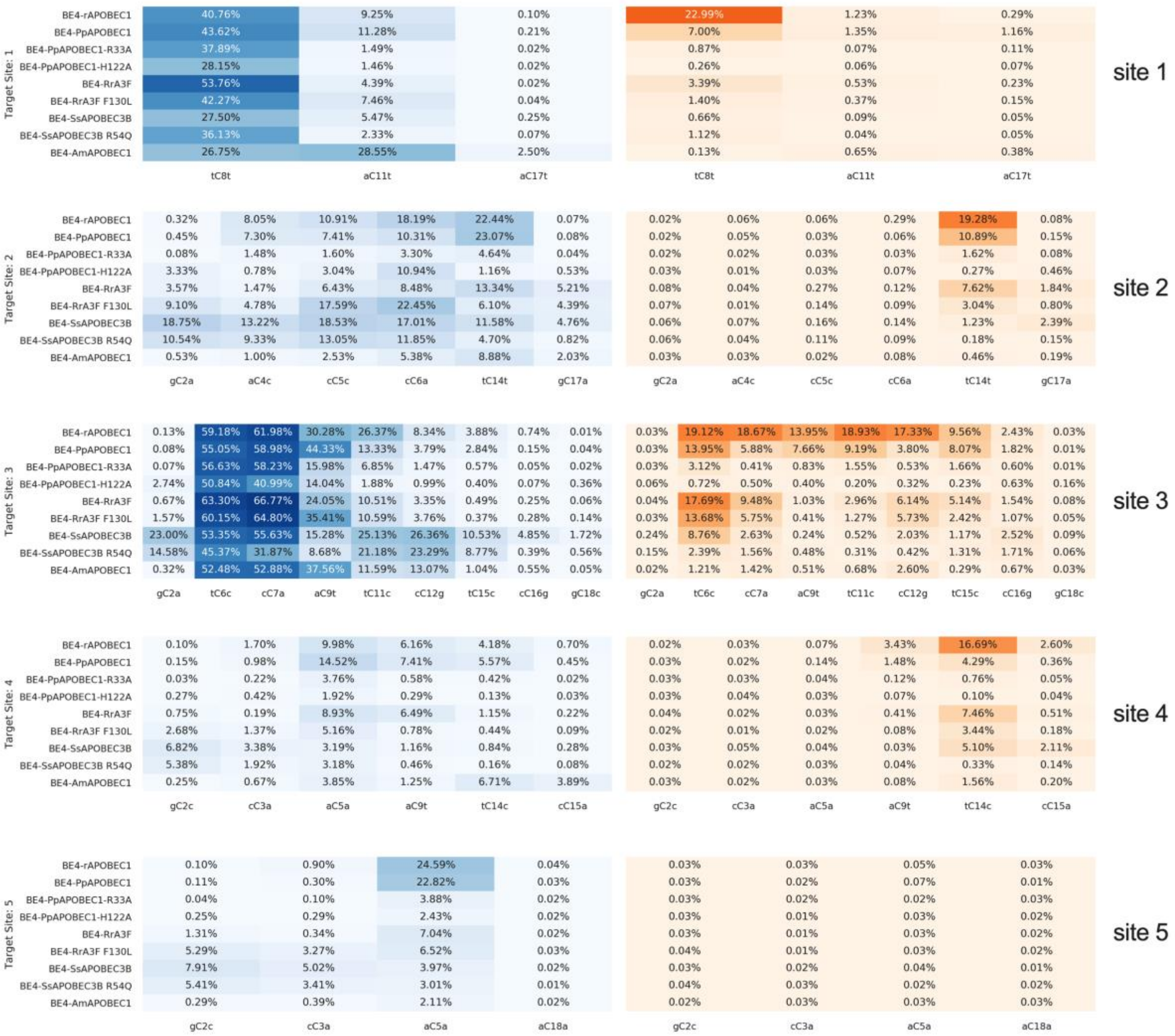

**Supplementary Figure 9 Continued.** Editing window of CBEs shown in Figure 2b at 10 target sites. Values reflect the mean of four independent biological replicates. *In cis* and *in trans* editing are presented as blue and orange heatmaps respectively.

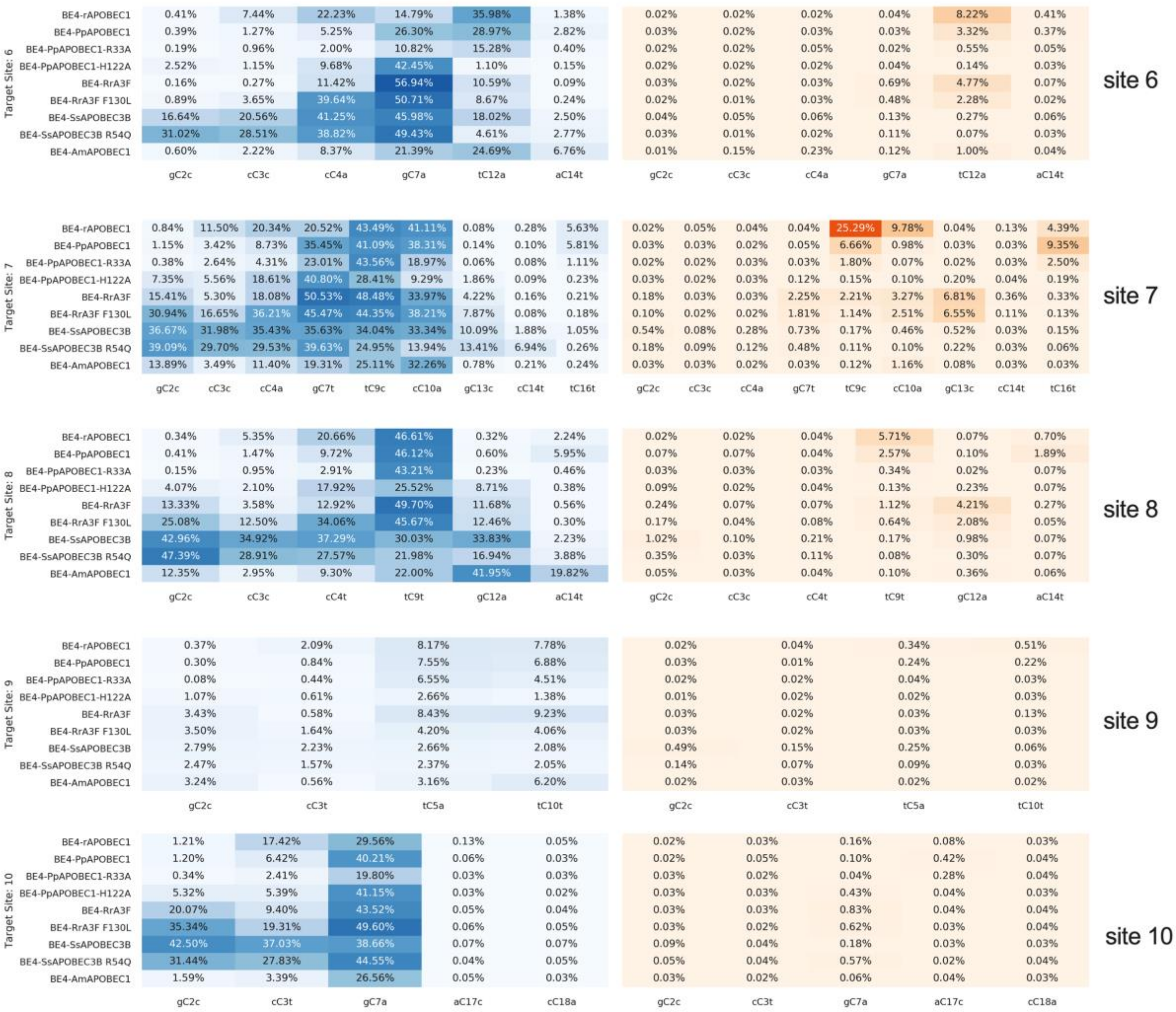

**Supplementary Figure 10.** Indel rates of CBEs shown in Figure 2b at 10 target sites. Values used to generate the heatmap reflect the mean of four independent biological duplicates.

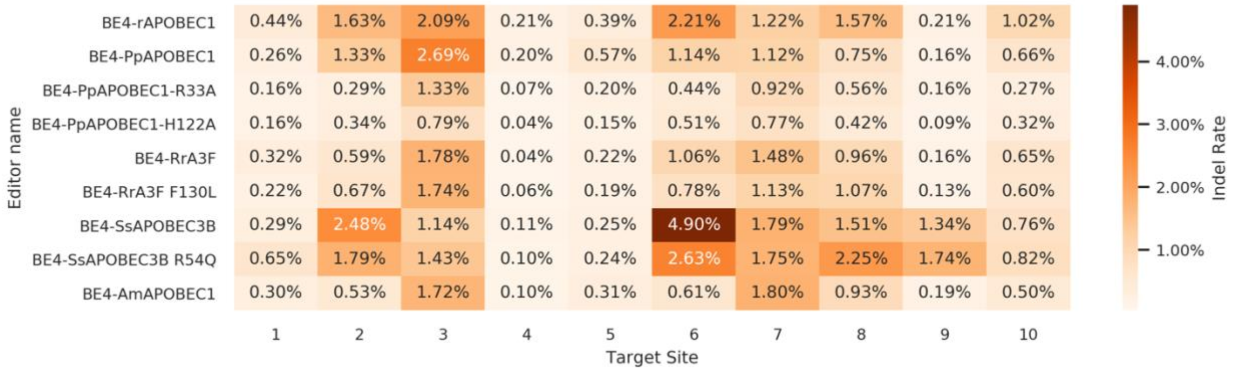

**Supplementary Figure 11.** Homology models of selected four cytidine deaminases based on existing crystal structures. **a**, Homology model of PpAPOBEC1 is based on based on a putative APOBEC3G structure (PDB ID 5K81). **b**, RrA3F is based on Vif-binding Domain of hAPOBEC3F (PDB ID 3WUS). **c**, AmAPOBEC1 is based on a hAPOBEC3B N-terminal domain (PDB ID 5TKM). **d,** SsAPOBEC3B is based on Vif-binding Domain of hAPOBEC3F (PDB ID 3WUS).

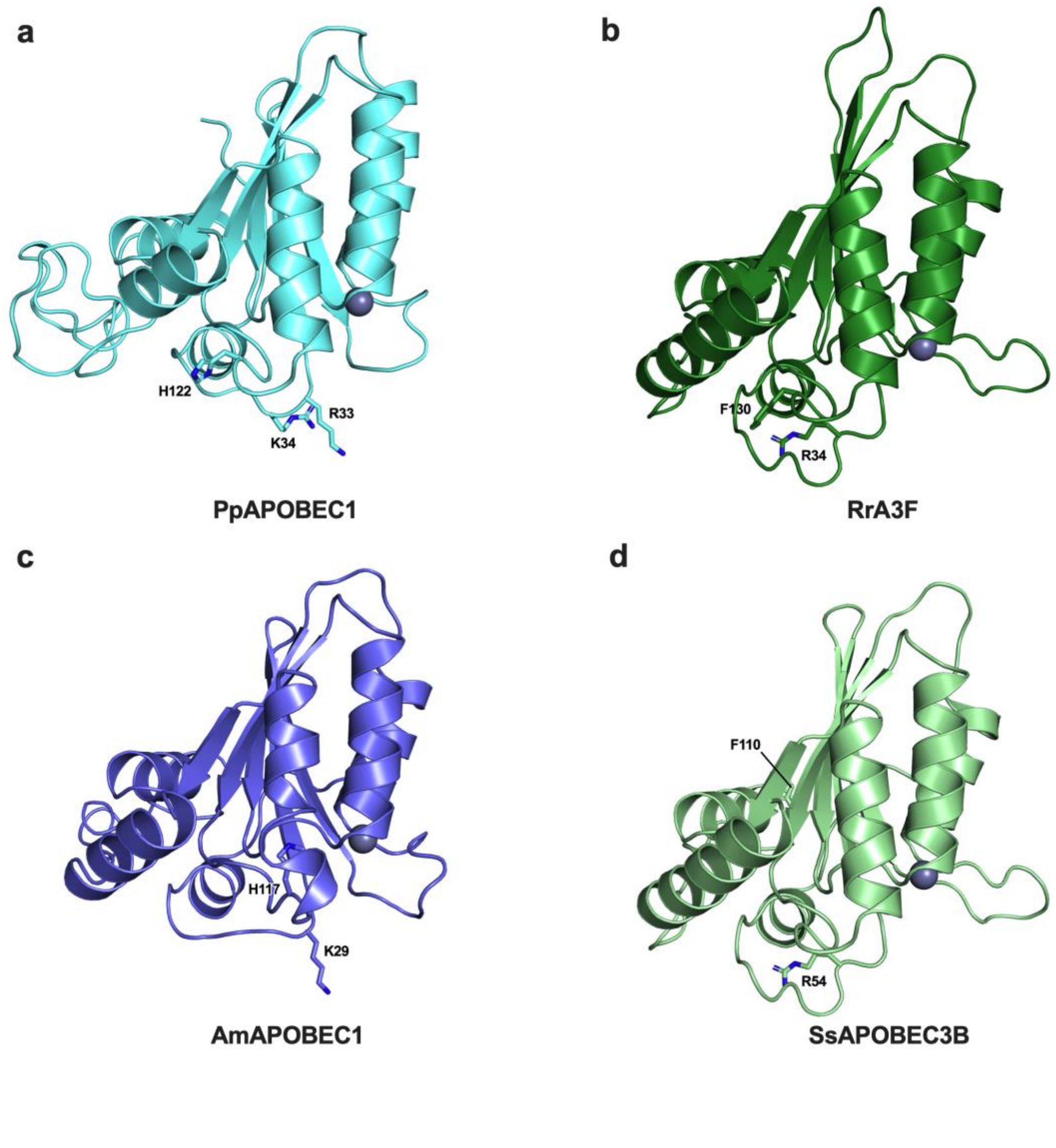

**Supplementary Figure 12.** Guided off-target editing of selected next generation CBEs. **a**, editing efficiency of next generation CBEs on HEK2, HEK3, HEK4 sites and **b**, reported guided off-target sites for HEK2 sgRNA, **c,** HEK3 sgRNA and **d,** HEK4 sgRNA. Base editing efficiencies were reported for the most edited base in the target sites. Values and error bars reflect the mean and s.d. of independent biological triplicates.

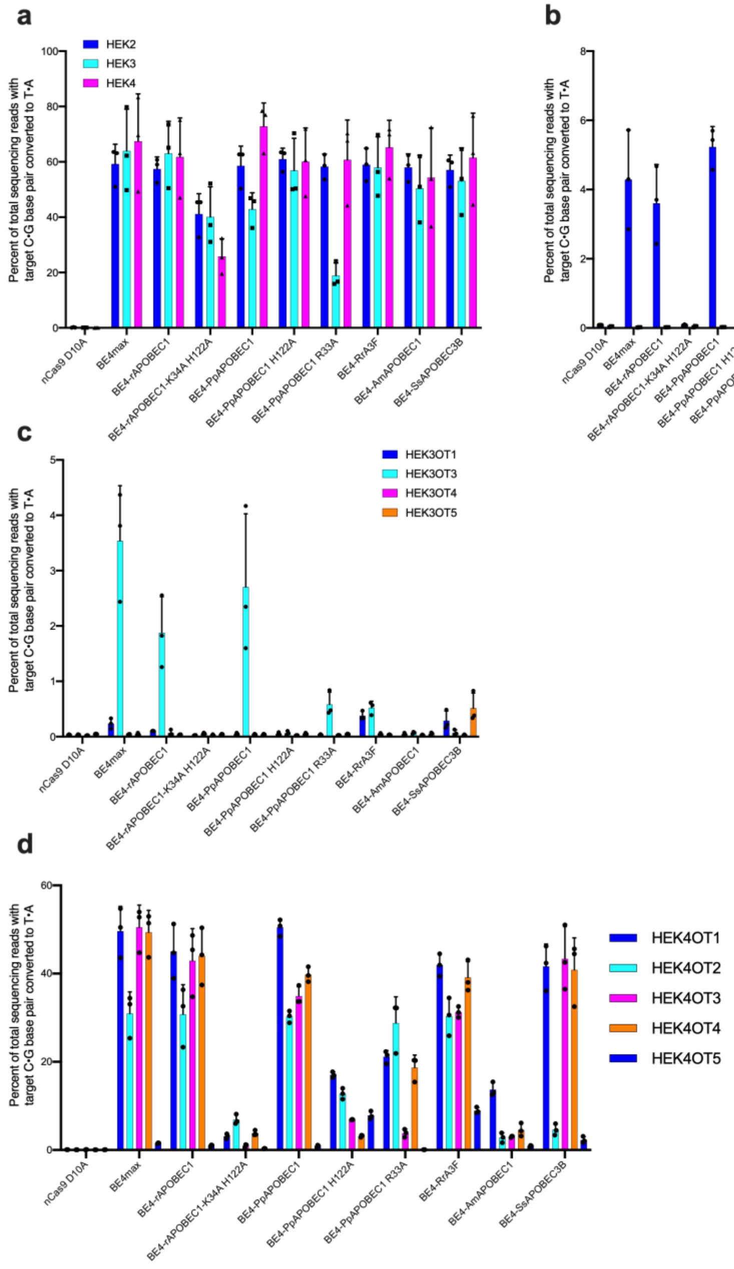

**Supplementary Figure 13.** C to T editing efficiency of selected CBEs on ssDNA substrates in *in vitro* enzymatic assay. The editing efficiencies were measured at all 25 cytidines in 2 ssDNA substrate, and group by NC sequence context. Sequences of the two substrates used are listed in Supplementary Table 2. Values and error bars reflect the mean and s.d., data were from independent biological duplicates.

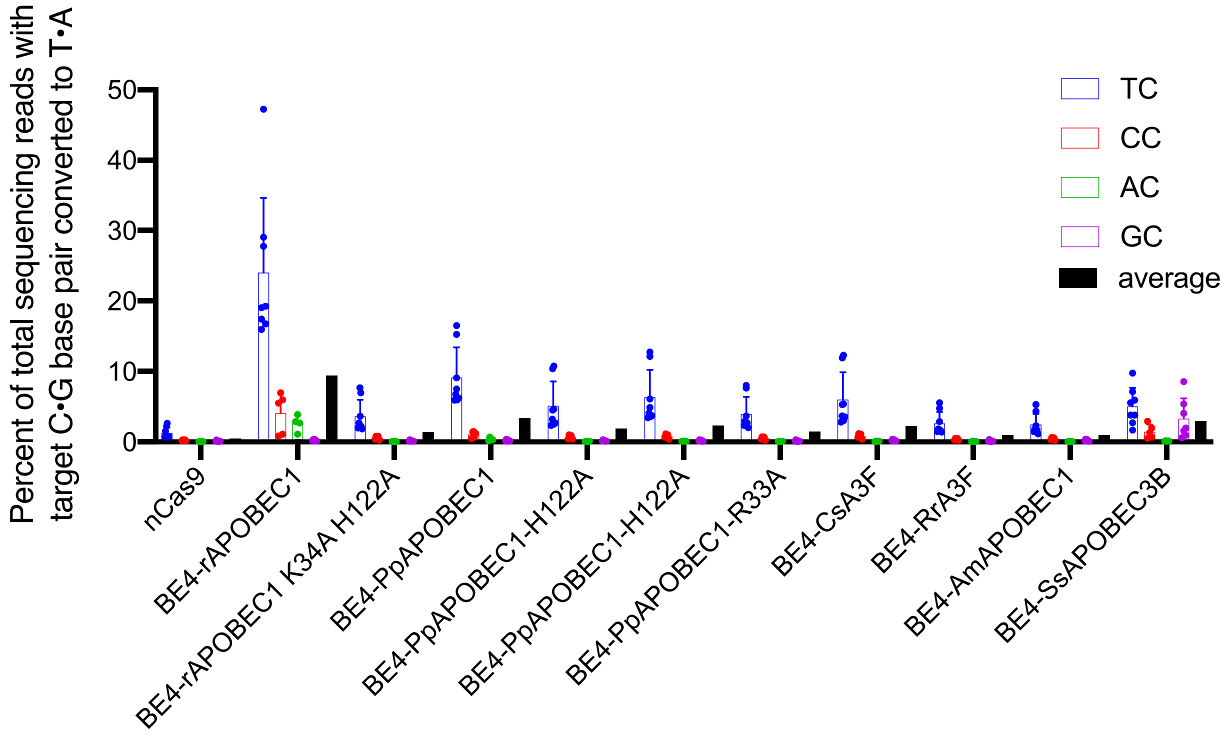

**Supplementary Table 1.** Amino acid sequences of all deaminases tested in this study.

| Gene /construct name | Species | Amino acid sequences (5'-3") |
|---|---|---|
| rAPOBEC-1 | *Rattus norvegicus* | MSSETGPVAVDPTLRRRIEPHEFEVFFDPRELRKETCLLYEINWGGRHSIWRHTSQNTNKHVEVNFIEKFTTERYFCP<br>NTRCSITWFLSWSPCGECSRAITEFLSRYPHVTLFIYIARLYHHADPRNRQGLRDLISSGVTIQIMTEQESGYCWRNFV<br>NYSPSNEAHWPRYPHLWVRLYVLELYCIILGLPPCLNILRRKQPQLTFFTIALQSCHYQRLPPHILWATGLK |
| mAPOBEC-1 | *Mus musculus* | MSSETGPVAVDPTLRRRIEPHEFEVFFDPRELRKETCLLYEINWGGRHSVWRHTSQNTSNHVEVNFLEKFTTERYFRP<br>NTRCSITWFLSWSPCGECSRAITEFLSRHPYVTLFIYIARLYHHTDQRNRQGLRDLISSGVTIQIMTEQEYCYCWRNFV<br>NYPPSNEAYWPRYPHLWVKLYVLELYCIILGLPPCLKILRRKQPQLTFFTITLQTCHYQRIPPHLLWATGLK |
| MaAPOBEC-1 | *Mesocricetus auratus* | MSSETGPVVVDPTLRRRIEPHEFDAFFDQGELRKETCLLYEIRWGGRHNIWRHTGQNTSRHVEINFIEKFTSERYFYP<br>STRCSIVWFLSWSPCGECSKAITEFLSGHPNVTLFIYAARLYHHTDQRNRQGLRDLISRGVTIRIMTEQEYCYCWRNF<br>VNYPPSNEVYWPRYPNLWMRLYALELYCIHLGLPPCLKIKRRHQYPLTFFRLNLQSCHYQRIPPHILWATGFI |
| hAPOBEC-1 | *Homo sapiens* | MTSEKGPSTGDPTLRRRIEPWEFDVFYDPRELRKEACLLYEIKWGMSRKIWRSSGKNTTNHVEVNFIKKFTSERDFHP<br>SMSCSITWFLSWSPCWECSQAIREFLSRHPGVTLVIYVARLFWHMDQQNRQGLRDLVNSGVTIQIMRASEYYHCW<br>RNFVNYPPGDEAHWPQYPPLWMMLYALELHCIILSLPPCLKISRRWQNHLTFFRLHLQNCHYQTIPPHILLATGLIHP<br>SVAWR |
| PpAPOBEC-1 | *Pongo pygmaeus* | MTSEKGPSTGDPTLRRRIESWEFDVFYDPRELRKETCLLYEIKWGMSRKIWRSSGKNTTNHVEVNFIKKFTSERRFHS<br>SISCSITWFLSWSPCWECSQAIREFLSQHPGVTLVIYVARLFWHMDQRNRQGLRDLVNSGVTIQIMRASEYYHCWR<br>NFVNYPPGDEAHWPQYPPLWMMLYALELHCIILSLPPCLKISRRWQNHLAFFRLHLQNCHYQTIPPHILLATGLIHPS<br>VTWR |
| OcAPOBEC1 | *Oryctolagus cuniculus* | MASEKGPSNKDYTLRRRIEPWEFEVFFDPQELRKEACLLYEIKWGASSKTWRSSGKNTTNHVEVNFLEKLTSEGRLG<br>PSTCCSITWFLSWSPCWECSMAIREFLSQHPGVTLIIFVARLFQHMDRRNRQGLKDLVTSGVTVRVMSVSEYCYCW<br>ENFVNYPPGKAAQWPRYPPRWMLMYALELYCIILGLPPCLKISRRHQKQLTFFSLTPQYCHYKMIPPYILLATGLLQPS<br>VPWR |
| MdAPOBEC-1 | *Monodelphis domestica* | MNSKTGPSVGDATLRRRIKPWEFVAFFNPQELRKETCLLYEIKWGNQNIWRHSNQNTSQHAEINFMEKFTAERHF<br>NSSVRCSITWFLSWSPCWECSKAIRKFLDHYPNVTLAIFISRLYWHMDQQHRQGLKELVHSGVTIQIMSYSEYHYCW<br>RNFVDYPQGEEDYWPKYPYLWIMLYVLELHCIILGLPPCLKISGSHSNQLALFSLDLQDCHYQKIPYNVLVATGLVQPF<br>VTWR |
| mAPOBEC-2 | *Mus musculus* | MAQKEEAAEAAAPASQNGDDLENLEDPEKLKELIDLPPFEIVTGVRLPVNFFKFQFRNVEYSSGRNKTFLCYVVEVQS<br>KGGQAQATQGYLEDEHAGAHAEEAFFNTILPAFDPALKYNVTWYVSSSPCAACADRILKTLSKTKNLRLLILVSRLFM<br>WEEPEVQAALKKLKEAGCKLRIMKPQDFEYIWQNFVEQEEGESKAFEPWEDIQENFLYYEEKLADILK |
| hAPOBEC-2 | *Homo sapiens* | MAQKEEAAVATEAASQNGEDLENLDDPEKLKELIELPPFEIVTGERLPANFFKFQFRNVEYSSGRNKTFLCYVVEAQG<br>KGGQVQASRGYLEDEHAAAHAEEAFFNTILPAFDPALRYNVTWYVSSSPCAACADRIIKTLSKTKNLRLLILVGRLFM<br>WEEPEIQAALKKLKEAGCKLRIMKPQDFEYVWQNFVEQEEGESKAFQPWEDIQENFLYYEEKLADILK |
| PpAPOBEC-2 | *Pongo pygmaeus* | MAQKEEAAAATEAASQNGEDLENLDDPEKLKELIELPPFEIVTGERLPANFFKFQFRNVEYSSGRNKTFLCYVVEAQG<br>KGGQVQASRGYLEDEHAAAHAEEAFFNTILPAFDPALRYNVTWYVSSSPCAACADRIIKTLSKTKNLRLLILVGRLFM<br>WEELEIQDALKKLKEAGCKLRIMKPQDFEYVWQNFVEQEEGESKAFQPWEDIQENFLYYEEKLADILK |
| BtAPOBEC-2 | *Bos taurus* | MAQKEEAAAAAEPASQNGEEVENLEDPEKLKELIELPPFEIVTGERLPAHYFKFQFRNVEYSSGRNKTFLCYVVEAQS<br>KGGQVQASRGYLEDEHATNHAEEAFFNSIMPTFDPALRYMVTWYVSSSPCAACADRIVKTLNKTKNLRLLILVGRLF<br>MWEEPEIQAALRKLKEAGCRLRIMKPQDFEYIWQNFVEQEEGESKAFEPWEDIQENFLYYEEKLADILK |
| mAPOBEC-3 | *Mus musculus* | MQPQRLGPRAGMGPFCLGCSHRKCYSPIRNLISQETFKFHFKNLGYAKGRKDTFLCYEVTRKDCDSPVSLHHGVFKN<br>KDNIHAEICFLYWFHDKVLKVLSPREEFKITWYMSWSPCFECAEQIVRFLATHHNLSLDIFSSRLYNVQDPETQQNLC<br>RLVQEGAQVAAMDLYEFKKCWKKFVDNGGRRFRPWKRLLTNFRYQDSKLQEILRPCYISVPSSSSSTLSNICLTKGLP<br>ETRFWVEGRRMDPLSEEEFYSQFYNQRVKHLCYYHRMKPYLCYQLEQFNGQAPLKGCLLSEKGKQHAEILFLDKIRS<br>MELSQVTITCYLTWSPCPNCAWQLAAFKRDRPDLILHIYTSRLYFHWKRPFQKGLCSLWQSGILVDVMDLPQFTDC<br>WTNFVNPKRPFWPWKGLEIISRRTQRRLRRIKESWGLQDLVNDFGNLQLGPPMS |
| hAPOBEC-3A | *Homo sapiens* | MEASPASGPRHLMDPHIFTSNFNNGIGRHKTYLCYEVERLDNGTSVKMDQHRGFLHNQAKNLLCGFYGRHAELRF<br>LDLVPSLQLDPAQIYRVTWFISWSPCFSWGCAGEVRAFLQENTHVRLRIFAARIYDYDPLYKEALQMLRDAGAQVSI<br>MTYDEFKHCWDTFVDHQGCPFQPWDGLDEHSQALSGRLRAILQNQGN |
| hAPOBEC-3B | *Homo sapiens* | MNPQIRNPMERMYRDTFYDNFENEPILYGRSYTWLCYEVKIKRGRSNLLWDTGVFRGQVYFKPQYHAEMCFLSWF<br>CGNQLPAYKCFQITWFVSWTPCPDCVAKLAEFLSEHPNVTLTISAARLYYYWERDYRRALCRLSQAGARVTIMDYEE<br>FAYCWENFVYNEGQQFMPWYKFDENYAFLHRTLKEILRYLMDPDTFTFNFNNDPLVLRRRQTYLCYEVERLDNGT<br>WVLMDQHMGFLCNEAKNLLCGFYGRHAELRFLDLVPSLQLDPAQIYRVTWFISWSPCFSWGCAGEVRAFLQENT<br>HVRLRIFAARIYDYDPLYKEALQMLRDAGAQVSIMTYDEFEYCWDTFVYRQGCPFQPWDGLEEHSQALSGRLRAIL<br>QNQGN |
| hAPOBEC-3C | *Homo sapiens* | MNPQIRNPMKAMYPGTFYFQFKNLWEANDRNETWLCFTVEGIKRRSVVSWKTGVFRNQVDSETHCHAERCFLS<br>WFCDDILSPNTKYQVTWYTSWSPCPDCAGEVAEFLARHSNVNLTIFTARLYYFQYPCYQEGLRSLSQEGVAVEIMDY<br>EDFKYCWENFVYNDNEPFKPWKGLKTNFRLLKRRLRESLQ |
| hAPOBEC-3D | *Homo sapiens* | MNPQIRNPMERMYRDTFYDNFENEPILYGRSYTWLCYEVKIKRGRSNLLWDTGVFRGPVLPKRQSNHRQEVYFRFE<br>NHAEMCFLSWFCGNRLPANRRFQITWFVSWNPCLPCVVKVTKFLAEHPNVTLTISAARLYYRDRDWRWVLLRLH<br>KAGARVKIMDYEDFAYCWENFVCNEGQPFMPWYKFDDNYASLHRTLKEILRNPMEAMYPHIFYFHFKNLLKACGR<br>NESWLCFTMEVTKHHSAVFRKRGVFRNQVDPETHCHAERCFLSWFCDDILSPNTNYEVTWYTSWSPCPECAGEVA<br>EFLARHSNVNLTIFTARLCYFWDTDYQEGLCSLSQEGASVKIMGYKDFVSCWKNFVYSDDEPFKPWKGLQTNFRLLK<br>RRLREILQ |

<table>
<tr><td>hAPOBEC-3F</td><td>Homo sapiens</td><td>MKPHFRNTVERMYRDTFSYNFYNRPILSRRNTVWLCYEVKTKGPSRPRLDAKIFRGQVYSQPEHHAEMCFLSWFCG<br>NQLPAYKCFQITWFVSWTPCPDCVAKLAEFLAEHPNVTLTISAARLYYYWERDYRRALCRLSQAGARVKIMDDEEFA<br>YCWENFVYSEGQPFMPWYKFDDNYAFLHRTLKEILRNPMEAMYPHIFYFHFKNLRKAYGRNESWLCFTMEVVKHH<br>SPVSWKRGVFRNQVDPETHCHAERCFLSWFCDDILSPNTNYEVTWYTSWSPCPECAGEVAEFLARHSNVNLTIFTA<br>RLYYFWDTDYQEGLRSLSQEGASVEIMGYKDFKYCWENFVYNDDEPFKPWKGLKYNFLFLDSKLQEILE</td></tr>
<tr><td>hAPOBEC-3G</td><td>Homo sapiens</td><td>MKPHFRNTVERMYRDTFSYNFYNRPILSRRNTVWLCYEVKTKGPSRPPLDAKIFRGQVYSELKYHPEMRFFHWFSK<br>WRKLHRDQEYEVTWYISWSPCTKCTRDMATFLAEDPKVTLTIFVARLYYFWDPDYQEALRSLCQKRDGPRATMKI<br>MNYDEFQHCWSKFVYSQRELFEPWNNLPKYYILLHIMLGEILRHSMDPPTFTFNFNNEPWVRGRHETYLCYEVER<br>MHNDTWVLLNQRRGFLCNQAPHKHGFLEGRHAELCFLDVIPFWKLDLDQDYRVTCFTSWSPCFSCAQEMAKFISK<br>NKHVSLCIFTARIYDDQGRCQEGLRTLAEAGAKISIMTYSEFKHCWDTFVDHQGCPFQPWDGLDEHSQDLSGRLRAI<br>LQNQEN</td></tr>
<tr><td>hAPOBEC-4</td><td>Homo sapiens</td><td>MEPIYEEYLANHGTIVKPYYWLSFSLDCSNCPYHIRTGEEARVSLTEFCQIFGFPYGTTFPQTKHLTFYELKTSSGSLVQ<br>KGHASSCTGNYIHPESMLFEMNGYLDSAIYNNDSIRHIILYSNNSPCNEANHCCISKMYNFLITYPGITLSIYFSQLYHTE<br>MDFPASAWNREALRSLASLWPRVVLSPISGGIWHSVLHSFISGVSGSHVFQPILTGRALADRHNAYEINAITGVKPYF<br>TDVLLQTKRNPNTKAQEALESYPLNNAFPGQFFQMPSGQLQPNLPPDLRAPVVFVLVPLRDLPPMHMGQNPNKP<br>RNIVRHLNMPQMSFQETKDLGRLPTGRSVEIVEITEQFASSKEADEKKKKKGKK</td></tr>
<tr><td>mAPOBEC-4</td><td>Mus musculus</td><td>MDSLLMKQKKFLYHFKNVRWAKGRHETYLCYVVKRRDSATSCSLDFGHLRNKSGCHVELLFLRYISDWDLDPGRCY<br>RVTWFTSWSPCYDCARHVAEFLRWNPNLSLRIFTARLYFCEDRKAEPEGLRRLHRAGVQIGIMTFKDYFYCWNTFVE<br>NRERTFKAWEGLHENSVRLTRQLRRILLPLYEVDDLRDAFRMLGF</td></tr>
<tr><td>rAPOBEC-4</td><td>Rattus norvegicus</td><td>MEPLYEEYLTHSGTIVKPYYWLSVSLNCTNCPYHIRTGEEARVPYTEFHQTFGFPWSTYPQTKHLTFYELRSSSGNLIQ<br>KGLASNCTGSHTHPESMLFERDGYLDSLIFHDSNIRHIILYSNNSPCDEANHCCISKMYNFLMNYPEVTLSVFFSQLYH<br>TENQFPTSAWNREALRGLASLWPQVTLSAISGGIWQSILETFVSGISEGLTAVRPFTAGRTLTDRYNAYEINCITEVKP<br>YFTDALHSWQKENQDQKVWAASENQPLHNTTPAQWQPDMSQDCRTPAVFMLVPYRDLPPIHVNPSPQKPRTV<br>VRHLNTLQLSASKVKALRKSPSGRPVKKEEARKGSTRSQEANETNKSKWKKQTLFIKSNICHLLEREQKKIGILSSWSV</td></tr>
<tr><td>MfAPOBEC-4</td><td>Macaca fascicularis</td><td>MEPTYEEYLANHGTIVKPYYWLSFSLDCSNCPYHIRTGEEARVSLTEFCQIFGFPYGTTYPQTKHLTFYELKTSSGSLVQ<br>KGHASSCTGNYIHPESMLFEMNGYLDSAIYNNDSIRHIILYCNNSPCNEANHCCISKVYNFLITYPGITLSIYFSQLYHTE<br>MDFPASAWNREALRSLASLWPRVVLSPISGGIWHSVLHSFVSGVSGSHVFQPILTGRALTDRYNAYEINAITGVKPFF<br>TDVLLHTKRNPNTKAQMALESYPLNNAFPGQSFQMTSGIPPDLRAPVVFVLLPLRDLPPMHMGQDPNKPRNIIRHL<br>NMPQMSFQETKDLERLPTRRSVETVEITERFASSKQAEEKTKKKKGKK</td></tr>
<tr><td>hAID</td><td>Homo sapiens</td><td>MDSLLMNRRKFLYQFKNVRWAKGRRETYLCYVVKRRDSATSFSLDFGYLRNKNGCHVELLFLRYISDWDLDPGRCY<br>RVTWFTSWSPCYDCARHVADFLRGNPNLSLRIFTARLYFCEDRKAEPEGLRRLHRAGVQIAIMTFKDYFYCWNTFVE<br>NHERTFKAWEGLHENSVRLSRQLRRILLPLYEVDDLRDAFRTLGL</td></tr>
<tr><td>ClAID</td><td>Canis lupus familiaris</td><td>MDSLLMKQRKFLYHFKNVRWAKGRHETYLCYVVKRRDSATSFSLDFGHLRNKSGCHVELLFLRYISDWDLDPGRCY<br>RVTWFTSWSPCYDCARHVADFLRGYPNLSLRIFAARLYFCEDRKAEPEGLRRLHRAGVQIAIMTFKDYFYCWNTFVE<br>NREKTFKAWEGLHENSVRLSRQLRRILLPLYEVDDLRDAFRTLGL</td></tr>
<tr><td>BtAID</td><td>Bos taurus</td><td>MDSLLKKQRQFLYQFKNVRWAKGRHETYLCYVVKRRDSPTSFSLDFGHLRNKAGCHVELLFLRYISDWDLDPGRCY<br>RVTWFTSWSPCYDCARHVADFLRGYPNLSLRIFTARLYFCDKERKAEPEGLRRLHRAGVQIAIMTFKDYFYCWNTFV<br>ENHERTFKAWEGLHENSVRLSRQLRRILLPLYEVDDLRDAFRTLGL</td></tr>
<tr><td>mAID</td><td>Mus musculus</td><td>MDSLLMNRRKFLYQFKNVRWAKGRRETYLCYVVKRRDSATSFSLDFGYLRNKNGCHVELLFLRYISDWDLDPGRCY<br>RVTWFTSWSPCYDCARHVADFLRGNPNLSLRIFTARLYFCEDRKAEPEGLRRLHRAGVQIAIMTFKDYFYCWNTFVE<br>NHERTFKAWEGLHENSVRLSRQLRRILLPLYEVDDLRDAFRTLGL</td></tr>
<tr><td>PmCDA-1</td><td>Petromyzon marinus</td><td>MAGYECVRVSEKLDFDTFEFQFENLHYATERHRTYVIFDVKPQSAGGRSRRLWGYIINNPNVCHAELILMSMIDRHL<br>ESNPGVYAMTWYMSWSPCANCSSKLNPWLKNLLEEQGHTLTMHFSRIYDRDREGDHRGLRGLKHVSNSFRMGV<br>VGRAEVKECLAEYVEASRRTLTWLDTTESMAAKMRRKLFCILVRCAGMRESGIPLHLFTLQTPLLSGRVVWWRV</td></tr>
<tr><td>PmCDA-2</td><td>Petromyzon marinus</td><td>MELREVVDCALASCVRHEPLSRVAFLRCFAAPSQKPRGTVILFYVEGAGRGVTGGHAVNYNKQGTSIHAEVLLLSAV<br>RAALLRRRRCEDGEEATRGCTLHCYSTYSPCRDCVEYIQEFGASTGVRVVIHCCRLYELDVNRRRSEAEGVLRSLSRLG<br>RDFRLMGPRDAIALLLGGRLANTADGESGASGNAWVTETNVVEPLVDMTGFGDEDLHAQVQRNKQIREAYANYA<br>SAVSLMLGELHVDPDKFPFLAEFLAQTSVEPSGTPRETRGRPRGASSRGPEIGRQRPADFERALGAYGLFLHPRIVSRE<br>ADREEIKRDLIVVMRKHNYQGP</td></tr>
<tr><td>PmCDA-5</td><td>Petromyzon marinus</td><td>MAGDENVRVSEKLDFDTFEFQFENLHYATERHRTYVIFDVKPQSAGGRSRRLWGYIINNPNVCHAELILMSMIDRHL<br>ESNPGVYAMTWYMSWSPCANCSSKLNPWLKNLLEEQGHTLMMHFSRIYDRDREGDHRGLRGLKHVSNSFRMGV<br>VGRAEVKECLAEYVEASRRTLTWLDTTESMAAKMRRKLFCILVRCAGMRESGMPLHLFT</td></tr>
<tr><td>yCD</td><td>Saccharomyces cerevisiae</td><td>MVTGGMASKWDQKGMDIAYEEAALGYKEGGVPIGGCLINNKDGSVLGRGHNMRFQKGSATLHGEISTLENCGRL<br>EGKVYKDTTLYTTLSPCDMCTGAIIMYGIPRCVVGENVNFKSKGEKYLQTRGHEVVVVDDERCKKIMKQFIDERPQD<br>WFEDIGE</td></tr>
<tr><td>pYY-BEM3.1</td><td>tr|F7B644|F7B644_HORSE</td><td>MPRGRARERQRRNPMEKLDAEAFSFHFLNMEFVYDRNCSYLCYQVEGRLSGSPVLSEQGVFPNEVCGKTRRHAEL<br>CFLDWFRGRLSPDEYYCVTWFISWSPCSNCAREVAEFLKRHRNVELSIFAARLYYCRDHEQGLQSLCNRGAQLAVM<br>LRKDFTYCWDNFVHNSGREFSPWENIDANSDLLARKLEDLLKNPMEKLHRKTFSFHFRNLKFAKGRKCSYLCYRVEG<br>RLSGSPGLSEQGVFLNEVCDENCRHAELCFLHWFRGRLSPHADYRVTWFISWSPCSNCAREVAEFLKQHRNVELHIS<br>AARLYYWQRNKPGLRNLRSSGAQLAIMFFWDFRDCWDNFVHNSGRHFIPWKKINVNSRLLATKLEDLLKNPLEKLH<br>PNTFSFHFCNLEFAYDRKYSYLCYQVEGRLSGSPGLSEQGVFLNEVCGKTRCHAELCFLDWFRVRLSPDEYYRVTWFI<br>SWSPCFYCAREVADFLKQYRNVKLSIFAARLYYCRDHAQGLRSLCSSGAQLAIMFFWDFRYCWDNFVHNSGREFRP<br>WKKINVNSRLLATKLEDILK</td></tr>
<tr><td>pYY-BEM3.2</td><td>tr|D1LZA1|D1LZA1_PANTI</td><td>MEPWRPSPRNPMDRIDPKTFRFQFPNLRYASGRKLCYLCFQVERDYFYYNDSDWGVFRNEVHPWAPCHAEQCFL<br>SWFRDQYPYRDEDYNVTWFLSWSPCPTCAEEVVEFLEEYRNLTLSIFTSRLYYFWHPNYQEGLCKLWDAGVQLDIM<br>SCDEFEYCWDNFVYHKGMRFQRRNLLKDYDFLAAKLQEILSPGQQRKRDWPFPPRPGAQVDPRSWVQEVTEPGI</td></tr>
</table>

| | | |
|---|---|---|
| | | NTRRHPLHLLVSFLLPRPTMNPLQEDIFYRQFGNQHRVPKPYYYRRKTYLCYQLKLPEGTLIDKDCLRNKKKRHAEICFIDKIKSLTRDTSQRFEIICYITWSPCPFCAEELVAFVKDNPHLSLRIFASRLYVHWRWKYQQGLRHLHASGIPVAVMSLPEFEDCWRNFVDHQDRLFQPWRNLDQYSESIKRRLGKILTPLNDLRNDFRNLKLE |
| pYY-BEM3.3 | tr\|A0A3Q0DM17\|A0A3Q0DM17_TARSY | MPMKRMYSNIYFDHFNNQRLLSGQNAPWLCFKVERVENCMLVPLETGVFGNQVSGCCGKTERPVEPTSLTRSVLVSPNPGTELRAQQPSRKGHLGKLGCVEYPSPGLALVMLGYGASTYCPDSSMYCPETCHHPEMCFLYWFEKTLSHEEQYQITWYVSWSPCVNCAEEVAEFLSVHPKVNLTIYAARLYCYQKLNHRQGLRRLCKEGACVKIMNYEEFDHCWENFVYNNYKSFKPWVKLQDNYELLATELDKILRIPMERMPQKKFRFHFQNLIAKDRNTTWLCFEVKNVRKKHPPDLLERGIFQNQVTPRINCHAEMCFLSWFLENMLLHGKRYQVTWYISWSPCSICAEEVAEFLSAHPKVSLTIYAARLYYFWVPGYRQGLRRLVEEGARVEIMNYEEFDYCWENFVSINNEPFQPWEGLHEKYGYLVTKLNNILG |
| pYY-BEM3.4 | tr\|A0A3Q0DNJ5\|A0A3Q0DNJ5_TARSY | MEDNPEPRPRQQMDQDTFIFNFNNDPSVRGRHQTFLCYEVEHLDDDTWVPQDKYLGFLHNQPQSRSNAYCAYHAELCFLELVSSWQLDPAQRYRVTCFISWSPCSSCAQEVAAFLKKNRHVTLRILAARIYDYYQGYEDGLRTLQGVGVDITVMTSAEFGHCWNTFVDHQGSPFQPWEGLDQHSQVIWQRMQDILQVIPAKYLMEKVKYTVTVDILFKGRVPGPRYLMDQNTFTRNFINNLSVSGRRQTLLCYEVERLGGDIWVPLDQLRGFLLSQARDVLNYYQGRHAEPCFLDLVSSWQLDPAQHYRVTWFISWSPCTSCAQAVAAFLRENRHVTLRILAARIYDYHQGYEEGLRTLQRTGAHIDIMTFKEFGHCWNTFVNHKGSPFKSWTGLDQHSQALRKRLQDILHTMASSLWDQSEPKKPIPSQEVTLPESIPPSHGNRFRLVKRPS |
| pYY-BEM3.5 | tr\|G5AYU5\|G5AYU5_HETGA | FCFLSCVHRKPIERIYKKAFRFYFRNLRCAYGRNKTFLCYEVKRERDNKVLHKGVVLNQVEPYMPLHAELRFLSWFHDTLLCPLGSYQVTLYVSWSPCSECAEELTTFLAGHRNVTMTIYVAQLYYCNWKSPNREGLKILIAEDARLRVMFYDEFLYCWRNFVKNDYNNFDPWSLLDENSRYHNRILQNILKGWGRPHRVGPEGEQTATPGGSGGHCISVFSLLRRREMTLKEETFRVQFNNAYKAPKPYRRRVTYLCYQLQEANGDPLTKGCLRTKKGYHAESRFIKRICSMDLGQDQSYQVTCFLTWSPCPHCAQELVSFKRAHPHLRLQIFTARLFFHWKRSYQEGLQRLCRAQVPVAVMGHPEFAYCWDNFVDHQPGPFEPPWAKLEYYSSCLKRRLQQILRSWGVDDLTNDFRNLQLGP |
| pYY-BEM3.6 | tr\|A0A2Y9QMV5\|A0A2Y9QMV5_TRIMA | MLSSPQTPGTRKPMKTLAPDEFSFNFENLRLAHGRNTTFLCFQVETKAPPSLNSPDSGIFQNQDHCPSHHHAEMVFLTWFQKRLSPAQHYEVTWYMSWSPCSRCAVQVAKFLKSNSTVNLSIFVARLYYPRELETKDGLHSLWQAGAQVQIMFFQDFKYCWENFVNNEGKPFQPWKNLDENSKDWDTELKDIHRNTTDLLTEEMFYSQFYNREKKSSIPRKTYLCYQLNEPQPVKRCLHYKKGYHAVTRFIDGIVSMNLDPARSYDITCYFTWSPCNRYARKLVSFIEDYPNLRLKVYTSRLYFHWCWTNMQGLQHLQNSRVTVAVMTFRDFEYCWKNFVDNQGKPFEPWEKLDLYSQSTERRLRRILKPLTPDVLNEDFGNLHL |
| pYY-BEM3.7 | tr\|H0XHI0\|H0XHI0_OTOGA | LSCAFRDPMNRMYPKTFCQNFEKEPCPSNQNSSWLCFEVETKNSAVFFHRGVFRNQPAPPPRAPTSVLLSQGPVKTPCHAEECFLTWIQGVLPPDHHYHVTWYVSRGPCANCANLIVHFLAMHRRVTLTIFAAHLNFFWESDFQQGLLRMDQEGVQLHIMGYEEFEYCWDNFVYNQRKQFVPWNGLNENYEFMVSTLEDILRSPLDRIRQKDFSIHFRNSLWLDDKSTWLCFEVKRTKSPVPLYRGVFRNQSPPKTPCHAEVRFFTWLQDLPPDFCCQFTWYLSWSPCADCADLVANFLAKHRNVSLTIFVARLYYYRDPEMHRGLRRMYQEGANVDIMSVIEFEYCWDNFVYNQGKQFVPWNGLNENYEFLVPRLQEILE |
| pYY-BEM3.8 | tr\|A0A3M0K4Y7\|A0A3M0K4Y7_HIRRU | MYISKKALRRHFDPRVYPRETYLLCELQWEGSRRVWIHWIRNVPDHHAEEYFLEEVFEPRNYGFCNITLYLSWSPCCTCCSKIRDFLKRNPNVKIDIRVARLIYPDYAETRSSLRELNGLQRVSIQVMEAAGLSCIESKNHRISQVERDPKGSSSPTLFTLQDHLKLSNMTESVIQDSVSIQICYQMRILGFQCHIRWKLQPEDFQRNYSPNQIGRVVYLLYEVRWRRGSIWRNWCSNNPEQHAEVNFLENHFHHRPQTPCSITWFLSTSPCGKCSRRILEFLKSQPNVTLEIYAAKLFRHHDIRNRQGLRNLMMNGVTIYIMNLEGNPASLCLSVD |
| pYY-BEM3.9 | tr\|A0A3P4LUZ8\|A0A3P4LUZ8_GULGU | MSFEDYEYCWETFVDHKGMYFQSWDLLRDNDLLAAELKNILRSTMNPLRQEIFYHQFGNQPRAPRPYHRRKTYLCYQLQPHEGPITARVCLQNKKKRHAEIRFIDNIRALRLDRSQTFEITCYLTWSPCPTCAKALAVFVQDHPHISLRLFASRLFIHWCWKYQEGLRLLHRSRIPVAVMRLQEFEDCWRNFVDNQDEPFQPWNKLEQYSESITRRLRRILGHPQNNLENDFRNLHI |
| pYY-BEM3.10 | tr\|G5BPM8\|G5BPM8_HETGA | RRRIEPWQFEASFDPRQLRRETCLLSEVRWGTSPRAWRGCSLNTARHAEVSFMDRLTSEGRLRGPVRCSITWFLSWSPCGACAQAIGEFLRQHPNVSLVIYIARLFWHVDEQNRQGLRDLVTRGVRMQVMSDPEFAHCWRNFVNYSPGQEARWPQVPPVWTWLYSLELHCILLNLPPCLKISRRHHNQLTFFQLILQNCHYQAIPSPVLLASGLIHPFVTW |
| pYY-BEM3.11 | tr\|H2M862\|H2M862_ORYLA | MITKLDSVLLPKKKFIYHYKNMRWARGRHETYLCFVVKRRVGPESLSFDFGHLRNRNGCHVELLFLRHLSALCPGLWGYGATGQGRVSYSITWFCSWSPCANCSFRLAQFLSQTPNLRLRIFVSRLYFCDLEDSREREGLRMLKKVGVHITVMSYKDYFYCWQTFVARKQSKFKPWDGLHQNSVRLSRKLNRILQPCETEDFRDAFKLLGL |
| pYY-BEM3.12 | tr\|H0Y0C6\|H0Y0C6_OTOGA | MYLKTFYRHFNNRPYLSRRNDTWLCFEVKTTSSNSPGSFYSGVFRNQGPRYCPWHTELCFLTWVRPIVSHHHFYQITWYMSWSPCANCAWQVATFLATHENVSLTNYTVRIYYFWRQDYRQGLLRMIEEGTQVYVMSSKEFQHCWENFVDHWGTRWVTCWNRLKKNYEFLVTRLSEILSDPKERISPNTFYNQFNNTPVPRGRKDTWLCFEVKEKNSNSPGSFHRGVFQNQVFSGTSSHARRCPPDHHYEVTWYTSWSPCAHCAWHVVNFLTSNPNVSLTIFAARLYYIYRPEIQQGLRRVFQEGAKVHIMSLKEFKYCWAKLVYNSGMRFMPWYQFNFNFLFPNTTLKGDLH |
| pYY-BEM3.13 | tr\|A0A3Q2Z5X6\|A0A3Q2Z5X6_HIPCM | MDVHFMNFIYHYKNMRWAKGRNETYLCFVVKRRVGPNSLTFDFGHLRNRNGCHVELLFLRYLGRRLSYSITWFCSWSPCANCSAALSQFLSRMPNLRLRIFVARLYFCDMEDSHEREGLRLLQKAGVQVTVMSYKDYYYCWQTFVDRKKSHFKAWEDLHQNSVRLSRKLNRILQPCEMDLRDAFKLLGL |
| pYY-BEM3.14 (RrA3F) | tr\|A0A2K6NVA7\|A0A2K6NVA7_RHIRO | MKPQIRDHRPNPMEAMYPHIFYFHFENLEKAYGRNETWLCFTVEIIKQYLPVPWKKGVFRNQVDPETHCHAEKCFLSWFCNNTLSPKKNYQVTWYTSWSPCPECAGEVAEFLAEHSNVKLTIYTARLYYFWDTDYQEGLRSLSEEGASVEIMDYEDFQYCWENFVYDDGEPFKRWKGLKYNFQSLTRRLREILQ |
| pYY-BEM3.15 | tr\|A0A2K6NY90\|A0A2K6NY90_RHIRO | MNPHIRNPMEAMYPGTFYFHFKNLWEADNRNESWLCFAVEVIKHHSTVSWKRGVFRNQVDPETHCHAEKCFLSWFCDNTLSPKKNYQVTWYTSWSPCPECAREVAKFLARHSNVMLTIYTARLYYSQYPNYQEGLRRLNEEGVPVEIMDYEDFKYCWENFVYNGDELFKPWKGLKYNFLFLDSKLQEILE |
| pYY-BEM3.16 | tr\|Q6ICH2\|Q6ICH2_HUMAN | MNPQIRNPMERMYRDTFYDNFENEPILYGRSYTWLCYEVKIKRGRSNLLWDTGVFRGPVLPKRQSNHRQEVDPETHCHAERCFLSWFCDDILSPNTNYEVTWYTSWSPCPECAGEVAEFLARHSNVNLTIFTARLCYFWDTDYQEGLCSLSQEGASVKIMGYKDFVSCWKNFVYSDDEPFKPWKGLQTNFRLLKRRLREILQ |

| | | |
|---|---|---|
| pYY-BEM3.17 | tr\|G8GPV1\|G8GPV1_CERNE | MDGSPASRPGHVMDPGTFTSNFNNKPWVSGQRETYLCYKVERSHNDTWVLLNQHRGFLRNQAKNRLHGDYGCHAELCFLGEVPSWRLDPTQTYRVTWFISWSPCFSGGCAEQVRAFLQENTHVRLRIFAARIYDYDFLYQEALRTLRDAGAQVSIMTYEEFKHCWDTFVDHQGRPFQPWDGLDEHSQALSGRLQAILQNQGN |
| pYY-BEM3.18 | tr\|Q1WBT6\|Q1WBT6_SYMSY | MALLTAKTFRLQFNNKRRVTKPYYPRKALLCYQLTPQNGSTPTRGYFKNKKKRHAEIRFINKIKSMGLDETQCYQVTCYLTWSPCPSCAWELVDFIKAHDHLNLGIFASRLYYHWCRHQQEGLRLLCGSQVPVEVMGFPEFADCWENFVDHEEPLSFNPSEMLEELDKNSRAIKRRLEKIK |
| pYY-BEM3.19 | tr\|A0A3B4CS14\|A0A3B4CS14_PYGNA | MDNTNRRKFIYHYKNVRWARGRHETYLCFVVKKRNSPDSLSFDFGHLRNRNGCHVELLFLRYIEVLCPGLWGSGVDGVRVSYAVTWFCSWSPCSNCAQRLTNFLSQTPNLRLRIFVARLYFCDEEDSLEREGLRHLQRAGVQITVMTYKDFFYCWQTFVASRERCFKAWEGLRQNSVRLSRKLNRILQVFISTPVISPLITTHLGQSWAGG |
| pYY-BEM3.20 | tr\|A0A087XZI4\|A0A087XZI4_POEFO | RKVSYSVTWFCSWSPCANCSIRLAQFLHQTPNLRLRIFVSRLYFCDLEDSREREGLRILKKAGVHITVMSYKDYFYCWQTFVAKSQSKFKPWDGLHQNYIRLSRKLNRILQPALDIKKFIYHYKNLRWARGRCETYLCFVVKKKLHLFMFVIVGRNRLFDLNVTMNNKSLYLIPLHLQLLFLRHLGALCPGLWGYGVTGERKVSYSVTWFCSWSPCANCSIRLAQFLHQTPNLRLRIFVSRLYFCDLEDSREREGLRILKKAGVHITVMSYKDYFYCWQTFVAKSQSKFKPWDGLHQNYIRLSRKLNRILQVQFF |
| pYY-BEM3.21 | tr\|A0A341AEK4\|A0A341AEK4_9CETA | MASDRGPSAGDATSRRRIEPWEFEVSFDPRELCKETRLLYEIKWGRSQHVWRHSGKNTTNHVECNFIEKFTSERPFHRSVSCCITWFLSWSPCWECSKAIREFLNQHPRVTLFIYVARLFQHMDPQNRQGLRDLIHSGVTIQIMGPTEYDYCWRNFVNYPPGKEAHWPRYPPPLMKLYALELHCIILVP |
| pYY-BEM3.22 | tr\|E2D879\|E2D879_MUSMI | RNLISRETFNFNFENLCYAKGRKNTFLCYEVTRKDCDSPVSLCHGVFKNKGSIHAEICFLYWFHDKVLKVLTPREEFKVTWYMSWSPCFECAEQVVRFLATHHNLNLTIFSSRLYNVSDPDTQQKLCRLVQEGAQVAVMDLSEFKKCWEKFVDNDGQQFRPWKRLRTNFRYQNSKLQEIL |
| pYY-BEM3.23 | tr\|A0A2K5RDN6\|A0A2K5RDN6_CEBCA | MWEAQSPGLSREWGSVAISPEDPGPLHIGRFLSCAFRHPMNAMYPGIFNFHFRNLRKAYGRNETWLCFTVEGIMNRSTVSWKSGVFRNQVGSDPFCHAEMCFLSWFRHNMLSPKKDYEVTWYASWSPCPECAGQVAEFLARHGNVRLTIFTAHLYYFWNPSFRQGLRRLSQEGASVLIMGYEDFEYCWDNFVYNDGQPFKPWKRLQDNSLSLYITLQEILQ |
| pYY-BEM3.24 | tr\|A0A2K5RDN7\|A0A2K5RDN7_CEBCA | MEASPASRPRPLMGPRTFTENFTNNPEVFGRHQTYLCYEVKCQGPDGTRDLMTEQRDFLCNQARNLLSGFDGRHAERCFLDRVPSWRLDPAQTYRVTCFISWSPCFSCAREVAEFLQENPHVNLRIFAARIYDCRPRYEEGLQMLQNAGAQVSIMTSEEFRHCWDTFVDHQGHPFQPWEGLDEHSQALSRRLQAILQGNRWMILSL |
| pYY-BEM3.25 | tr\|A0A1C9CJ69\|A0A1C9CJ69_CERAL | NPMKAMDPHIFYFHFKNLRKAYGRNETWLCFAVEIIKQRSTVPWRTGVFRNQVDPESHCHAERCFLSWFCEDILSPNTDYRVTWYTSWSPCLDCAGEVAEFLARHSNVELAIFAARLYYFWDTHYQQGLRSLSEKGASVEIMGYEDFKYCRENFVCDDGKPFKPWKGLKTNFRFLKRRLQEILE |
| pYY-BEM3.26 | tr\|A0A2R2Z4D2\|A0A2R2Z4D2_PTEAL | MHLQVWRKVTEAWREGYTLKPWSRNPMERLYHDYFYFHFYNLPTPKHRNGCYICYQVEGTKKHSRMPLLRGVFENQESLDMMLSPGEKYRVTWYISWSPCFACVDEVIKFLREHTNVELIIFAARLYHSDILQYRQGLRKLHDAGVHVAIMSYYEFKHCLNDFVFHQGRSFCPWNDLNKNSKNLSNTLEDILRNQED |
| pYY-BEM3.27 | tr\|B7T161\|B7T161_SHEEP | MTEGWAGSGLPGRGDCVWTPQTRNTMNLLRETLFKQQFGNQPRVPPPYYRRKTYLCYQLKELDDLMLDKGCFRNKKQRHAEIRFIDKINSLNLNPSQSYKIICYITWSPCPNCASELVDFITRNDHLNLQIFASRLYFHWIKPFCRGLHQLQKAGISVAVMTHTEFEDCWEQFVDNQLRPFQPWDKLEQYSASIRRRLQRILTAPT |
| pYY-BEM3.28 | tr\|A0A2R2X2G4\|A0A2R2X2G4_PTEAL | MAGLGQACEGCCGQMPEISYPMGRLDPKTFSFEFKNLPYAYGRKSSYLCFQVEREQHSSPVPSDWGVFKNQFCGTEPYHAELCFLNWFRAEKLSPYEHYDVTWFLSWSPCSTCAEEIAIFLSNHKNVRLNIFVSRIYYFWKPAFRQGLQELDHLGVQLDAMSFDEFKYCWENFVDNQGMPFRCWKKVHQNYKSVLRKLNEILRRR |
| pYY-BEM3.29 | tr\|G1Q1M4\|G1Q1M4_MYOLU | YAELSFLDLFQSWNLDRGRQYRLTWYMSWSPYPDCAQKLVEFLGENSHVTLRIFAADIHSLCSGYEDGLRKLRDARAQLAIMTRDELQYCWVTFVDNQGQPFRPWPNLVEHIKTKKQELKDILGNPMRRMYPKTFNFNFQNLNSYGRKSTFLCFEVETWEDGSVLDYQNGVFQNQLDPGHAELCFIEWFHEKVLFPDEVRCPDAQYHVTWYISWSPCFECAEQVAGFLNEHENVDLSISAARLYLCEDEDEQGLQDLVAAGAKVAMMAPEDFEYCWDNFVYNRGWPFTYWKHVRRNYGRLQEKLDEILW |
| pYY-BEM3.30 | tr\|A0A1S3AN78\|A0A1S3AN78_ERIEU | RRIEPWEFEDFFDPRQFRPETCLLYEVRWGSSRNAWRSTARNTTRHAEVNFLERFAAERHFDKPVSCSITWFLSWSPCWECSQAIGAFLSQHPQVTLAIHVTRLFHHEDEQNRQGLRDLLARGVTLQVMGDSEYAHCWRTFVNSPPGAEGHYPRYPSDFTRLYALELHCIILGLPPCLEILRRYQNQFTLFRLVPQNCHYQMIPHLNFFVVRHYFF |
| pYY-BEM3.31 (AmAPOBEC1) | tr\|A0A151P7C9\|A0A151P7C9_ALLMI | MADSSEKMRGQYISRDTFEKNYKPIDGTKEAHLLCEIKWGKYGKPWLHWCQNQRMNIHAEDYFMNNIFKAKKHPVHCYVTWYLSWSPCADCASKIVKFLEERPYLKLTIYVAQLYYHTEEENRKGLRLLRSKKVIIRVMDISDYNYCWKVFVSNQNGNEDYWPLQFDPWVKENYSRLLDIFWESKCRSPNPW |
| pYY-BEM3.32 | tr\|Q4VUI3\|Q4VUI3_XENLA | MTMDSMLLKRNKFIYHYKNLRWARGRHETYLCYIVKRRYSSVSCALDFGYLRNRNGCHAEMLFLRYLSIWVGHDPHRNYRVTWFSSWSPCYDCAKRTLEFLKGHPNFSLRIFSARLYFCEERNAEPEGLRKLQKAGVRLSVMSYKDYFYCWNTFVETRESGFEAWDGLHENSVRLARKLRRILQPPYDMEDLREVFVLLGL |
| pYY-BEM3.33 | tr\|E2RL86\|E2RL86_CANLF | MNPLQEETFYQQFSNQRVPKPTYQRRTYLCYQLKPHEGSVIAKVCLQNQEKRHAEICFIDDIKSRQLDPSQKFEITCYVTWSPCPTCAKKLIAFVNDHPHISLRLFASRLYFHWRQKYKRELRHLQKSGIPLAVMSYLEFKDCWEKFVDHKGRPFQPWNKLKQYSESIGRRLQRILQPLNNLENDFRNLRL |
| pYY-BEM3.34 | tr\|G1LWB0\|G1LWB0_AILME | SSAAPASIHLLDEDTFTENFRNDDWPSRTYLCYKVEGPDQGSGVPLGQDKGILHNKPAQGPEPSRHAECYLLEQIQSWNLDPKLHYGVTCFLSWSPCAKCAQKMARFLQENSHVSLKLFASRLYTRERWDEDYKEGLRTLKRAGASIAIMTYREFEHCWKTFVLHDQEGSCFQPWPFLHKESQKFSEKLQAILQVGVLLLSLPPPLPSSPLSSPWPFPAPLRASTG |
| pYY-BEM3.35 | tr\|A0A1U7S7K7\|A0A1U7S7K7_ALLSI | MGEHWQYAGSGEYIPQDQFEENFDPSVLLAETHLLSELTWGGRPYKHWYENTEHCHAEIHFLENFSSKNRSCTITWYLSWSPCAECSARIADFMQENTNVKLNIHVARLYLHDDEHTRQGLRYLMKMKRVTIQVMTIPDYTYCWNTFLEDDGEDESDDYGGYAGVHEDEDESDDDDYLPTHFAPWIMLYSLELSCILQGFAPCLKIIQGNHMSPTFQLHVQDQEQKRLLEPANPWGAD |
| pYY-BEM3.36 | tr\|A0A2R2X2J8\|A0A2R2X2J8_PTEVA | MPRIGNMNLLSEKTFNYHFGNQLRVKKPQGRRRTYLCYKLKLPNETLVKGYFINKKKNHAEIRFINKIRSLNLDQTQSYKITCYITWSPCSYCAGKLVALVKSCPHLSLQIFTSRLYYHWLWKNQAGLRYLWKINISVLVMKEPEFADCWDNFVNHQSRRFKPWEKLTQYSNSTERRLLRILRINRTDLFLAQSSEQDPGLNDLVDAIKRLFLDAHRPRD |

<table>
<tr><td>pYY-BEM3.37</td><td>tr|A0A151P6M4|A0A151P6M4_ALLMI</td><td>MAVEEEKGLLGTSQGWKIELKDFQENYMPSTWPKVTHLLYEIRWGKGSKVWRNWCSNTLTQHAEVNCLENAFGKLQFNPPVPCHITWFLSWSPCCQCCRRILQFLRAHSHITLVIKAAQLFKHMDERNRQGLRDLVQSGVHVQVMDLPDYRYCWRTFVSHPHEGEGDFWPWFFPLWITFYTLELQHILLQQHALSYNL</td></tr>
<tr><td>pYY-BEM3.38</td><td>tr|A0A2K6MNR2|A0A2K6MNR2_RHIBE</td><td>IWLCFTMEIIKQCSTVSWKRGVFRNQVDPETHCHAERCFLSWFWEDTLSPNTNYQVTWYTSWSPCLDCAGEVAEFLARHSNVKLAIFAARLYYFWDTDYQQGLRSLSEEGTSVEIMGYEDFKYCWENFVYNGDEPFKPWKGLKYNFLFLDSKLQEILE</td></tr>
<tr><td>pYY-BEM3.39 (SsAPOBEC3B)</td><td>tr|D3U1S2|D3U1S2_PIG</td><td>MDPQRLRQWPGPGPASRGGYGQRPRIRNPEEWFHELSPRTFSFHFRNLRFASGRNRSYICCQVEGKNCFFQGIFQNQVPPDPPCHAELCFLSWFQSWGLSPDEHYYVTWFISWSPCCECAAKVAQFLEENRNVSLSLSAARLYYFWKSESREGLRRLSDLGAQVGIMSFQDFQHCWNNFVHNLGMPFQPWKKLHKNYQRLVTELKQILREEPATYGSPQAQGKVRIGSTAAGLRHSHSHTRSEAHLRPNHSSRQHRILNPPREARARTCVLVDASWICYR</td></tr>
<tr><td>pYY-BEM3.40</td><td>tr|F1CGT0|F1CGT0_ANOCA</td><td>KAAILLSNLFFRWQMEPEAFQRNFDPREFPECTLLLYEIHWDNNTSRNWCTNKPGLHAEENFLQIFNEKIDIKQDTPCSITWFLSWSPCYPCSQAIIKFLEAHPNVSLEIKAARLYMHQIDCNKEGLRNLGRNRVSIMNLPDYRHCWTTFVVPRGANEDYWPQDFLPAITNYSRELDSILQD</td></tr>
<tr><td>pYY-BEM3.41</td><td>tr|C7AGG3|C7AGG3_HORSE</td><td>MDPQAPTQRGGLGQAYQGGDYVQAPGNGNTQHLLSEDVFKKQFGNQRRVTKPYYRRKTYVCYQLKLLRGPTIAKGYFRNKKKRHAEIRFIDKINSLGLDQDQSYEITCYVTWSPCATCACKLIKFTRKFPNLSLRIFVSRLYYHWFRQNQQGLRQLWASSIPVVVMGYQEFADCWENFADNRGNPFQSWEKLTEYSKGIKRRLQKILEPLNLNGLEDAMGNLKLGSVDLG</td></tr>
<tr><td>pYY-BEM3.42</td><td>tr|A0A250YMK7|A0A250YMK7_CASCN</td><td>MSLLKEDIFLYQFNNQQQVQKPYFRRRTYLCYQLEQPNGSRPQWPAKGCLQNKKGHHAEIRFIKRIHSMGLEQDQDYQITCYITWSPCLACACALAELKNHFPRLTLRIFASRLYFHWIRKFQMGLQHLYKSGVLVAVMSLPEFTDCWEKFVNHRQVFFTPWDKLEEHSRSIQRRLRRILQSWDVDDLTDDFRNLRL</td></tr>
<tr><td>pYY-BEM3.43</td><td>tr|B7T160|B7T160_SHEEP</td><td>MPWISDHVARLDPETFYFQFHNLLYAYGRNCSYICYRVKTWKHRSPVSFDWGVFHNQVYAGTHCHSERRFLSWFCAKKLRPDECYHITWFMSWSPCMKCAELVAGFLGMYQNVTLSIFTARLYYFQKPQYRKGLLRLSDQGACVDIMSYQEFKYCWKKFVYSQRRPFRPWKKLKRNYQLLAAELEDILG</td></tr>
<tr><td>pYY-BEM4.1</td><td>tr|A0A182D0J1|A0A182D0J1_BLAVI</td><td>MTNPESPPQAPCDFNEDALLNREPLRGSPIKFVSPVDYPDLVFALAGPVGVDIDYIQQSISDCLKSFDYSTEFIRITEIMQDIKCSKTIDCTDMLKEYQSKIEYANELRRAYRAKDLLAALTISAISKLREQIKERDEATNKSNIQPSRRKLAWVVRQLKTPEEVRLLRAVYGKQFVLVSIYSSPQRREDFLISKIKIKSRGTIDNNTSSEGAQRLIERDSKEDNEYGQNLSGTFCLGDIFVDSNNKESAIVSIDRFLNAFFGSNEISPTRDEYGMYLAKTASLRSCDLSRQVGAAIFSKTGEIISLGSNEVPKAGGGTYWTGDNADSRDIRLGHDPNEINKVEIFAEIISRLLEDKLLSNDLLNKDAASIVTILLSKNEGKRYKDLRVMDIIEFGRIIHAEMSAICDAARNGRAIIGATLFCTTFPCHLCAKHIVASGIGRIVYLEPYPKSYAKKLHSDSIQVEDHSDSEKVSFEPFIGISPSRYRELFEGGRRKDPFGEALKWKNDPRKPVIDVVVPPHFEAEKLVIAQLGKLIVSGTG</td></tr>
<tr><td>pYY-BEM4.2</td><td>tr|A0A2D6EXD2|A0A2D6EXD2_9ARCH</td><td>MIIGLVGTIGAGKQTIIDYLQEKYGYNALSCSDVLREILKKQGKPVTRDNLREIGNKTREEGGNGAIAKILLEKLRNNWKANYIVDSLRHPDEVSVLRTSPLFHLVAVDADLRIRFERVKARKREEEPTTLPAFVERDQKEMFGTGNEQRIRETMELADELVLNNGTVEELKQRIDDLNLVSDERLRPSWDDYFMRLARLAAQRSNCMSRKVGAIITKDRRVIATGYNGTPRGVKNCNEGGCERCNSAVAKGTAISECLCLHGEENAIIEAGRVRSEGATIYTSFLPCLWCTKMIIQAGLKEVVFSEVYDLHEASIKLFETSGVLIRRLK</td></tr>
<tr><td>pYY-BEM4.3</td><td>tr|F7YVM7|F7YVM7_9THEM</td><td>MNEFKYMSLALKLAKKGKYTTSPNPMVGAVIVKDGKILATGYHKKAGQPHAEINALSKLNFQAQNCEMYVTLEPCSHYGRTPPCADAIIRSGIRKVVIATLDPNPLVNGKGVEKLKNAGIEVVCGVLEEKAKKLNEKFFKYITTKIPFVALKIAQTLDGKIALKNGESKWITSEKSREYVHKLRMEYDAVLTGIGTILKDDPQLNVRLKKVYKQPLRIILDSKLKIPLSAKVLEDPSKVIILTTALADKEKLEELRSKGVEVIITNEKNGIVDLESALKILGEKKITSVMVEAGPTLLTSFLKESLFDKIYLFIAPKIFGADSKSVFSELGLEDISKSQKFSLESVKKIGEDLLLELYPKQLKKLEE</td></tr>
<tr><td>pYY-BEM4.4</td><td>tr|A0A3M6UNF1|A0A3M6UNF1_9CNID</td><td>MEEKSELENELMRSTSPKPSVPNGSKGNECEQRETRITKENLYMVLALWMEEFPVVEQTSSAKRLNKVGVVFVLPTDRVLAADCSRDGVHGVARVMVNHCGKLEGCKVFVSRKPCSLCAKLLVQSKVSRVFYLPIEPESENKGEIARADNLFKNSSVGQSVFVPCVEQKVLDKLEDKLPKEIITPDDISECRDNLLKKCGWSAEWFARAQASLPWPCFEGKMKSQVDNDFKSLIKWIAVVKAPMDKGVAFPKVKLTSDSRVVPDCDADNFPDSKTAYHMMIFAKMLARQTDDPKTGVGAVIVRGKVPDIVSLGWNGFPSKALYGEFPRASDDDRALQKKFPYVIHAEQNALMVRNVKDLTDGILFVTKPPCDECAPMIKLSGVKTIVIGEKIEKSRGGELSYNLIKEYIKEGIMTCYQMEATKTKAKRLASDPETRKRLKSSCSNSNDV</td></tr>
<tr><td>pYY-BEM4.5</td><td>tr|A0A2G3K826|A0A2G3K826_9BURK</td><td>MTKIIDDVNTAAAAVLDQATAAANQTTFAVGGVMVNNQTGEVISAIHNNVIIPLSNNVSFTFDPTAHGERQLVYWYYANKEALKLPEPNQITVITSLDPCAMCTGALLTAGFNVGVVAIDTYAGINCAQNFQFATLPANLRTKAQKNFGYYASGAANFKPLTRSYVGGPSVAFKNGVVTPANLRDCGTVFTQSVDTVRNTSNSTGLAPSQMSNPAELPSNSAILQAYRAIYKKAFTIKIDNPRLPDAQILTELKAVLADAPNARNAVAFIDPFGNLVLCMADAFNTSPVHAAFMNVTQEYAKTRWDLMNKYAQASTTDNPALYLTHPKYGTFVYLYAPDPDDSITIMSLGAYGSTMEGPIPNMFPSNLQFYYPPRNGAQFSELVPVVNELPPFYTQNVNISLMQVPGVTQAPTK</td></tr>
<tr><td>pYY-BEM4.6</td><td>tr|K1ZCJ4|K1ZCJ4_9BACT</td><td>MSSRAKKNRSTNLKKSIGQKSIENKPTDQKKDQVLVAYVPVIHEGYRRFFRHFPAVKELWLISQELSHELRSLQKDIRALKASETKKLLQTWGQFQKIKLLTPSSLAILQKTTTQLVFPDEEISHHLVEKYFAQNRVLFASFFLRWDKKSSLKKHDLQEYSEISNKEFDQMMIAIAQQEADKSDDWWRQVGGLIFKDETILLLAHNQHTPTEAEAYFAGDPRADFHQGEYLKISTAIHAEAYLIAQAAKQGISLEGADLYVTTFPCPVCAKQVAYSGIKRVFFREGYSLLDGETILKANGVKLIRVTV</td></tr>
<tr><td>pYY-BEM4.7</td><td>tr|A0A1G3PNQ8|A0A1G3PNQ8_9SPIR</td><td>MRDLPLLVLGLTGPMGAGCTRFARDISKMEPGKVIKKQGLLDQVAHEISELSKKASEIRLQCISNGKNSELAELKRLNRRLNAKLAERACLHVIAKSSLPEPLFISLNTIVIKIAVDSITAPEFAEWAKNHAKVADLLKWLRTQWESELTLYETWGQDAGRFSQDELEKMDAMFAEFERIGDEILKEDFETYFGKRNNDFSIRMFSENIRLSGNPFRPAENGGGGGKYDEPSMVMIARETDRYIRFYRTRSDQKRSHFFIIDEIKNPREAEYFRARHQNFFLVSIFSSSEIRASRMRRGLGHDAGVSDADFQHLFRELDSRDWGADDFDAHGLHRQNIYRCFNLADIAINNDVEDERFSEVLFNKFIRYYALMLSPGCVQPTPQETYMHLAYSLSLRSTCISRQVGAVITDLEDRILSLGWNEVPEGQIGCGLKVKKDYTDKENPLFEMEIWDNVITAEDLAVWDDEDSICVKDILSRIEIKTKLKSVSLTPEERADVLKALRIKRLEYSRSLHAEENAILQVASRGGVGLKDGTIYVTTFPCELCSKKIYQVGISKIYYTEPYPNSISEKVILKDGIRNIKILQFEGVKSYSYFKLFKPGFDKKDAQMLEGRGI</td></tr>
</table>

<table>
<tr><td>pYY-BEM4.8</td><td>tr|A0A1G0PGF4|A0A1G0PGF4_9BACT</td><td>MKHNNQLRKEIEKLLGQNSIIKNDELKKLQKEYKIETDELLISFLPYAAEFAKVPISKYKVGAVVLGKSGNIYFGSNMEFEAGALSATVHAEQSAVNNAWLNGETGINKIAVTAAPCGYCRQFLNELTTAKQLHVLLKDKNLEAAKVFKLTELLPEAFGPRDLEIEGGLMKVENHKLKIENINDELINAALEAANKSYAPYSKNYSGVSIQLSDGTIFSGRYSENAAYNPSLLPFQSALAFMNMNTKKGSNNKIVDAVLVEAVSNISQKDAAGTLLNSISKTKLRYYKIKN</td></tr>
<tr><td>pYY-BEM4.9</td><td>tr|A0A0P4WGY5|A0A0P4WGY5_9EUCA</td><td>MEENSSATSQPKCASRTKQGGNDLSTDMSNLSVGETKRTDFLPWDDYFMAVAFLSAMRSKDPSSQVGACIVNADKKIVGIGYNGMPIGCSDDELPWNKESLDPLQTKYMYVCHAEMNAIMNKNSSDLAGCCVYYVALFPCNECAKLVIQAGIREVVFFSDKHQQKPETVASKKMLNMAGVAYRQYTPSQSKIELNLSLKEQEKSEPTADITQSSERDQNSKRKDYLSWEEYFMAMAHLSALRSKDPITQVGACIVNSKKKIVGIGYNGMPLGCNDDLMPWGNSSSNKLETKYMYVCHAGVNAIMNKNSCDVSGCTLYVALFPCNECAKVIIQAGIKTIIYASDTNKDQASILASKKMLDMAGIKYRADNLSQRKIVIDFKTIDWNSRFMNDHQNDPTCL</td></tr>
<tr><td>pYY-BEM4.10</td><td>tr|A0A3D8IG27|A0A3D8IG27_9HELI</td><td>MRKNILYFILTLFFLSGLYATSLPEDNVVSGVIYEKIDTVSAEVDHIYPMLALAIVYKDWQEKNMLNKQGHNIGLVIVDENNMPVFWVRNSVHATHNGTQHGEVRLVSNLLNCEGFNKYLDKYTLYTTLEPCIMCAGMLSMVQIPKVVYAQKDLSCGNTQEIISTAKYPRYYKAFTVENGYKKDLEECFEQYKICKNDSITDFLVNDSAKEIFRKASNDLQDYKVKFKENRRVIKVAQEFLQNIQTKDNLDVLQCPKNM</td></tr>
<tr><td>pYY-BEM4.11</td><td>tr|A0A351C8C4|A0A351C8C4_9BACT</td><td>MNELTKQSEHLRNEALRIATRSYVPYTGQQEGVIILLENGDLIPGVRVENASFQLTIPALQNALSTMYALQRTDISMIVSSIPFTDSDLAYTGGMAEIAWEMVGASLLLVAGAHIPEAGTFIDPARGENLLDVSREAALNAFIPESDFPVGSAIQTSDDVVIDGCNVEHSDWSKIICAERNVLSTARSYGLGQITTIYVSCPKEPGGTPCGACRQVIVELAPDATVWMDRGNQEPIAMKATKLLPGHFTGNVLKKQ</td></tr>
<tr><td>pYY-BEM4.12</td><td>tr|A0A1G6V2K7|A0A1G6V2K7_PEPNI</td><td>MPIVRVNEIGARLPEDWEALETAIWQAYVSREDLPDAGELDLTLVDDATIQELNKTHRQLDKSTDVLSFPMYDDRDDLAADVQAGLPVILGDIMISVPTAERQAQAYGHSFKREMAYLLVHGLLHIAGYDHMSAEEKSAMRRAEEAILADVDVPRDTAPSKTAAVLDEADVQALIDAARAARLQAYAPYSGYAVGAALLAADGRRFCGVNVENASYGATCCAERTALFAAVTAGARDFIALALVTEGDEPAPPCGLCRQALAEFSPDLAIYLAGPTGETYRRTSLAALFPEAFSLSTKESV</td></tr>
<tr><td>pYY-BEM4.13</td><td>tr|F2NP91|F2NP91_MARHT</td><td>MPVMETHALEARFKEALARLCPEGRLLAAVSGGGDSVALLYLLKAAGRDTIVAHLDHALRPDSAADAAFVEKLAQRLGFPLETEHVDVRALAHRKRINLEAAAREVRYAFLARVARRWKARCILTAHTLDDNAETVLLQILRGAGRGLGIRPLQRRVARPLLEFSRAELRAYLEARGARWLEDPTNRSLELDRNYLRHAVLPRITARFPHALEALARFSQAQQADDWALEALSARHLIPDRRWPVPAYRALPLERAPEALRRRAIRGVLEALGVRPEARLVADVEAALGGRAQTLPGGVVVRRQRGTLFFIPPTVRFPKVQPPAGLEARPPRPGDYLVFPYGRKRLVDFLNERGVPRELKRRWPVGAVGAEVRWVYGLWPEPDEDRYMRRALVLARAAARQGEVPIGAVLVRDGAVLAEAANAVEASRDATAHAELLALRTALRRVGEKVLPGATLYVTLEPCPMCYGAILEARVARVVYGVENLKAGAFTVHGLEPRVALEAGRVEGECAKVLKDFFARLRPGRDGA</td></tr>
<tr><td>pYY-BEM4.14</td><td>tr|A0A316TX77|A0A316TX77_9BACT</td><td>MINGYTPYSGNQNTCYVKGESGTFYPGVRIENVSYPLTISSVQAAVCSCLANSDNPVEYYTGDHQPELLQVWADEYDMKPGGKLPDSPLKLFDPLVPSIPDIKKELDVLTEKSVTPNSGFPVSALLQTEKGYIRGVNIELSSWALGLCAERVAISRALTAGYTQFKSIHIYAPEADFVSPCGACRQVLLEVMPDADTELYHGDGTLSKHIVSDLLPFGFTSHKLKK</td></tr>
<tr><td>pYY-BEM4.15</td><td>tr|R6VYG3|R6VYG3_9FIRM</td><td>MIHKGTQTIETKRLILRAFTPDDAEAAFENWMSDPKVTEFLRWKTHADISDSRKIVNEWANGSADPEFYQWAIVPKDVNEPIGTISVVDRNDALGIFHIGYCIGSKWWHKGITSEAFSAVIHFLFEEVGANRIESQHDPENIHSGDVMKKCGLTFEGTLRQADFNNRGIVDACVYSILQSEWQNNTSVWQRLYNAALTVQNDRVVSPFIDAGGVAAALMTKKGNIYTGICIDTASTLGMCAERNAVANMLTNGESRIDKIVAVMPDGKVGAPCGACREYMMQLDRDSGDIEILLDLETEKTVRLKDLIPDWWGAERFGDTE</td></tr>
<tr><td>pYY-BEM4.16</td><td>tr|A0A3C1HZ18|A0A3C1HZ18_9BACI</td><td>MGDIMENWNELSEPWKRCFLQAWKAYCHGSIPIGAVLVDSEGEIFLEGRNRVHELTAPEGQLCDCRIAHAEMNVLVQVKTSDYEKLSGATIYSTMEPCIQCFGAIILSRIKNISFAAIDDKLAGATTLEDRHGFIKSRNLNIAGPFSHLGEIQIILRTDFLLRIFDSEYADPLIAAHEKDYPIGVALGRHYHRNNRLQVAKKETIPFGELFNEFSFDIKRAREGYTLGK</td></tr>
<tr><td>pYY-BEM4.17</td><td>tr|A0A1M6KV24|A0A1M6KV24_9BACT</td><td>MEASQQNILLKIEGKGPVAEINFTVTLPEWLVEQVQSGSTVFLTQKEKMRFVLELARKNVAQETGGPFAAAVFSLESGELVSAGVNVVVESRCSSAHAEVVALSLAQKAVDSHDLGAAGLPRMVLVSSAEPCAMCMGAIPWSGVKQVICGARDEDVRSVGFDEGAKPLEWVEDFAERGIEVIRDVLREEATEVLWDYRERGGEIY</td></tr>
<tr><td>pYY-BEM4.18</td><td>tr|A0A2U0T9B4|A0A2U0T9B4_9RHIZ</td><td>METAELISRLLDVIEKDIAPVTAKGVARGNKLFGAAILKKSDLAVIVAETNNEIENPLWHGEMQAIKRFFELPADQRPATRDCLFLATHEPCSLCLSGITWSGFDNFYYLFSHQDSRDGFAIPYDIQILKSVYAVPEPETGTVSPARDLYNRSNDFWTSHGLQDMIAGLARSNREALLARIDDLNALYAELSERYQRDKGGKGIPLP</td></tr>
<tr><td>pYY-BEM4.19</td><td>tr|A0A2K9PN08|A0A2K9PN08_9FLAO</td><td>MSDKKESKIKISKTSESIELDEIHSLLSYSIVQKFWENDDRNGRGYNVGVILVDENKNIVDWDINSVNKTENSTQHGEMRLISRYLDKDELYSLKGYTMYPTLEPCAMCAGMMTMTNVYRTVNGQMDYFYSKALERLSIDTRECGGYPPYPRTVISEISPSSISTRLDAEYKQYTNAGNKPIITKFLSTYKAKTIYDDAFNQFINFKCKFPENKTKYENAIKFYNSLPESI</td></tr>
<tr><td>pYY-BEM4.20</td><td>tr|F4PWM7|F4PWM7_CAVFA</td><td>MRFSLSLLFVILSVLLAGVLACKDPYNPETVDYGQCASATKANYEVRSDSKVLTPADLPADELAVHESRMRHIIDIARVNNKKFVSSIYFPNGTLACIGINTGKPNMIAHGEIVAIQNCTEIHGISMYTNYSIYTTGEPCSMCASAILWSRFKTVVWSTYNSDLYCKICMSNIPIDSSYIFSRAYGLGIEAPVAIGGVVKAEGDAWFGTYCNRPTSIYYIAPKCACQDPAKVSPLKFTQTRTTVWVEGGDKVVTQWNAIISNPSNSTIVDPPIVISPSVVFKGAPWGISAASEPNTYKLSYNKVLFPGQTFSFGYSVYGLEEVAFTALEA</td></tr>
<tr><td>pYY-BEM4.21</td><td>tr|U7QZM1|U7QZM1_PHOTE</td><td>MNKTRRKLLATLGIMSISMSFIAQAGEKKTQVINNILSKQEITEHEKYMREAIKEAIKNPKHPFGAVIVNRNNGEILSRGVNTGRNNPILHGEIQAINHYITQYGNQGWENVALYTTGEPCSMCMSALVWIGIREVIWATSISVIRNSGIRQIDISAHEIAERASSFYNPITLVGGILANETDKLFLERKRGN</td></tr>
<tr><td>pYY-BEM4.22</td><td>tr|A0A081CH48|A0A081CH48_PSEA2</td><td>MASRRHLLATQVTGNHRKLSLWHLRGWLSPYTKLVDAVYFLTTNSFYHSLQTPPVQSITMLLSSIITSLALAAQASAYREGLHPEFQSGLSINSVPATDRDHWMRLANSAIYYPPVSHPCPQAPFGTAIVNTTSNELICAIANRVGSTGDPTQHGEITAIQHCTNVMRKKGLSPQEIIAAWKQLSLYTNAEPCTMCLSAIRWAGFKEVIYGTSVGTISENGRNQIYIPSNLVLEKSYSFGHATLMLGNILTHETDPFFQHQFNESAPCPVGCERTQVGEARVKTCEPVPNWQKLVRLEYSEDSRVGSEPVAHTPLHLEL</td></tr>
<tr><td>pYY-BEM4.23</td><td>tr|A0A3D3HMU1|A0A3D3HMU1_9GAMM</td><td>MDYSDAILGAITSIRRNSKQPGVNVTDNVTDSSTQYNNDEYWMRRALALAREAGEAGEIPVGAVLVKDNQQVAGGFNQPIRSHDPAAHAEILTLREAGAVLGNYRLIDTTLYVTLEPCMMCAGALVHSRIKRLVFGAAEPKTGAAGSFIDLLTLPRLNHYMEVTGGVLGEECSVLLSDFFRRRRAEKKALKRQNSESGSDSAS</td></tr>
</table>

| | | |
|---|---|---|
| pYY-BEM4.24 | tr\|A0A1N5WT13\|A0A1N5WT13_9ACTN | MLEKIERRLVAAAEAVVRSPSTGDAHTVAAAAMDANGDIYSGVNVFHFTGGPCAELVVIGSAAAANAPPLITIVAVGDGDRGVIAPCGRCRQVMLDLHPDVFVIVPTGDGQLAAKPVRELLPFGYVARTGSTAPRVVYFHPRHYDTISSGLKTATVRFQDSVQTGPAVFVFDDGESIRRLDAVVEKVESRRLDHLTEEDAHHEALPDSDALRDAIKTQYPMLGDGDVVDVATFRLTAISAPDPDPRSSYPPAVSRCNPAGPRADLLVGQS |
| pYY-BEM4.25 | tr\|X0SAC5\|X0SAC5_9ZZZZ | MTKDGRVIASAHDTEVTDQDSTAHAEINAIRKASKIYRKDLTGCLIISTHEPCPMCTGSIIWSNISKVVYGVSIRDSIKAGRDMINLSCKEIIKKPNAEINIYDGILKKECLKLYNNDTRKLVKKFRKYEWINIEENLLNKRMQWFENNKTMIRKLKGNDLEKAYHLILMKIGIKRSEAPIVKKSESKIIFHSKNYCPSLEACIILDLDTREVCKEIYERPTEELIRRLNSKLRFTRNYDCIRPYSDYCEEIIILEK |
| pYY-BEM4.26 | tr\|A0A3B8IC10\|A0A3B8IC10_9BACT | MPSHEDFIHQCLELGKEALLQGNPPVGSVIVWQDQVIGRGIENGRSSGDITQHAELLALQEAVATGQRDKLKEAIIYSTHEPCVMCAYPIRQYKIPTVVYSVAVPELGGHTSSWHLLTTEDVPKWGKAPKIITGISAEEVEALNAAFQDSLKKG |
| pYY-BEM4.27 | tr\|A0A2N9P8B9\|A0A2N9P8B9_9FLAO | MFIFKLISPPVSIEVYQDKIIQKLYICFMENIFTDEYFMKKALQEAETAFQQGEIPVGAVIVIDNRIIARSHNLTEMLNDVTAHAEMQAITASANFLGGKYLKDCTLYVTLEPCQMCAGALYWSQISKIVYGATDEQRGYRAMGAQLHPKTKVISGIMQNECTHLMKDFFKQRRSKSTKD |
| pYY-BEM4.28 | tr\|K1KX30\|K1KX30_9BACT | MVKNPVNNNELYFGKHSEIPMNEEQKAYMKMAVDLSRSGMESGKGGPFGCVIVKDGKVIGIGSNSVLETNDPTAHAEIVAIRDACRNLGHFQLDGCEVYTSCEPCPMCLGAIYWARPSKVFFANDKRDAAEAGFDDDFIYQELELPYEKRKIPFEQGMQDTAKEVFQEWILKEDKTLY |
| pYY-BEM4.29 | tr\|R4XI84\|R4XI84_TAPDE | MSSEIEPPSTDVHKHAVAEAADESGAADAFMQIALQQAETALLNKEVPVGCVFVHQPTGTVLATGANQTNASLNGTLHAEFVAIESILRDHPPSIFRESDLYVTVEPCVMCASALRQLQVRKVYFGCGNDRFGGCGSVFSIHSDASKTGDAAYMVESGIFRKEAIMLLRRFYLLQNESAPKPALKSTRVLKEHFDE |
| pYY-BEM4.30 | tr\|A0A239CVF7\|A0A239CVF7_9DELT | MSPASKKHFPSLFSLFLLLTIGLICGTAHAQPQGHTADDTAATLANASLKEHEPFIRRCYQLAIDAGKKGNHPFGALLVHKGKIVLEAENTVLTDNDFTNHAEMNLIAEAARTLSRQIIPEATVYTSCAPCAMCTATLAMAGFTRIVYGVSHDALNKRFGLKGKSVSCPALFKTMGMELEFVGPVLEKEGLRVFDFWPEKDPHAQMLKKQARK |
| pYY-BEM4.31 | tr\|A0A1Q3NME1\|A0A1Q3NME1_9BACT | MTEFNYDWAKLAFSSKRPLTNLKATFIIAPREISEKRFTQLLKEYLPKGDILLGISKEDYVEGLEGQPQFAMLQQKTLQKLIDKVNDASAHKVYTLRYFQRELPAIIEKLTPPRVVGIHGSWHHSFHTLPIYYLLSEKRIPYQLVAAFSDEDEARAYEVATDKKIVRPTLEGSFDDTTVLQLTDEVAKSSYDYGFQTGAILAEKVNGVYQPVAAGFNKVVPYQTYALLNGASRETNFSPANDMNHYDTIHAEMQILVEAAKQGISLKDKTLFVNLMPCPSCARTLSQTELSEIVYRIDHSGGYAVDLLTKVGKDIRRIVY |
| pYY-BEM4.32 | tr\|A0A2G6N4N7\|A0A2G6N4N7_9DELT | MKERTVSYSDRHFMAEALEMAESALTQGEFPVGCVIADGTAVVARGHRTGTTAGAVNEIDHAEINALRHLGLAGEHLDRTDLTIYSTMEPCLMCFAAIVLSGINRIVYAYEDVMGGGTGCDLTGLPPLYRDAPLTLVAGVRRRASLNLFRRFFTDPENGYWAGSLLSRYTLNQTKDSHRL |
| pYY-BEM4.33 | tr\|A0A0G0RBB8\|A0A0G0RBB8_9BACT | MQSVQYNKLTHLQRRALDEAEQVLENSYNPYSHFYVGACLISEDEQLIAGTNFENAAYGSAICAERAAVLRANAMSIRRFRGIAIIARGEDFNTTEVTGPCGSCRQVLYEISQVSGCDLQVILATSKKDKIVITTIRELLPLAFGPLDLGVDIGKY |
| pYY-BEM4.34 | tr\|A0A327L2Q5\|A0A327L2Q5_9RHIZ | MVTSRDGEDEAMMARCVALSRIAVGKGEYPFGAVVAREGRIVAEAINRTIRDGDVSRHAEVIALARAQKAIGRRELRECSLYSNVEPCAMCSYCIREAWVGRVVYALGSPVMGGVSKWNILRDDGLSGRMPQVFDAAPEVVSGVLVEQAQAAWRDWSPLAWEMITLRGLMTDPSARPECRTRAARPRSLWHHLVALIERPPRPYVDPTSAAEGHADL |
| pYY-BEM4.35 | tr\|S2DR30\|S2DR30_9BACT | MKMKKKIEITVSLEVIQKSEWSKEDRSLIERAIHAVEHAHAPYSNFMVGTALLLDNGQIFSANNQENVSFPVGICAERAVLSYAMGNFPNNRPVKLAVVAKRRSDSTWATVTPCGLCRQTINEYEVKFGHPIEILMLNPGEEILKASGIDQLLPFRFNDLNS |
| pYY-BEM4.36 | tr\|A0A369QGF1\|A0A369QGF1_9BACT | MEEHEKWMHWCLNLAQQALQQGDFPVGAVVVQKGKLIGQGVEAGQLKKDITCHAEMEAIRDARQTINTADLQNCILYSTHEPCIMCSYVIRHHKISRVVVGTTVPEVGGSSSAYPLLSAPDISIWVAPPHLVTGVLAEACQALSQAYKQKFKK |
| pYY-BEM4.37 | tr\|A0A1W6X4U4\|A0A1W6X4U4_9RHIZ | MTNPSRQERWDRRFLELAKVFGTWSKDRSAGTGCVIVGPDRLLRASGYNGFARGIDDEVPERHERPAKYSWTEHAERNAIYNAAKLGISLDGCTAYVNWFPCIDCARAIVQAGIVRLVGLHPDHADQRWGSEFKFATEMLRESGIEIILYDIPELAARK |
| pYY-BEM4.38 | tr\|A0A238BW09\|A0A238BW09_9BILA | MEEMARKIRTKAKKANSYCNTMTFLISKASIVLLKAECKRIELTVVIFRFLIKMNASEPNNELCDMTVIKSMLKITHVIFDLDGLLIDTEVVFSKVNQCLLSKYNKKFTPHLRGLVTGMPKKAAVTYILEHEKLSAKVDVDEYCKKYDEMAEEMLPKCSLMPGVMKLVRHLKTHSIPMAICTGATKKEFEIKTRYHKELLDLISLRVLSGDDPAVKRGKPAPDPFLVTMDRFKQKPEKAENVLVFEDAANGVCAAIAAGMNVIMVPDLTYMKIPEGLQNKINSFSDNLIISNDLNVALMSLKKELSEEEVHFLNRAFEIAVDAVLNNEVPVGCVFVFEGQEVAFGRNDVNRTKNPTYHAEMVALKMMKQWCMDNGRDLEEIMRRTTLYVTLEPCIMCASALYHLRLKKILYGAANERFGGLVSVGTREKYGAKHFIEIMPNLSVDRAVKLLKEFYEKQNPFCPEEKRKVKKPKKSGNNNDNSDDAVALNV |
| pYY-BEM4.39 | tr\|A0A1J5H6Z0\|A0A1J5H6Z0_9BACT | MAYQPSEKFMQMAIDKTREGVLSGQTPFGACIVKDGKVVACEHNTVWQDTDITSHGEVHTIRAACKAIGSIDLSGCILYSTCEPCPMCFSAIHWARIDTVVYGAFIADAQDAGFNELTISNEKMKEFGGSPVNFISGFMRDENVALFKLWKEQGANNVY |
| pYY-BEM4.40 | tr\|A0A3C2D945\|A0A3C2D945_9BACT | MKTTEIRIIVHEYQNIDELTENDQYLLHEARRITEFAYAPYSGFHVGAAILLGNGMIVKGNNQENSAYPSGLCAERVALFYANANYPDSEVKTIAISAAKNGILVNDPIKPCGGCRQTLSEAEVRFGSPIRIILDGQDSILVLHGVESLLPLSFSKKDLASPLAATGR |
| pYY-BEM4.41 | tr\|A0A1I7EYS3\|A0A1I7EYS3_9BURK | MKFKLDPSRPPDEDDYYLGVALAVRRKANCTGNRVAAVIVKNKRVIATGYNGVPEDMPNCLDGGCLRCSNPGGQFKSGTRYDLCICVHAEQNALLTAARFGISVEGAHLYTTMQPCFGCAKEILQAKIEKVFYLHPWVPTDVDPVMDAAMKAEYAKIIGKLKVKKLDFDDPVATWAVTTMRQAALASDKNPDKKTPPKTAKKKVAKKKSRTSPR |
| pYY-BEM4.42 | tr\|H8GQX8\|H8GQX8_METAL | MNHEHFMRRAIELARQAPQYPFGAVIVRRDDGQCVGQGFNRSDLNPTYHGEMVAINDCAVRHCAEDWRGFDLYTTAEPCAMCQGAIEWAGIGRVFYGTSIPYLQKLGWWQIDLRAAEVSARAVFRDTLIVGGILETECNALFAAARRGCFGTGSE |

| | | |
|---|---|---|
| pYY-BEM4.43 | tr\|A0A0S8HZN3\|A0A0S8HZN3_9CHLR | MDEHDIRFLRASFDVARNARKNGNHPFGALLVDEHGRIVMEAENTVITAKDCTGHAETNLMREASSKYDSDFLANCTIYTSTEPCPMCAGAIFWSNVRRVVYGLSEESLYEIAGRGSEEVLFLSCREIFERGKKLIEVIGPLLEDEAREVHMGFWR |
| pYY-BEM4.44 | tr\|E3SF31\|E3SF31_9CAUD | MKPTTVLQIAYLVSQESKCCSWKVGAVIEKNGRIISTGYNGSPAGGVNCCEHAEEQGWLLNKPKPVLIPGHKSECVRFSQVDRFVLAKAHREAHSAWSKNNEIHAELNAILFAARMGSSIEGATMYVTLSPCPDCAKAISQSGIKKLVYCETYDKNIPGWDDILKNAGIEVFNVPKRSLDKLNWENINEFCGE |
| pYY-BEM4.45 | tr\|F8AAC6\|F8AAC6_THEID | MIRAPWHEYFMLLAKIVALRSGCNSRPSGAVIVKNKRILATGYNGPMPGAWHCTDRGPGYCFRREKGIPDIDKYNFCRATHAEANAIAQAARFGISVEGASLYCTLAPCYVCLKLIASAGIKKVYYEHDYGSRDFERDQFWKEAIKEAGLEKFEQITVSQEVMEQLQEILPYPTSKRRLAPTEFLDEFEDGKKYGVPSIEVLFNKLNYLTRQALKDITFVIEKTTVTEEPEGISFYLSGKMVELSELINTVKKQINADQNFYFLAKHNAIEAKIEILREAENIRLKAFLNECPLESFKRIAESLDYILYQVSNSLSLPTRLELSVNLLRI |
| pYY-BEM4.46 | tr\|A0A2H4ZNK4\|A0A2H4ZNK4_9EUKA | MKKQLSRKIQEEWMSRLLRNAYDAGTYGEVPIAAVILNESGQCIGWGRNCREKDQNPLGHAEIIALRQASYLKKSWRFNECTMLVTLEPCPMCAGALLQARINHIIYGASDYKRGGFGGVLDLSKNSSAHHKIEITRGVKSIQSCQLLETWFRRRRRV |
| pYY-BEM4.47 | tr\|A0A239N5N1\|A0A239N5N1_9PSED | MEGRAGIIPFDEGGAAMGPAEEDSPMQHLAYMREALALARANVEAGGRPFGAVLVRDGEVIARAANGTHLDHDPTAHAELLALRAAGRALGSPRLDGCVVYASGHPCPMCLAAMHLSGVSAAYYAYSNADGEPYGLSTAAVYAQMAQPVEWQSLPLQALRPEDEEGLYGFWRERRP |
| pYY-BEM4.48 | tr\|A0A328VTR2\|A0A328VTR2_9PSED | MHPEHLALLQQAPASTHADDTWARLCCEQALLAVEEGCYAVGALLVDGAGELLCSGRNQVFAPAYASAAHAEMRVLDQLEAEHAQVDRRSLTLYVSLEPCLMCYGRILLAGITRVRYLARDRDGGFALRHGRLPPAWANLASGLSVVQAKADPYWLDLAEHAIGRLQDRQTLRQRVIRAWRGQRTLTDEFSSTKRTHSG |
| pYY-BEM4.49 | tr\|A0A103YG48\|A0A103YG48_CYNCS | YIRELHASSLRRDEHEIQNPKILVIVDRLSSPSLHVSLSLSLSLVIFPPFIPLNQTPTHMENAKVVEAKDGTIAVASAFSGHQEVVQDRDHKFLTRAVEEAYKGVECGDGGPFGAVVVHKDEVVASCHNMVLKHTDPTAHAEVTAIREACKKLNKIELSDCEIYASCEPCPMCFGAIHLSRIKRLIYGAKAEAAIAIGFDDFIADALRGTGFYQKAHLEIKQADGNGAMIAEQVFEKTKAKFAIDHKFLTRAVEEAYKGVECGDGRPFGALVVHKDEVVVSCHNMVLNYTDPTAHAEITAIREACKKLNRIELSDCEMYSSCEPCPMCFGAIQISRIKRLVYGAKAEASIASGIPIGDFISDALKGTGFHEKANFEIKQADGNGAMIAEQVFERTKAMFPKR |
| pYY-BEM4.50 | tr\|W5M1M8\|W5M1M8_LEPOC | NSSTRESRVMAQMEINGGASPPKKPGKGQSAADQDMITGLINKALQAKEFAYCPYSNFRVGAALMTNDGRVFTGCNVENACYNLGVCAERTAILKAVSEGYESFRAIAVSSDLQDQFISPCGACRQVMREFGTGWDVFLTKVDGSYVRMTVDELLPMSFGPDDLKKKKVFSLQNGHEVSTQFYTHSPCEAGENNN |
| pYY-BEM4.51 | tr\|A0A3N5YPZ2\|A0A3N5YPZ2_9ALTE | MSNSETEHIQALVDAAQAAQKQSYSPYSSFQVGAAIFADDGNTYSGCNIENVAYPLGQCAEATAIGMMIMQGAKRIEDIMIASPNDQVCPPCGGCRQKISEFGTAETKIHMVTRSGEVSTVTLGELLPLAFDSL |
| pYY-BEM4.52 | tr\|A0A2A9NC86\|A0A2A9NC86_9AGAR | MTNSTLSNEDRTRLIQGAFQARKKTYSPYSNFPVGAALLTTDGRIIEGANIENASYGGTICAERTAIVKAVSDGYRHFAGIAVTTKMPTRVSPCGICRQVLREFCSLDMPVLLVPGDYPQRNPVDDDGADKPGVITEGGVRETTLGALLPDSFGPENLPPRA |
| pYY-BEM4.53 | tr\|A0A2D6RD43\|A0A2D6RD43_9GAMM | MNIENLITENDETLIRRCIELAGESVKNGDKPFGALLAKDGNIIFESSNNAKTKVPYHAEILTLMDAQDKLNTTDLSDYALYSNCEPCPMCSFMIREYKLDKVVFSVHSPYMGGQSRWNILEDDVLTRFKPYFSKPPNVVGGVLESEGKRIFDKVGLWMFGKE |
| pYY-BEM4.54 | tr\|A0A0H3AVL6\|A0A0H3AVL6_BRUO2 | MHAKGYSQQERRIIPFANRFRFRELCSNKSLHGLRAKFPEQYTKWDPMRKAASITKANSATPMDIALEEAHAAGERGEVPIGAVIVRDGEIIARAGNRTREFNDVTAHAEILTIRQAGEMLGSERLIDCDLYVTLEPCAMCAAAISFARIRRLYYGASDPKGGGIEHGGRFYTQPTCHHAPEIYPGFCEADARKILKDFFREKR |
| pYY-BEM4.55 | tr\|A0A242H531\|A0A242H531_9ENTE | MFIVKNNIEVIQQQAELDAKFMKQALKLAKDASNNGNEPFGAVLVKNDKVILTGENQIHTESDPTYHAELGIIRDFCTSQKITDLSEYTLYTSCEPCCMCAGAMVWSNLDRMVYGLGHDELAEIAGFNIMIGSEEIFSKSPNRPEVAKGVLKEAAVPVYVDYFQR |
| pYY-BEM4.56 | tr\|A0A2R6XZE2\|A0A2R6XZE2_9BACL | MSGRISWHEYFMAQAKLIALRATCTRLMVGAVIVRDRRVIAGGYNGSIAGDEHCIDVGCKVRDGHCIRTIHAEQNALMQCAKFGVSTDGAELYVTHFPCLNCTKLLIQAGIRHIYYEVPYRVDPYAIELLEKAGVGTTQITVDLNAYVQVMSKVSTDPALTYVPESKAQKDEYGQSVGKIV |
| pYY-BEM4.57 | tr\|A0A139SHT6\|A0A139SHT6_9BACT | MSEANASSESLPSRNSPVELIAEAAGKFGRRPTWDEYFMATAVLISTRSSCERLNVGCVIVTAGESHKNRIVAAGYNGHLPGSPHTSRMRDGHEQATVHAEQNAISDAARRGSSVEGCTAYVTHYPCINCAKILASAGIAKICYRLDYHNDPLVKPMLAEAGIEIVQLGEAAS |
| pYY-BEM4.58 | tr\|A0A261DBH2\|A0A261DBH2_9RICK | MVMKKKLITVKRSTEFNNFFMEEALKQAQFALDKNEIPVGAIIVNRITNKVIAKAHNIVEQTKNPVLHAEIVAINQSCQILSSKNLSDCDMYVTLEPCVMCSGAISFARIGRLFYAANDPKQGGAIENGGRFFNSKSCFYRPEIYSGFSAKISENLIKEFFYNVRYQKCNP |
| pYY-BEM4.59 | tr\|A0A2N0XZK6\|A0A2N0XZK6_9VIBR | MTDNSLHESYMRQAFELSKSALPGCRPNPPVGCVFVKDGEVVSSGFSQPPGNHHAEAGAIAAYTGSYDGLVAYVTLEPCSFQGRTPSCAKALVRVRPEKVYVAILDPDTRNSGAGIKILEDAGIDVEVGLLGEEVASFLNPYLIRN |
| pYY-BEM4.60 | tr\|A0A1V5R0F9\|A0A1V5R0F9_9BACT | MTKKETTKLHALDDFCMKKALLLAKRAFRADEVPVGALVVDSSNKVIGRGYNQVEKRKSQRAHAEQLAIEQACKKIGDWRLEGCTLYVTLEPCTMCMGLIKLSRIERVVFGAASPLFGYQLDKNRKSQLYKKGVIKIRKGVGKATAAALLKDFFKNKRM |
| pYY-BEM4.61 | tr\|A0A2W0H8Y3\|A0A2W0H8Y3_9BACI | MKNNGRLDHEYFMTEALQEAKEAGQRGDLPIGAVIVHNGRIIARGSNMRKTAGIKISHAENNAMHNCAPYLMKHASECVIYTTLEPCIMCLTTLVMANIDSIVFAADDKYMNMKPFIDANSYIRDRIHQYKGGVCRGESEALLRKYSPYAAELALNGTHPHHRKGGA |
| pYY-BEM4.62 | tr\|A0A261BDB7\|A0A261BDB7_CAERE | LYKLYIFRMTTTKANLTQFEQELVDKAVGAMEKAYCKYSGFKVGAALVCEDGEIIIGANHENASYGATICAERSAMVTALTKGHRKFKLLAVATELEAPCSPCGICRQYLIEFGDYKVILGSSTSDQIIETTTYGLLPYAFTPKSLDDHEKEAEERNHQEGEKKH |
| pYY-BEM4.63 | tr\|A0A2E1PHI6\|A0A2E1PHI6_9GAMM | MKELLIHSWLMLNSNSKLIMERVIELSEINLKNGKIPIAAVIVDKKNYEIISESQNEDSPIGHAELLAITKALKKLNTNRLDSTNLFVTIEPCPMCAYAISKCHINRLYFGSEDEKGGGVINGPRIFESHNLKKIDYVSHCYHEKTTQLMQSFFQLKRNQQL |

<table>
<tr><td>pYY-BEM4.64</td><td>tr|A0A378LUA7|A0A378LUA7_9GAMM</td><td>MDTIIKKMISNAHNTLAHSYSPYSKFSVASCICTDKDNFYTGVNVENSAYGLAICAETSAISAMVTAGEKRIKSMVVMAGTNILCSPCGACRQRIYEFSTPDTLIHLCDKNSILRTFKINELLPEAFKFDFNP</td></tr>
<tr><td>pYY-BEM4.65</td><td>tr|A0A139HQ78|A0A139HQ78_9PEZI</td><td>MADSLKSKPGHARHDTALIHGLSQSDVQKLSESCVDAKSKAYCPYSHFRVGCAVLLANGDVVQGANVENAAYPVGTCAERVALGTAVGAKKGDFRALAVSTDISPPASPCGMCRQFIREFCELNTPILMYDKDGKSVVMTLEQLLPMSFGPDKLLPPGQLENGLMQTQTQSSFVTRAFSTTSSRRQDDTPQVPQSHYDFFPQTFPQGPPPKTSFSPDLKQLRKEFLQLQAKAHPDLAPQDQKRRAEALSMRINEAYKTLQSPLRRAQYLLSQQGIDVEDETAKLDDSSLLMEVMEAREAVEEVEDEEQLNEIRAENNGRIEESVRVLEDAFRDNEFEKAAQEAIRLRYWVNIEESIQGWEKGNGGGILHH</td></tr>
<tr><td>pYY-BEM4.66</td><td>tr|A0A2A9FXV0|A0A2A9FXV0_9VIBR</td><td>MCNLKENKDMDKYFHFACDATIEGMREGTGGPFGATLTRNGEVVCSVANTVLKDMDISGHAEMVAVREACKKLDTLDLSDCVMYATCEPCPMCVSVMLWAGIKTCYYASTHLDAAKHGFSDQQLRDYLDGSDTSTLNMVHIEDNRDDCAKIWTEFRHLNETKNDG</td></tr>
<tr><td>pYY-BEM4.67</td><td>tr|A0A1A8AG96|A0A1A8AG96_NOTFU</td><td>MEHSDRWSRAEPGLSTSSRETRDGSTQTDCKLQGHGPRLSKVNLFTLLSLWMELFPQEQDEENGQSQIRRSGLVVVREGKVVGLHCSGADLHAGQAAILQHGASLANCQLFFSRRPCATCLKMIINAGVRQITFWPGDPEISMLTSNQTHSQRTSQSITEASLDATAVEKLKSNSRPQICVLMQPLAPGVLQFVDETSRRSDFMERMMDDDPELDSEKLFNSDRLRHLKDFCRHFLIQTDQRHKDILSQMGLKNFCVEPYFSNLRSNMTELVEVLAAVAAGMPQQHYGFYREESLSLDPHPVDVSQAVARHCIVQARLLSYRTEDPKVGVGAVIWAKGQSACCCGTGRLYLIGCGYNAYPAGSKYAEYPQMDNKQEDRERRKYRYIVHAEQNALTFRTRDIKPDECSMLFVTKCPCDECIPLIRGAGVKHIYTSDQDRDKDKGDISYLRFGSLKGVCKFIWQRSPPVSSASSLHLTNGCVGKHVRQAEQQIYKNKKLCTKGSSGSSDIC</td></tr>
<tr><td>pYY-BEM4.68</td><td>tr|A0A3E2VN88|A0A3E2VN88_9FIRM</td><td>MEKEITNMDKQKLIQMAVDGLGRSYAPYSHFHVSAALLCADGTVYTGNNIENAAYTPSVCAERCAIFKAVGDGRREFEAIAVCGGPDGVIEDYCPPCGVCRQVMREFCDPSSFRVLVAKTAEDYREYTLEQLLPDGFGPDHLTGSGER</td></tr>
<tr><td>pYY-BEM4.69</td><td>tr|A0A2D5ZRJ2|A0A2D5ZRJ2_9BACT</td><td>MARPVHLHTGERRTEEGATESRAVAAVATAITRAPRAPPRPATGRERDGPPPRRVFGGGLRVGDPSGYDRGESKPIGGPLTEKRSDWHSYFMRIAGEVATRATCDRKHVGAVIVRNRTILSTGYNGSIRGMPHCDDVGHDMVDGHCIATIHAEANAILQAARNGVMIQDGSIYITASPCWNCFKLVANAGLKRVYYGEFYRDKRSFEVARRLGIDLMHIEV</td></tr>
<tr><td>pYY-BEM4.70</td><td>tr|A0A1B8WPS3|A0A1B8WPS3_9BACI</td><td>MEGVQLIYQFQWGNLIMTVNKEDLYLIDVARNTIKTLYVDGKHHVGAAVRTKTGKIYSAVHLEANIGRVSVCAEAIALGKAISEGESEFDTIVAVRHPDPTQENQKIEVVSPCGICRELISDYGKGTNVILKNKEGYIKTVISDLLPNKYIREDN</td></tr>
<tr><td>pYY-BEM4.71</td><td>tr|A0A1W5ZQK9|A0A1W5ZQK9_9BACI</td><td>MNRFMERAVSLAAENVRVGGQPFGAVLVKDDELVAEGVNEMHLNYDVSGHAELLAIRRAQGELQTHDLSGYTMYASGEPCPMCLSAMYFAGIKDVFYCATVEEAAQVGLEKSKNVYDDLQKSKGERSLVMKQMPLEDDQEDPMKLWDERTNHNGTS</td></tr>
<tr><td>pYY-BEM4.72</td><td>tr|A0A378V0W4|A0A378V0W4_MYCFO</td><td>MVHAQFDPTARQALAATAVEAKTRKDLTWQQIADAAELSPAFVTAAVLGQHALPARSAEAVAALLGLDDDAALLLQTIPIRGSIPGGIPTDPTIYRFYEMLQVYGTTLKALVHEQFGDGIISAINFKLDVRKVADPEGGERAVITLDGKYLPPNPFDRVRYRGGLMDFAQRTIDIARQNVAEGGRPFATVIVKNGEILAESPNLVAQTHDPTAHAEILAIRKACTRIGTEHLIGATIYVLAQPCPMCLGSLYYCSPDEVVFLTTRDAYEPHYVDDRKYFELNMFYDEFAKPWDQRRLPMRYEPRDAAVDVYKLWQERNGGERRVPGAPTSTRPGKNPRGE</td></tr>
<tr><td>pYY-BEM4.73</td><td>tr|I3XF03|I3XF03_RHIFR</td><td>MKQRCMSPKSAQRFWDNDMHNNKDRPMSENELFVAAREAMAKAHAPYSKFPVGAAIRAEDGQIYTGANIENLSFPEGWCAETTAISHMVMAGQRKIMEVAVIAEKLALCPPCGGCRQRLAEFSGASTRIYLCDETGIKKSLALSDLLPHSFETEILG</td></tr>
<tr><td>pYY-BEM4.74</td><td>tr|F8IEF3|F8IEF3_ALIAT</td><td>MDAKELETRGWLCMRAVDVIDKKRRGEALAEEELRFLIEGYVAGRIPDYQMSAFLMAVVWRGMTREETLVLTRLLADSGERLDLSGIPGVKVDKHSTGGVGDKATLVVLPLVASIGVPVIKMSGRGLGHTGGTIDKLESIPGFRTDLSVAELVAQVRQVGIALGGQTADLAPADKKLYALRDVTGTVESLPLIASSVMSKKLAGGADAIVLDVKVGDGAFMKSRSDARRLARLMVEIGEAAGRRTVAVLSNMDQPLGCAIGNALEVAEAIRVLSGEGPFDLAEIALALAEEMTVLAGVAATREEARRMLRQSVAEGRALETLRRWIAAQGGDPAVVDDPSRLPQAPVQMPYLPKKAGFVAKLSALAFGLAAMRLGAGRETKEEAIDPSVGIVLHAKVGDRVQTHRPMFTVHARTGEDALRCIQELEAAIQISDDPVEAPPLILARIDRSEALPYADLMDAAREARDRAYVPYSGFAVGAALELADGRMVTGANVENASYGLTNCAERSAVFRAVAEGGPGTKPEIRAVAVIADSPEPVSPCGACRQVLAEFCSPDTPVYLGNLQGDVRETTVGALLPGAFTDAQMANVRRQDKEA</td></tr>
<tr><td>pYY-BEM4.75</td><td>tr|A0A1G3M638|A0A1G3M638_9SPIR</td><td>MKTTNINALDKWDLRFLQMAEHVAEWSKDPSTKVGAVIVRPDRTIASVGFNGFARGVRDTVERLWNRELKYPLTVHAELNAILSAHEPVRGHSLYVSPLSPCSNCAGVIIQSGIARVVAKCGQVNNPAQWSESFNLALTAFAEAGVSVILVEH</td></tr>
<tr><td>pYY-BEM4.76</td><td>tr|A0A3D9LFR2|A0A3D9LFR2_9MICC</td><td>MEQNDHGSSGAFSDPFEDDIPLTASLPRITGTGSGIDWQRLESTARAAMTRAYVPYSRFPVGAAALVEDGRVVAGCNIENASLGLTLCAECSLVSNLQMSGGGRIVAFYCVDGNGEVLMPCGRCRQLLYEFHAPGMRLMGPDGELTMDEVLPLAFGPADMTHLSDSAASTDDPGRTR</td></tr>
<tr><td>pYY-BEM4.77</td><td>tr|A0A3B9YGB5|A0A3B9YGB5_9BACT</td><td>MAKPISKKYRKLIETAKAARKKAYSPYSRYQVGAAVLTESGRIYSGANMENASYGLCMCAERVAIANAVTRGEKVLQAVCVVGKKARPCGACRQVMLEFSTKETELLMVDIDPNARRDTVIRTRVYSMLPNPFDPFESGMLPQHPQNLLRRRKSPQPRRKRRSRPVHREVSR</td></tr>
<tr><td>pYY-BEM4.78</td><td>tr|A0A182F569|A0A182F569_ANOAL</td><td>MPRPSQFRVSSSQSLSNSQIQASQSSDSVVDITSYVNAVVKALLNLSCTKTIIKRADLVNIALKGNGRLIGRVLQDANIELKEIYGYELIEVEKSKTMILCSTLAAGSMDELNDANRRRYTFLYLILGYIFMKNGSVPETIVWEFLETLGIEEQQEHNYFGDVRKLYDSLFKQAYLTRTKQALEGLNDDVMLISWGVRSKHEVSKKDILAGFCKVMNRDPVDFKAQYIEANEKDDKMNNNINGTVDGRNTVEYSSLDASVKELIEAAIKVRNNAYCPYSNFAVGAALRTVGGDIVTGCNVENGTFGPSVCAERTAVCKAVSEGHREFTAVAVVAFQETEFTAPCGTCRQTLSEFSRKDIPIYLVKPSPVRVMVTSLFQLLPHAFSPSFLNK</td></tr>
<tr><td>pYY-BEM4.79</td><td>tr|A0A264Z0D4|A0A264Z0D4_9BACI</td><td>MEPKKLIEEAIVASKQAYVQYSNFHVGAALLTKDGKLYHGCNIENASYGLTNCAERTAIFKAVSEGEKEFQAIAVVGDTEGPISPCGACRQVLAEFFSPDTVVILANLKGDHVVTNINELLPGFFSSKDLQKKVKNCFEKNALGSSCLRPI</td></tr>
<tr><td>pYY-BEM4.80</td><td>tr|A0A1L9Q1R3|A0A1L9Q1R3_ASPVE</td><td>MPLSAEEAALVETATATINSIPLSEDYSVASAAKASDGRVFTGVNVYHFTGGPCAELVVLGVAAAAGAAQLTHIVAVANEQRGILSPCGRCRQVLLDLQPNIQVIVGKEGSEQSVPVAQLLPFSYRQPDQHTPVIFKALTSSGPVVVDFFATWCGPCKAVAPVVGKLSETYTDVRFIQVDVDKARSISQEHDIRAMPTFVLYKDGKLLDKRVVGGNMKELEEQIKAIIA</td></tr>
</table>

**Supplementary Table 2.** Sequences of sgRNAs used in this study. Target sites for guided off-target[1]and targeted RNA-seq[2] are the same as previous publication and not listed.

*S. pyogenes* SgRNA scaffold*:*
GUUUUAGAGCUAGAAAUAGCAAGUUAAAAUAAGGCUAGUCCGUUAUCAACUUGAAAAAGUGGCACCGAGUCGGUGC

*S. aureus* SgRNA scaffold*:*
GUUUUAGUACUCUGUAAUGAAAAUUACAGAAUCUACUAAAACAAGGCAAAAUGCCGUGUUUAUCUCGUCAACUUGUUGGCGAGA

| site | protospacer sequence | PAM | Cas9 scaffold | PAM | Cas9 scaffold |
|---|---|---|---|---|---|
| 1 | GAUGUGUCUACUGUUACUUACA | AGGAAT | S. aureus | AGG | S. pyogenes |
| 2 | GCACCCAGGGGUUCUGCAGAGC | AGGGAT | S. aureus | AGG | S. pyogenes |
| 3 | GCAUUCCACUCCGUCCGCCUC | CGGAGT | S. aureus | CGG | S. pyogenes |
| 4 | GCCACAGACUUUUCCAUUUGC | AGGAGT | S. aureus | AGG | S. pyogenes |
| 5 | GCCACAGUGGGAGGGGACAUG | GGGAAT | S. aureus | GGG | S. pyogenes |
| 6 | GCCCAGCAAUUCACUGUGAAG | AGGGAT | S. aureus | AGG | S. pyogenes |
| 7 | GCCCAGCUCCAGCCUCUGAUG | AGGGGT | S. aureus | AGG | S. pyogenes |
| 8 | GCCCUGAUCUGCACUGAACAG | AGGGGT | S. aureus | AGG | S. pyogenes |
| 9 | GCCUCAAGUCUGGUUAUUUUAG | GGGGAT | S. aureus | GGG | S. pyogenes |
| 10 | GCCUGGCAGAUGAGAACCAGG | AGGAAT | S. aureus | AGG | S. pyogenes |
| 11 | GCGAAAGGCUCGCGGCGAAGGA | AGGAAT | S. aureus | AGG | S. pyogenes |
| 12 | GCUCCUCUCACCCUUAUGACUC | AGGGAT | S. aureus | AGG | S. pyogenes |
| 13 | GCUGCAAGGGUUGGCCAGGCU | GGGAAT | S. aureus | GGG | S. pyogenes |
| 14 | GGCCUCCGUAUCACUCUCUGAC | TGGGGT | S. aureus | TGG | S. pyogenes |
| 15 | GGGUACCUGAGUGGGGUGCAUU | TGGGGT | S. aureus | TGG | S. pyogenes |
| 16 | GGUCGACCCUUGGUAUCCAUG | GGGGAT | S. aureus | GGG | S. pyogenes |
| 17 | GGUCGUAGCCAGUCCGAACCC | CGGAGT | S. aureus | CGG | S. pyogenes |
| 18 | GUAACUGAACCCCUGCAAUCAA | TGGGAT | S. aureus | TGG | S. pyogenes |
| 19 | GGCCUCCGUAUCACUCUCUGAC | TGGGGT | S. aureus | TGG | S. pyogenes |
| 20 | GCUUUCCUUAGCUGUAAAAGAA | AGGGAT | S. aureus | AGG | S. pyogenes |
| 21 | GUGGCACUGCGGCUGGAGGU | GGGGGT | S. aureus | GGG | S. pyogenes |
| 22 | GUAGGGCCUUCGCGCACCUCA | TGGAAT | S. aureus | TGG | S. pyogenes |
| 23 | GGCCUCCCCAAAGCCUGGCCA | GGGAGT | S. aureus | GGG | S. pyogenes |
| 24 | GCACAUUCACGGUCUCAGUGC | AAGGAT | S. aureus | AAG | S. pyogenes |
| 25 | GGAAACCUUGAAUAAGAAUGGA | AGGGGT | S. aureus | AGG | S. pyogenes |
| 26 | GUAUUACUAUUAUUAUCUGAGA | TGGGGT | S. aureus | TGG | S. pyogenes |
| 27 | GUGGGACUGAUCCCUUAAUGUG | TGGGGT | S. aureus | TGG | S. pyogenes |
| 28 | GAAAGAGACAGAGAAGGGGCA | GGGGGT | S. aureus | GGG | S. pyogenes |
| 29 | GAAGGCUUUACUGUAUUACAGA | AGGGGT | S. aureus | AGG | S. pyogenes |
| 30 | GACCAAAACGAGGGACAUUUA | GGGGAT | S. aureus | GGG | S. pyogenes |
| 31 | GACCAGGUCAGCAAACAUGUU | TGGAAT | S. aureus | TGG | S. pyogenes |
| 32 | GACUCAGCGCCCCUGCCGGGCC | TGGGAT | S. aureus | TGG | S. pyogenes |
| 33 | GAGAAGAAACCAGGGAACAGGU | AGGAGT | S. aureus | AGG | S. pyogenes |
| 34 | GAGUGGGAACUUUCUGAUGCCA | TGGAAT | S. aureus | TGG | S. pyogenes |

**Supplementary Table 4.** DNA sequences of oligos used in this study. Primers for guided off-target[1] and targeted RNA-seq[2] are the same as previous publication and not listed.

Oligos used in vitro assays (adaptor sequences were highlighted in yellow, * stands for phosphorothioate bonds):

oligo 1 (figure 3d):
G*G*TGGTTTGTGTATTGGGTGCCTTCTATTTCCAGCTCGAAGCGAAAAAACAGATAAGTTCATAACC
GCATGTAGGAATTTTGGTGGGA*T*A

oligo 2 (figure 3d):
G*G*TGGTTTGTGTATTGGGTGTATCTTAACAATGTTAATAACGTATAAAGGCTGTTCATTCCCTCGCG
CATGTAGGAATTTTGGTGGGA*T*A

oligo 3 (figure 3e):
T*G*GTTTGTGTATTGGGTGAAGGTGAAAGGGTGAAAAAAATTG**TCT**GTAAGTAAGGGTGGTAAAGAA
TAAATGTAGGAATTTTGGTGGG*A*T

HTS primers:

| Primer name | Primer sequence (5' to 3') |
|---|---|
| HTS-FP-site1 | ACACTCTTTCCCTACACGACGCTCTTCCGATCTACTGTCTTTTGATCTACAGCAGTTAAT |
| HTS-FP-site2 | ACACTCTTTCCCTACACGACGCTCTTCCGATCTAGCCTCTTTCCTGCTAGAGC |
| HTS-FP-site3 | ACACTCTTTCCCTACACGACGCTCTTCCGATCTCTTTCGCTGCCCTTTCCTCT |
| HTS-FP-site4 | ACACTCTTTCCCTACACGACGCTCTTCCGATCTGATATCTCCAGGCTCCTGTCCATTCT |
| HTS-FP-site5 | ACACTCTTTCCCTACACGACGCTCTTCCGATCTCCATCCTAAGTGAAGCAGCATATTTGA |
| HTS-FP-site6 | ACACTCTTTCCCTACACGACGCTCTTCCGATCTAGGTGGGGGTGACTCCTTTTTTGGA |
| HTS-FP-site7 | ACACTCTTTCCCTACACGACGCTCTTCCGATCTCTTGTCTGTCCAAGGAGAATGAGGTC |
| HTS-FP-site8 | ACACTCTTTCCCTACACGACGCTCTTCCGATCTGACCTGGAGGCCTGGGATCCACA |
| HTS-FP-site9 | ACACTCTTTCCCTACACGACGCTCTTCCGATCTCCTTTAGGACACATGCTGTCTACCACA |
| HTS-FP-site10 | ACACTCTTTCCCTACACGACGCTCTTCCGATCTGCCAAAGTCTGAGGTTTAGTTGACTAA |
| HTS-FP-site11 | ACACTCTTTCCCTACACGACGCTCTTCCGATCTGTGGGAACATCACCGGAGCCTGG |
| HTS-FP-site12 | ACACTCTTTCCCTACACGACGCTCTTCCGATCTCTGACACTAAATATGTGGTTTTTTGCT |
| HTS-FP-site13 | ACACTCTTTCCCTACACGACGCTCTTCCGATCTCGAACTCCTAGGCTCAAGTAATCCA |
| HTS-FP-site14 | ACACTCTTTCCCTACACGACGCTCTTCCGATCTGCCAGTAATTGCATTAAACCCTCACTA |
| HTS-FP-site15 | ACACTCTTTCCCTACACGACGCTCTTCCGATCTGGCTCCCACTCTCTCCCAGTGTCCTCA |
| HTS-FP-site16 | ACACTCTTTCCCTACACGACGCTCTTCCGATCTTCTGCCTGTGTGAAGCTCCC |
| HTS-FP-site17 | ACACTCTTTCCCTACACGACGCTCTTCCGATCTGGGAGTCCTCCCTTCACCCCTGC |
| HTS-FP-site18 | ACACTCTTTCCCTACACGACGCTCTTCCGATCTGTGCCAAGGCATAAAAGCCTTCCCTG |
| HTS-FP-site19 | ACACTCTTTCCCTACACGACGCTCTTCCGATCTACTCGCTGGCCTGGCCTTTCTTCTC |
| HTS-FP-site20 | ACACTCTTTCCCTACACGACGCTCTTCCGATCTAAGCGGGTTCTCATTGTTCCCGTGTCT |
| HTS-FP-site21 | ACACTCTTTCCCTACACGACGCTCTTCCGATCTAACCAGTCCCTGTCCTGAATCTATCTA |
| HTS-FP-site22 | ACACTCTTTCCCTACACGACGCTCTTCCGATCTTTGCTTTCGGGTATCTACTAGGAGTCA |
| HTS-FP-site23 | ACACTCTTTCCCTACACGACGCTCTTCCGATCTGGGGCTGGGCTTGCGTTGCCGCT |
| HTS-FP-site24 | ACACTCTTTCCCTACACGACGCTCTTCCGATCTGGGCTATCAAACCTCATGATTGGC |

**Supplementary Table 4 Continued.** DNA sequences of oligos used in this study.

| Primer name | Primer sequence (5' to 3') |
|---|---|
| HTS-FP-site25 | ACACTCTTTCCCTACACGACGCTCTTCCGATCTAAGCTGTCCAGCTGGAAGCCTGGTAA |
| HTS-FP-site26 | ACACTCTTTCCCTACACGACGCTCTTCCGATCTGCCTAAGTTATATGCAAACATCATGCC |
| HTS-FP-site27 | ACACTCTTTCCCTACACGACGCTCTTCCGATCTGCTGCTGGAATACCGAGGAC |
| HTS-FP-site28 | ACACTCTTTCCCTACACGACGCTCTTCCGATCTACGAGGTAAGTGTGTGGATTAGTTTCA |
| HTS-FP-site29 | ACACTCTTTCCCTACACGACGCTCTTCCGATCTAGTGGTTACTTTGCCGGGTT |
| HTS-FP-site30 | ACACTCTTTCCCTACACGACGCTCTTCCGATCTNNNNGAACCCAGGTAGCCAGAGAC |
| HTS-FP-site31 | ACACTCTTTCCCTACACGACGCTCTTCCGATCTNNNNCATTGCAGAGAGGCGTATCA |
| HTS-FP-site32 | ACACTCTTTCCCTACACGACGCTCTTCCGATCTNNNNCAGAGTGCTGCTTGCTGCT |
| HTS-FP-site33 | ACACTCTTTCCCTACACGACGCTCTTCCGATCTTTTAGTGACTAGCCGCCACC |
| HTS-FP-site34 | ACACTCTTTCCCTACACGACGCTCTTCCGATCTNNNNGAAACCATGTCTCTGGATGCC |
| HTS-FP-site35 | ACACTCTTTCCCTACACGACGCTCTTCCGATCTNNNNAGGCCTTTTCTTGGGGATGC |
| HTS-RP-site1 | TGGAGTTCAGACGTGTGCTCTTCCGATCTAAGAAACAGATTACAGAAGTAGATGCA |
| HTS-RP-site2 | TGGAGTTCAGACGTGTGCTCTTCCGATCTTCTCTCCTATGTGCTGGCCT |
| HTS-RP-site3 | TGGAGTTCAGACGTGTGCTCTTCCGATCTCTACACTGGAACCCCGACTC |
| HTS-RP-site4 | TGGAGTTCAGACGTGTGCTCTTCCGATCTCCAGCCGATATTTCAGAACTAATCAGA |
| HTS-RP-site5 | TGGAGTTCAGACGTGTGCTCTTCCGATCTAACAATGGCAAGGGCCTGCCCTG |
| HTS-RP-site6 | TGGAGTTCAGACGTGTGCTCTTCCGATCTGGGCAGAAGGAAAAATCTATCCTGGAA |
| HTS-RP-site7 | TGGAGTTCAGACGTGTGCTCTTCCGATCTGCACAGAACCCGCTGCTAGAGACTCCA |
| HTS-RP-site8 | TGGAGTTCAGACGTGTGCTCTTCCGATCTGGAAAGTCTGGTTAGAGCTCAGAGGGA |
| HTS-RP-site9 | TGGAGTTCAGACGTGTGCTCTTCCGATCTGTGGTGGAGTGCTCTGTGTTTGTCT |
| HTS-RP-site10 | TGGAGTTCAGACGTGTGCTCTTCCGATCTATTACAGGTGTGGGCCACCTTGCCC |
| HTS-RP-site11 | TGGAGTTCAGACGTGTGCTCTTCCGATCTTGCATAACCTACACACATCCTCTGATA |
| HTS-RP-site12 | TGGAGTTCAGACGTGTGCTCTTCCGATCTGGATTGCGGAAATCCCCAACTTATAGC |
| HTS-RP-site13 | TGGAGTTCAGACGTGTGCTCTTCCGATCTGCCTGGACTCCAGACAGGCTTCC |
| HTS-RP-site14 | TGGAGTTCAGACGTGTGCTCTTCCGATCTAAGGCCAAGAATCTTGCTAGTAGTGGA |
| HTS-RP-site15 | TGGAGTTCAGACGTGTGCTCTTCCGATCTGGATAGAGCAAAAGAAGTAGTGCCTGG |
| HTS-RP-site16 | TGGAGTTCAGACGTGTGCTCTTCCGATCTTGAAACTGTCACTGAAACATCTGGT |
| HTS-RP-site17 | TGGAGTTCAGACGTGTGCTCTTCCGATCTGTTCTCAAGAAAAGGCCACCCCTCAG |
| HTS-RP-site18 | TGGAGTTCAGACGTGTGCTCTTCCGATCTTGCTTAGAGGGTAAAAACCCAGGAGGA |
| HTS-RP-site19 | TGGAGTTCAGACGTGTGCTCTTCCGATCTGGGAGAGAGGCAGGGCGGGCATG |
| HTS-RP-site20 | TGGAGTTCAGACGTGTGCTCTTCCGATCTTCCGCCTCCGGAGTAGGGCTGCAGAGA |
| HTS-RP-site21 | TGGAGTTCAGACGTGTGCTCTTCCGATCTGGAAGGCAGACTGTATCTGGTCTTTT |
| HTS-RP-site22 | TGGAGTTCAGACGTGTGCTCTTCCGATCTTCTAGCAGGAAAGAGGCTCAGGCCCA |
| HTS-RP-site23 | TGGAGTTCAGACGTGTGCTCTTCCGATCTAGACCGAGTGGCAGTGACAGCAAGC |
| HTS-RP-site24 | TGGAGTTCAGACGTGTGCTCTTCCGATCTACACACAGACACTGCAGAGAATAACA |
| HTS-RP-site25 | TGGAGTTCAGACGTGTGCTCTTCCGATCTCCGCCCAGCACTCGCAGAGCAGA |
| HTS-RP-site26 | TGGAGTTCAGACGTGTGCTCTTCCGATCTGATGAGAATGCACCATGATTCCAATCA |
| HTS-RP-site27 | TGGAGTTCAGACGTGTGCTCTTCCGATCTGCAACTCTCTTTTCTCCGGGA |
| HTS-RP-site28 | TGGAGTTCAGACGTGTGCTCTTCCGATCTCTACCAAGGAGAGTCATTCCTTTCAGA |
| HTS-RP-site29 | TGGAGTTCAGACGTGTGCTCTTCCGATCTAAGACAGTCTGGGAAGCGTG |
| HTS-RP-site30 | TGGAGTTCAGACGTGTGCTCTTCCGATCTTCCTTTCAACCCGAACGGAG |
| HTS-RP-site31 | TGGAGTTCAGACGTGTGCTCTTCCGATCTGGGGTCCCAGGTGCTGAC |
| HTS-RP-site32 | TGGAGTTCAGACGTGTGCTCTTCCGATCTAAAAGGGAGATTGGAGACACGGAGA |
| HTS-RP-site33 | TGGAGTTCAGACGTGTGCTCTTCCGATCTTGCGCTTTACAGGTCTCCAG |
| HTS-RP-site34 | TGGAGTTCAGACGTGTGCTCTTCCGATCTAGAGAAATCACACTAGCTAGCCT |
| HTS-RP-site35 | TGGAGTTCAGACGTGTGCTCTTCCGATCTAGAGAAATCACACTAGCTAGCCT |
| HTS-FP-ssoligo | ACACTCTTTCCCTACACGACGCTCTTCCGATCTNNNNGGTGGTTTGTGTATTGGGTG |
| HTS-RP-ssoligo | TGGAGTTCAGACGTGTGCTCTTCCGATCTTATCCCACCAAAATTCCTACAT |

**Supplementary Table 4.** DNA sequences of mammalian expression plasmids for the core CBEs showed in this study. The deaminase sequence is highlighted for BE4-rAPOBEC1. For the rest of constructs, only the deaminase sequences are shown. The backbone sequences are identical.

**BE4-rAPOBEC1**

TGCTTCGCGATGTACGGGCCAGATATACGCGTTGACATTGATTATTGACTAGTTATTAATAGTAATCAATTACGGGGTCATTAGT
TCATAGCCCATATATGGAGTTCCGCGTTACATAACTTACGGTAAATGGCCCGCCTGGCTGACCGCCCAACGACCCCCGCCCAT
TGACGTCAATAATGACGTATGTTCCCATAGTAACGCCAATAGGGACTTTCCATTGACGTCAATGGGTGGAGTATTTACGGTAAAC
TGCCCACTTGGCAGTACATCAAGTGTATCATATGCCAAGTACGCCCCCTATTGACGTCAATGACGGTAAATGGCCCGCCTGGCA
TTATGCCCAGTACATGACCTTATGGGACTTTCCTACTTGGCAGTACATCTACGTATTAGTCATCGCTATTACCATGGTGATGCGG
TTTTGGCAGTACATCAATGGGCGTGGATAGCGGTTTGACTCACGGGGATTTCCAAGTCTCCACCCCATTGACGTCAATGGGAGT
TTGTTTTGGCACCAAAATCAACGGGACTTTCCAAAATGTCGTAACAACTCCGCCCCATTGACGCAAATGGGCGGTAGGCGTGTA
CGGTGGGAGGTCTATATAAGCAGAGCTGGTTTAGTGAACCGTCAGATCCGCTAGAGATCCGCGGCCGCTAATACGACTCACTA
TAGGGAGAGCCGCCACCATGAGCAGCGAGACAGGCCCTGTGGCCGTGGACCCCACCCTGCGGCGGAGAATCGAGCCTCATG
AGTTCGAGGTGTTCTTCGACCCTCGGGAACTGAGAAAAGAGACATGCCTGCTGTACGAGATCAACTGGGGCGGAAGACACAGC
ATCTGGCGGCACACCAGCCAGAACACCAACAAGCACGTGGAAGTGAATTTCATCGAGAAGTTCACCACCGAAAGATACTTCTG
CCCCAACACCAGATGCAGCATCACATGGTTCCTGTCTTGGTCCCCTTGCGGCGAGTGCTCTAGAGCCATCACCGAGTTCCTGA
GCAGATATCCTCACGTGACACTGTTCATCTACATCGCCAGACTGTATCACCACGCCGATCCTAGAAATAGACAGGGCCTGCGG
GACCTGATCAGCTCCGGCGTGACCATCCAGATCATGACCGAGCAGGAGAGCGGCTACTGTTGGAGAAACTTCGTGAACTACTC
TCCTAGCAACGAGGCCCACTGGCCTAGATACCCCCACCTGTGGGTGCGGCTGTACGTGCTGGAACTGTACTGCATCATCCTGG
GACTGCCTCCATGTCTGAACATCCTGAGAAGAAAGCAGCCTCAGCTGACCTTCTTCACAATCGCCCTGCAGAGCTGCCACTAC
CAGAGACTGCCCCCCCACATCCTGTGGGCCACCGGCCTGAAGCTTAAGAGCGGAGGATCTCTTAAGAGCGGAGGATCTAGCG
GCGGCTCTAGCGGATCTGAGACACCTGGCACAAGCGAGTCTGCCACACCTGAGAGTAGCGGCGGATCTTCTGGTGGCTCTGA
CAAGAAGTACAGCATCGGCCTGGCCATCGGCACCAACTCTGTGGGCTGGGCCGTGATCACCGACGAGTACAAGGTGCCCAGC
AAGAAATTCAAGGTGCTGGGCAACACCGACCGGCACAGCATCAAGAAGAACCTGATCGGAGCCCTGCTGTTCGACAGCGGCG
AAACAGCCGAGGCCACCCGGCTGAAGAGAACCGCCAGAAGAAGATACACCAGACGGAAGAACCGGATCTGCTATCTGCAAGA
GATCTTCAGCAACGAGATGGCCAAGGTGGACGACAGCTTCTTCCACAGACTGGAAGAGTCCTTCCTGGTGGAAGAGGATAAGA
AGCACGAGCGGCACCCCATCTTCGGCAACATCGTGGACGAGGTGGCCTACCACGAGAAGTACCCCACCATCTACCACCTGAG
AAAGAAACTGGTGGACAGCACCGACAAGGCCGACCTGCGGCTGATCTATCTGGCCCTGGCCCACATGATCAAGTTCCGGGGC
CACTTCCTGATCGAGGGCGACCTGAACCCCGACAACAGCGACGTGGACAAGCTGTTCATCCAGCTGGTGCAGACCTACAACCA
GCTGTTCGAGGAAAACCCCATCAACGCCAGCGGCGTGGACGCCAAGGCCATCCTGTCTGCCAGACTGAGCAAGAGCAGACGG
CTGGAAAATCTGATCGCCCAGCTGCCCGGCGAGAAGAAGAATGGCCTGTTCGGAAACCTGATTGCCCTGAGCCTGGGCCTGA
CCCCCAACTTCAAGAGCAACTTCGACCTGGCCGAGGATGCCAAACTGCAGCTGAGCAAGGACACCTACGACGACGACCTGGA
CAACCTGCTGGCCCAGATCGGCGACCAGTACGCCGACCTGTTTCTGGCCGCCAAGAACCTGTCCGACGCCATCCTGCTGAGC
GACATCCTGAGAGTGAACACCGAGATCACCAAGGCCCCCCTGAGCGCCTCTATGATCAAGAGATACGACGAGCACCACCAGG
ACCTGACCCTGCTGAAAGCTCTCGTGCGGCAGCAGCTGCCTGAGAAGTACAAAGAGATTTTCTTCGACCAGAGCAAGAACGGC
TACGCCGGCTACATTGACGGCGGAGCCAGCCAGGAAGAGTTCTACAAGTTCATCAAGCCCATCCTGGAAAAGATGGACGGCAC
CGAGGAACTGCTCGTGAAGCTGAACAGAGAGGACCTGCTGCGGAAGCAGCGGACCTTCGACAACGGCAGCATCCCCCACCAG
ATCCACCTGGGAGAGCTGCACGCCATTCTGCGGCGGCAGGAAGATTTTTACCCATTCCTGAAGGACAACCGGGAAAAGATCGA
GAAGATCCTGACCTTCCGCATCCCCTACTACGTGGGCCCTCTGGCCAGGGGAAACAGCAGATTCGCCTGGATGACCAGAAAGA
GCGAGGAAACCATCACCCCCTGGAACTTCGAGGAAGTGGTGGACAAGGGCGCTTCCGCCCAGAGCTTCATCGAGCGGATGAC
CAACTTCGATAAGAACCTGCCCAACGAGAAGGTGCTGCCCAAGCACAGCCTGCTGTACGAGTACTTCACCGTGTATAACGAGC
TGACCAAAGTGAAATACGTGACCGAGGGAATGAGAAAGCCCGCCTTCCTGAGCGGCGAGCAGAAAAAGGCCATCGTGGACCT
GCTGTTCAAGACCAACCGGAAAGTGACCGTGAAGCAGCTGAAAGAGGACTACTTCAAGAAAATCGAGTGCTTCGACTCCGTGG
AAATCTCCGGCGTGGAAGATCGGTTCAACGCCTCCCTGGGCACATACCACGATCTGCTGAAAATTATCAAGGACAAGGACTTC
CTGGACAATGAGGAAAACGAGGACATTCTGGAAGATATCGTGCTGACCCTGACACTGTTTGAGGACAGAGAGATGATCGAGGA
ACGGCTGAAAACCTATGCCCACCTGTTCGACGACAAAGTGATGAAGCAGCTGAAGCGGCGGAGATACACCGGCTGGGGCAGG
CTGAGCCGGAAGCTGATCAACGGCATCCGGGACAAGCAGTCCGGCAAGACAATCCTGGATTTCCTGAAGTCCGACGGCTTCG
CCAACAGAAACTTCATGCAGCTGATCCACGACGACAGCCTGACCTTTAAAGAGGACATCCAGAAAGCCCAGGTGTCCGGCCAG
GGCGATAGCCTGCACGAGCACATTGCCAATCTGGCCGGCAGCCCCGCCATTAAGAAGGGCATCCTGCAGACAGTGAAGGTGG
TGGACGAGCTCGTGAAAGTGATGGGCCGGCACAAGCCCGAGAACATCGTGATCGAAATGGCCAGAGAGAACCAGACCACCCA
GAAGGGACAGAAGAACAGCCGCGAGAGAATGAAGCGGATCGAAGAGGGCATCAAAGAGCTGGGCAGCCAGATCCTGAAAGAA
CACCCCGTGGAAAACACCCAGCTGCAGAACGAGAAGCTGTACCTGTACTACCTGCAGAATGGGCGGGATATGTACGTGGACCA
GGAACTGGACATCAACCGGCTGTCCGACTACGATGTGGACCATATCGTGCCTCAGAGCTTTCTGAAGGACGACTCCATCGACA
ACAAGGTGCTGACCAGAAGCGACAAGAACCGGGGCAAGAGCGACAACGTGCCCTCCGAAGAGGTCGTGAAGAAGATGAAGAA
CTACTGGCGGCAGCTGCTGAACGCCAAGCTGATTACCCAGAGAAAGTTCGACAATCTGACCAAGGCCGAGAGAGGCGGCCTG
AGCGAACTGGATAAGGCCGGCTTCATCAAGAGACAGCTGGTGGAAACCCGGCAGATCACAAAGCACGTGGCACAGATCCTGG
ACTCCCGGATGAACACTAAGTACGACGAGAATGACAAGCTGATCCGGGAAGTGAAAGTGATCACCCTGAAGTCCAAGCTGGTG
TCCGATTTCCGGAAGGATTTCCAGTTTTACAAAGTGCGCGAGATCAACAACTACCACCACGCCCACGACGCCTACCTGAACGC
CGTCGTGGGAACCGCCCTGATCAAAAAGTACCCTAAGCTGGAAAGCGAGTTCGTGTACGGCGACTACAAGGTGTACGACGTGC
GGAAGATGATCGCCAAGAGCGAGCAGGAAATCGGCAAGGCTACCGCCAAGTACTTCTTCTACAGCAACATCATGAACTTTTTCA
AGACCGAGATTACCCTGGCCAACGGCGAGATCCGGAAGCGGCCTCTGATCGAGACAAACGGCGAAACCGGGGAGATCGTGTG

GGATAAGGGCCGGGATTTTGCCACCGTGCGGAAAGTGCTGAGCATGCCCCAAGTGAATATCGTGAAAAAGACCGAGGTGCAG
ACAGGCGGCTTCAGCAAAGAGTCTATCCTGCCCAAGAGGAACAGCGATAAGCTGATCGCCAGAAAGAAGGACTGGGACCCTAA
GAAGTACGGCGGCTTCGACAGCCCCACCGTGGCCTATTCTGTGCTGGTGGTGGCCAAAGTGGAAAAGGGCAAGTCCAAGAAA
CTGAAGAGTGTGAAAGAGCTGCTGGGGATCACCATCATGGAAAGAAGCAGCTTCGAGAAGAATCCCATCGACTTTCTGGAAGC
CAAGGGCTACAAAGAAGTGAAAAAGGACCTGATCATCAAGCTGCCTAAGTACTCCCTGTTCGAGCTGGAAAACGGCCGGAAGA
GAATGCTGGCCTCTGCCGGCGAACTGCAGAAGGGAAACGAACTGGCCCTGCCCTCCAAATATGTGAACTTCCTGTACCTGGCC
AGCCACTATGAGAAGCTGAAGGGCTCCCCCGAGGATAATGAGCAGAAACAGCTGTTTGTGGAACAGCACAAGCACTACCTGGA
CGAGATCATCGAGCAGATCAGCGAGTTCTCCAAGAGAGTGATCCTGGCCGACGCTAATCTGGACAAAGTGCTGTCCGCCTACA
ACAAGCACCGGGATAAGCCCATCAGAGAGCAGGCCGAGAATATCATCCACCTGTTTACCCTGACCAATCTGGGAGCCCCTGCC
GCCTTCAAGTACTTTGACACCACCATCGACCGGAAGAGGTACACCAGCACCAAAGAGGTGCTGGACGCCACCCTGATCCACCA
GAGCATCACCGGCCTGTACGAGACACGGATCGACCTGTCTCAGCTGGGAGGTGACTCTGGTGGAAGCGGAGGATCTGGCGG
CAGCACCAATCTGAGCGACATCATCGAGAAAGAGACAGGCAAGCAGCTGGTCATCCAAGAGTCCATCCTGATGCTGCCTGAAG
AGGTGGAAGAAGTGATCGGCAACAAGCCCGAGTCCGACATCCTGGTGCACACCGCCTACGATGAGAGCACCGACGAGAACGT
GATGCTGCTGACCTCTGACGCCCCTGAGTACAAGCCTTGGGCTCTCGTGATCCAGGACAGCAACGGCGAGAACAAGATCAAGA
TGCTGAGCGGCGGCTCTGGTGGCTCTGGCGGATCTACAAACCTGTCCGATATTATTGAGAAAGAAACCGGGAAACAGCTCGTG
ATTCAAGAGTCTATTCTCATGCTCCCGGAAGAAGTCGAGGAAGTCATTGGAAACAAGCCTGAGAGCGATATTCTGGTCCATACA
GCCTACGACGAGTCTACCGATGAGAATGTCATGCTCCTCACCAGCGACGCTCCCGAGTATAAGCCATGGGCACTTGTCATTCA
GGACTCCAATGGGGAAAACAAAATCAAAATGCTCCCAAAGAAAAAACGCAAGGTGGAGGGAGCTGATAAGCGCACCGCCGATG
GTTCCGAGTTCGAAAGCCCCAAGAAGAAGAGGAAAGTCTAACCGGTCATCATCACCATCACCATTGAGTTTAAACCCGCTGATC
AGCCTCGACTGTGCCTTCTAGTTGCCAGCCATCTGTTGTTTGCCCCTCCCCCGTGCCTTCCTTGACCCTGGAAGGTGCCACTCC
CACTGTCCTTTCCTAATAAAATGAGGAAATTGCATCGCATTGTCTGAGTAGGTGTCATTCTATTCTGGGGGGTGGGGTGGGGCA
GGACAGCAAGGGGGAGGATTGGGAAGACAATAGCAGGCATGCTGGGGATGCGGTGGGCTCTATGGCTTCTGAGGCGGAAAG
AACCAGCTGGGGCTCGATACCGTCGACCTCTAGCTAGAGCTTGGCGTAATCATGGTCATAGCTGTTTCCTGTGTGAAATTGTTA
TCCGCTCACAATTCCACACAACATACGAGCCGGAAGCATAAAGTGTAAAGCCTAGGGTGCCTAATGAGTGAGCTAACTCACATT
AATTGCGTTGCGCTCACTGCCCGCTTTCCAGTCGGGAAACCTGTCGTGCCAGCTGCATTAATGAATCGGCCAACGCGCGGGGA
GAGGCGGTTTGCGTATTGGGCGCTCTTCCGCTTCCTCGCTCACTGACTCGCTGCGCTCGGTCGTTCGGCTGCGGCGAGCGGT
ATCAGCTCACTCAAAGGCGGTAATACGGTTATCCACAGAATCAGGGGATAACGCAGGAAAGAACATGTGAGCAAAAGGCCAGC
AAAAGGCCAGGAACCGTAAAAAGGCCGCGTTGCTGGCGTTTTTCCATAGGCTCCGCCCCCCTGACGAGCATCACAAAAATCGA
CGCTCAAGTCAGAGGTGGCGAAACCCGACAGGACTATAAAGATACCAGGCGTTTCCCCCTGGAAGCTCCCTCGTGCGCTCTCC
TGTTCCGACCCTGCCGCTTACCGGATACCTGTCCGCCTTTCTCCCTTCGGGAAGCGTGGCGCTTTCTCATAGCTCACGCTGTA
GGTATCTCAGTTCGGTGTAGGTCGTTCGCTCCAAGCTGGGCTGTGTGCACGAACCCCCCGTTCAGCCCGACCGCTGCGCCTTA
TCCGGTAACTATCGTCTTGAGTCCAACCCGGTAAGACACGACTTATCGCCACTGGCAGCAGCCACTGGTAACAGGATTAGCAG
AGCGAGGTATGTAGGCGGTGCTACAGAGTTCTTGAAGTGGTGGCCTAACTACGGCTACACTAGAAGAACAGTATTTGGTATCTG
CGCTCTGCTGAAGCCAGTTACCTTCGGAAAAAGAGTTGGTAGCTCTTGATCCGGCAAACAAACCACCGCTGGTAGCGGTGGTT
TTTTTGTTTGCAAGCAGCAGATTACGCGCAGAAAAAAAGGATCTCAAGAAGATCCTTTGATCTTTTCTACGGGGTCTGACGCTCA
GTGGAACGAAAACTCACGTTAAGGGATTTTGGTCATGAGATTATCAAAAAGGATCTTCACCTAGATCCTTTTAAATTAAAAATGAA
GTTTTAAATCAATCTAAAGTATATATGAGTAAACTTGGTCTGACAGTTACCAATGCTTAATCAGTGAGGCACCTATCTCAGCGATC
TGTCTATTTCGTTCATCCATAGTTGCCTGACTCCCCGTCGTGTAGATAACTACGATACGGGAGGGCTTACCATCTGGCCCCAGT
GCTGCAATGATACCGCGAGACCCACGCTCACCGGCTCCAGATTTATCAGCAATAAACCAGCCAGCCGGAAGGGCCGAGCGCA
GAAGTGGTCCTGCAACTTTATCCGCCTCCATCCAGTCTATTAATTGTTGCCGGGAAGCTAGAGTAAGTAGTTCGCCAGTTAATA
GTTTGCGCAACGTTGTTGCCATTGCTACAGGCATCGTGGTGTCACGCTCGTCGTTTGGTATGGCTTCATTCAGCTCCGGTTCCC
AACGATCAAGGCGAGTTACATGATCCCCCATGTTGTGCAAAAAAGCGGTTAGCTCCTTCGGTCCTCCGATCGTTGTCAGAAGTA
AGTTGGCCGCAGTGTTATCACTCATGGTTATGGCAGCACTGCATAATTCTCTTACTGTCATGCCATCCGTAAGATGCTTTTCTGT
GACTGGTGAGTACTCAACCAAGTCATTCTGAGAATAGTGTATGCGGCGACCGAGTTGCTCTTGCCCGGCGTCAATACGGGATA
ATACCGCGCCACATAGCAGAACTTTAAAAGTGCTCATCATTGGAAAACGTTCTTCGGGGCGAAAACTCTCAAGGATCTTACCGC
TGTTGAGATCCAGTTCGATGTAACCCACTCGTGCACCCAACTGATCTTCAGCATCTTTTACTTTCACCAGCGTTTCTGGGTGAGC
AAAAACAGGAAGGCAAAATGCCGCAAAAAAGGGAATAAGGGCGACACGGAAATGTTGAATACTCATACTCTTCCTTTTTCAATAT
TATTGAAGCATTTATCAGGGTTATTGTCTCATGAGCGGATACATATTTGAATGTATTTAGAAAAATAAACAAATAGGGGTTCCGCG
CACATTTCCCCGAAAAGTGCCACCTGACGTCGACGGATCGGGAGATCGATCTCCCGATCCCCTAGGGTCTTACTCTCAGTACA
ATCTGCTCTGATGCCGCATAGTTAAGCCAGTATCTGCTCCCTGCTTGTGTGTTGGAGGTCGCTGAGTAGTGCGCGAGCAAAATT
TAAGCTACAACAAGGCAAGGCTTGACCGACAATTGCATGAAGAATCTGCTTAGGGTTAGGCGTTTTGCGC

**BE4-PpAPOBEC1**

ATGACCTCTGAGAAGGGCCCTAGCACAGGCGACCCCACCCTGCGGCGGAGAATCGAGAGCTGGGAGTTCGACGTGTTCTACG
ACCCTAGAGAACTGAGAAAGGAAACCTGCCTGCTGTACGAGATCAAGTGGGGCATGAGCAGAAAGATCTGGCGGAGCTCTGG
CAAGAACACCACCAACCACGTGGAAGTGAATTTCATCAAGAAGTTCACCAGCGAGAGAAGGTTCCACAGCAGCATCAGCTGCA
GCATCACCTGGTTCCTGAGCTGGTCCCCTTGCTGGGAATGCAGCCAGGCCATCAGAGAGTTCCTGAGCCAACACCCCGGAGT
GACACTGGTGATCTACGTGGCCAGACTGTTCTGGCACATGGACCAGAGAAACAGACAGGGCCTGAGAGATCTGGTCAACAGC
GGCGTGACTATCCAGATCATGCGGGCCAGCGAGTACTACCACTGTTGGCGGAACTTCGTGAACTACCCCCCCGGCGATGAGG
CCCACTGGCCTCAGTACCCTCCTCTGTGGATGATGCTGTACGCCCTGGAACTGCACTGCATCATCCTGTCTCTGCCTCCATGTC
TGAAGATCTCTAGAAGATGGCAGAACCACCTGGCCTTCTTCAGACTGCACCTGCAGAATTGCCACTACCAGACCATCCCCCCC
CACATCCTGCTGGCTACAGGCCTGATCCACCCTTCTGTGACCTGGAGA

**BE4-RrA3F**

ATGAAGCCCCAGATCAGGGACCACCGCCCCAATCCTATGGAGGCCATGTACCCTCACATCTTCTATTTTCACTTCGAGAACCTG
GAGAAGGCCTACGGCCGGAATGAGACCTGGCTGTGCTTTACAGTGGAGATCATCAAGCAGTATCTGCCAGTGCCCTGGAAGAA
GGGCGTGTTCCGGAACCAGGTGGATCCAGAGACCCACTGCCACGCCGAGAAGTGTTTTCTGTCCTGGTTCTGTAACAATACAC
TGTCTCCCAAGAAGAATTACCAGGTGACCTGGTATACAAGCTGGTCCCCTTGCCCAGAGTGTGCAGGAGAGGTGGCAGAGTTT
CTGGCAGAGCACAGCAACGTGAAGCTGACCATCTACACAGCCCGGCTGTACTATTTCTGGGACACCGATTATCAGGAGGGCCT
GAGATCTCTGAGCGAGGAGGGCGCCTCCGTGGAGATCATGGACTACGAGGATTTTCAGTATTGCTGGGAGAACTTCGTGTACG
ACGATGGCGAGCCTTTTAAGAGGTGGAAGGGCCTGAAGTATAATTTCCAGTCTCTGACACGGAGACTGCGCGAGATCCTGCAG

**BE4-AmAPOBEC1**

ATGGCCGACAGCTCCGAGAAGATGAGGGGCCAGTACATCAGCCGCGACACCTTTGAGAAGAATTATAAGCCCATCGATGGCAC
AAAGGAGGCCCACCTGCTGTGCGAGATCAAGTGGGGCAAGTACGGCAAGCCTTGGCTGCACTGGTGTCAGAATCAGCGGATG
AACATCCACGCCGAGGACTATTTCATGAACAATATCTTTAAGGCCAAGAAGCACCCTGTGCACTGCTACGTGACCTGGTATCTG
TCTTGGAGCCCATGCGCCGATTGTGCCTCCAAGATCGTGAAGTTCCTGGAGGAGCGGCCCTACCTGAAGCTGACCATCTATGT
GGCCCAGCTGTACTATCACACAGAGGAGGAGAATAGGAAGGGCCTGCGGCTGCTGCGGAGCAAGAAAGTGATCATCCGCGTG
ATGGACATCTCCGATTACAACTATTGCTGGAAGGTGTTCGTGTCTAACCAGAATGGCAACGAGGACTACTGGCCACTGCAGTTT
GATCCCTGGGTGAAGGAGAATTATTCTCGGCTGCTGGATATCTTCTGGGAGTCCAAGTGTAGATCTCCCAACCCTTGG

**BE4-SsAPOBEC3B**

ATGGACCCACAGAGGCTGCGCCAGTGGCCCGGCCCTGGCCCAGCAAGCAGGGGCGGCTACGGCCAGCGGCCAAGAATCAG
GAACCCCGAGGAGTGGTTTCACGAGCTGTCTCCCCGGACCTTCAGCTTTCACTTCCGCAACCTGAGGTTCGCATCCGGCCGCA
ATCGGTCTTATATCTGCTGTCAGGTGGAGGGCAAGAACTGCTTCTTTCAGGGCATCTTTCAGAATCAGGTGCCACCTGACCCAC
CATGCCACGCAGAGCTGTGCTTCCTGTCTTGGTTCCAGAGCTGGGGCCTGTCCCCCGATGAGCACTACTATGTGACATGGTTT
ATCTCTTGGAGCCCTTGCTGTGAGTGTGCCGCCAAGGTGGCCCAGTTCCTGGAGGAGAACCGCAACGTGAGCCTGTCTCTGA
GCGCCGCAAGGCTGTACTATTTCTGGAAGTCCGAGTCTAGAGAGGGACTGCGGAGACTGAGCGACCTGGGAGCACAAGTGGG
AATCATGTCCTTTCAGGATTTCCAGCACTGCTGGAACAATTTTGTGCACAACCTGGGCATGCCCTTCCAGCCTTGGAAGAAGCT
GCACAAGAATTACCAGAGGCTGGTGACCGAGCTGAAGCAGATCCTGCGCGAGGAGCCTGCCACATATGGCTCTCCACAGGCC
CAGGGCAAGGTGAGAATCGGAAGCACCGCAGCAGGACTGAGGCACAGCCACTCCCACACACGCTCCGAGGCACACCTGAGG
CCTAACCACAGCTCCAGACAGCACAGGATCCTGAATCCTCCACGGGAGGCCAGAGCCAGGACCTGCGTGCTGGTGGATGCCT
CTTGGATCTGTTACAGA

**Supplementary Note 1:** Methods of creating similarity network of cytidine deaminases. To focus the search space within the APOBEC1-like protein family, human APOBEC1 was used as a query sequence for a protein BLAST search against the NCBI non-redundant protein sequences database (nr_v5). The top 1000 sequences were used to generate a sequence similarity network (SSN) with a protein BLAST -log(E-value) edge-threshold of 115. A set of 43 deaminases was selected to sample the sequence space within the SSN.

To identify deaminases from other families that could act as base-editing enzymes, we sampled 80 sequences from a SSN built from all deaminases with the following InterPro annotations IPR002125 (Cytidine and deoxycytidylate deaminase domain), IPR016192 (APOBEC/CMP deaminase, zinc-binding), and IPR016193 (Cytidine deaminase-like). This set of 82,043 sequences was first clustered at 55% identity using Cd-HIT[3] before generating a SSN network by protein BLAST with a -log(E-value) edge-threshold of 50. Sequences were chosen based on their centrality within a cluster of sequence in the network.

**Supplementary Note 2:** Discussion about protein expression level of base editors.

We would like to examine if different protein expression level of editors contributes to changes in *cis/trans* editing profile. Quantification of base editor mRNA and protein was performed on cells transfected with editor plasmids (**Supplementary Figure 12a-b)**, we showed that HiFi mutations like K34A and H122A didn't cause significant changes in base editor transcription and translation. For all 4 new CBEs we characterized in this study, the protein expression level was not dramatically lower than BE4-rAPOBEC1 (**Supplementary Figure 12c)**. As a result, we believe that changes in *cis/trans* editing profile came from the intrinsic characteristics of deaminases.

**Supplementary Figure 14.** Quantification of CBE protein concentration in HEK293T cells transfected with base editor expression plasmids. Base editor protein concentration quantified by measuring the total Cas9 protein concentration and the amount of total protein in cell lysate. BE protein concentration was normalized to BE4-rAPOBEC1. Values and error bars reflect the mean and s.d. of two or more independent biological replicates.

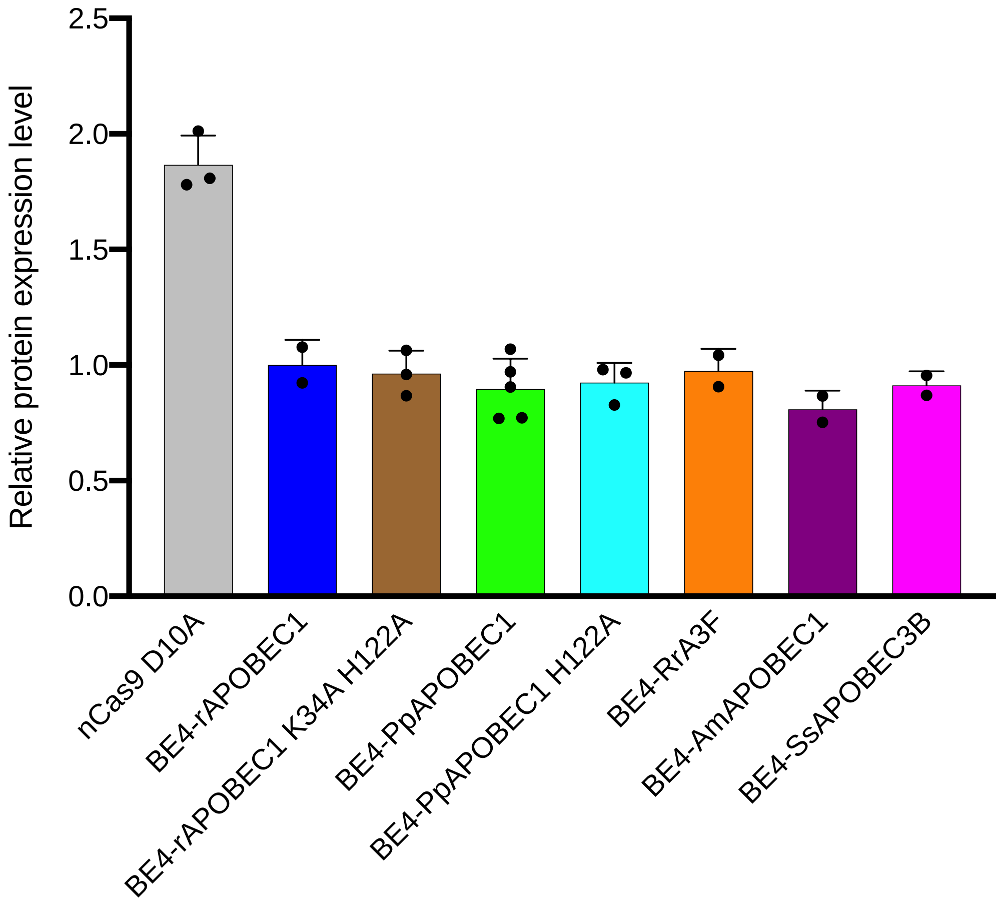

**Supplementary Note 3:** Targeted NGS analysis details

1. To generate FASTQ files from the base call files (BCF) generated by the MiSeq, demultiplexing was performed by running Illumina bcl2fastq (v2.20.0.422) with the following parameters:

```
bcl2fastq \
    --ignore-missing-bcls \
    --ignore-missing-filter \
    --ignore-missing-positions \
    --ignore-missing-controls \
    --auto-set-to-zero-barcode-mismatches \
    --find-adapters-with-sliding-window \
    --adapter-stringency 0.9 \
    --mask-short-adapter-reads 35 \
    --minimum-trimmed-read-length 35 \
```

2. The FASTQ files created in step (1) were processed using trimmomatic (v0.39)[4] with parameters set up to clip Illumina TruSeq adapters, exclude reads shorter than 20 bases, and trim the remaining 3' end of reads if the average base quality (Phred score) in a 4-bp sliding window dropped below 15. In addition, any bases with quality scores of 3 or lower at the end of reads were removed. Finally, because the round 1 PCR primers include four randomized bases after the read 1 primer sequence, the first four bases of each read were trimmed. The command used to execute trimmomatic is shown below:

```
trimmomatic SE -phred33 $input_fastq $output_fastq \
    ILLUMINACLIP:illumine_adapters.fa:2:30:10 \
    LEADING:3 TRAILING:3 \
    SLIDINGWINDOW:4:15 \
    MINLEN:20 \
    HEADCROP:4
```

3. Reads were aligned to amplicon sequences using bowtie2 (v2.35)[5], in end-to-end mode with the alignment parameters specified by the --very sensitive flag. Reference sequences were determined as the expected amplicon sequences (including primers) for each primer pair based on the human genome (GRCh38).The SAM files created by bowtie2 were converted to BAM files, sorted, and indexed using the samtools package (v1.9)[6]. Only samples with at least 5,000 aligned reads were considered for analysis.

4. The BAM files created in step (3) were processed using the bam-readcounts tool (https://github.com/genome/bam-readcount) to generate plain text files summarizing the number of non-reference bases, deletions and insertions at each position in the alignment. The minimum base quality (Phred score) for counting a non-reference base was set to 29 in order to exclude low confidence base calls from statistics about editing rates. Only reads with insertions and/or deletions that overlapped the base editor target site (defined as its protospacer + PAM sequence) were counted towards insertion and deletion rates. Editing rates for each position in the target site were calculated as the fraction of non-reference bases of a given type (e.g., G) to the total number of bases passing the base quality threshold at a given position in the alignment.

**Supplementary Note 4:** Transcriptome sequencing analysis method.

FASTQ files were downloaded from Novagene and aligned to the human genome (Gencode GRCh38v31) using STAR (v2.7.2a). Genome alignments were then duplicate marked and sorted with Picard (v2.20.5). Reads that contain Ns in their cigar string because they span splicing junctions were split using GATK (v4.1.3.0) and then base quality score recalibration was performed with Picard. Variant calls were generated with GATK Haplotype Caller with standard settings for variant calling in RNA: minimum-mapping-quality 30, minimum-base-quality 20, dont-use-soft-clipped-bases, standard-call-conf 20.

To identify somatic mutations private to our base-editor treated samples, background filtration was performed using an nCas9 treated sample. Only substitutions on canonical chromosomes were considered. A mutation was determined to be private to the base-editor treated sample if its genomic position had ≥30x coverage in the base-editor treated sample and ≥20x coverage in the nCas9 sample with 99% of reads containing the reference base.

### References:

1. Tsai, S.Q. et al. GUIDE-seq enables genome-wide profiling of off-target cleavage by CRISPR-Cas nucleases. *Nat Biotechnol* **33**, 187-197 (2015).
2. Rees, H.A., Wilson, C., Doman, J.L. & Liu, D.R. Analysis and minimization of cellular RNA editing by DNA adenine base editors. *Sci Adv* **5**, eaax5717 (2019).
3. Li, W. & Godzik, A. Cd-hit: a fast program for clustering and comparing large sets of protein or nucleotide sequences. *Bioinformatics* **22**, 1658-1659 (2006).
4. Bolger, A.M., Lohse, M. & Usadel, B. Trimmomatic: a flexible trimmer for Illumina sequence data. *Bioinformatics* **30**, 2114-2120 (2014).
5. Langmead, B. & Salzberg, S.L. Fast gapped-read alignment with Bowtie 2. *Nat Methods* **9**, 357-359 (2012).
6. Li, H. et al. The Sequence Alignment/Map format and SAMtools. *Bioinformatics* **25**, 2078-2079 (2009).
